# Investigation of Disease-Relevant Lysine Acetylation Sites in α-Synuclein Enabled by Non-canonical Amino Acid Mutagenesis

**DOI:** 10.1101/2025.01.21.634178

**Authors:** Marie Shimogawa, Ming-Hao Li, Grace Shin Hye Park, Jennifer Ramirez, Hudson Lee, Paris R. Watson, Swati Sharma, Zongtao Lin, Chao Peng, Virginia M.-Y. Lee, Benjamin A. Garcia, David W. Christianson, Elizabeth Rhoades, David Eliezer, E. James Petersson

**Affiliations:** Department of Chemistry School of Arts and Sciences, University of Pennsylvania 231 South 34th Street, Philadelphia, PA 19104, USA; Department of Biochemistry Weill Cornell Medicine 1300 York Avenue, New York, NY, 10065, USA; Graduate Group in Biochemistry, Biophysics, and Chemical Biology Perelman School of Medicine, University of Pennsylvania 206 Anatomy-Chemistry Building, 3620 Hamilton Walk, Philadelphia, PA 19104, USA; Department of Biochemistry and Molecular Biophysics Washington University in St Louis 4523 Clayton Ave, St Louis, MO 63130, USA; Department of Neurology David Geffen School of Medicine, University of California - Los Angeles 710 Westwood Plaza, Room C-224, Los Angeles, CA 90095, USA; Department of Pathology and Laboratory Medicine, Center for Neurodegenerative Disease Research, University of Pennsylvania, 3600 Spruce Street, Philadelphia, PA 19104, USA; Department of Biochemistry and Biophysics, Perelman School of Medicine, University of Pennsylvania 421 Curie Boulevard, Philadelphia, PA 19104, USA

## Abstract

Aggregates of α-synuclein (αS) are hallmarks of synucleinopathies, including Parkinson’s Disease (PD) and Multiple System Atrophy (MSA). We have recently shown that αS lysine acetylation in the soluble monomer pool varies between healthy controls, PD, and MSA patients. We used non-canonical amino acid (ncAA) mutagenesis to express and purify all 12 currently known disease-relevant variants of αS and studied their binding to membranes as well as their aggregation propensities. We found that acetylation of lysine 12, 43, and 80 had particularly strong effects. To understand the implications for acetylation of monomeric αS found in healthy cells, we performed NMR experiments to study protein conformation and fluorescence correlation spectroscopy experiments to quantify lipid binding. We also investigated the effects of acetylation at lysine 12, 43, and 80 on fibril seeding in neurons. Collectively, our biochemical and cell biological investigations indicated that acetylation of K_80_ could inhibit aggregation without conferring negative effects on monomer function in healthy cells. Therefore, we studied the structures of fibrils with K_80_ acetylation through cryo-electron microscopy to uncover the structural basis for these effects. Finally, we identified inhibition of HDAC8 as a way of potentially increasing acetylation at K_80_ and other inhibitory sites for therapeutic benefit.

## Introduction

α-Synuclein (αS) is a 14 kDa protein that typically exists at presynaptic terminals in healthy neurons, where its primary function is believed to be in synaptic vesicle trafficking and regulating neurotransmission **(Burré *et al*., 2010; Cabin *et al*., 2002)**. Aggregates of αS commonly characterize several neurodegenerative diseases such as Parkinson’s Disease (PD), Dementia with Lewy Bodies (DLB) and Multiple System Atrophy (MSA), which are referred to as synucleinopathies. Evidence indicates that distinct pathology is caused by αS fibrils formed in different disease environments, or αS “strains.” Aggregation seeding experiments showed that αS strains have distinct abilities to propagate pathology, where αS fibrils from MSA patients are much more potent in seeding aggregation than those from DLB **(Peng *et al*., 2018)**. In addition to this, recent cryo-electron microscopy (cryo-EM) experiments showed that structures of αS fibrils vary between different pathological contexts in PD/DLB **(Yang *et al*., 2022)** and MSA **(Schweighauser *et al*., 2020)**. Despite these findings, the mechanism underlying these differences remains to be understood. It has been suggested that post-translational modifications (PTMs) may contribute to these differences **(Baskakov, 2021)**. Among the PTMs that have been studied on αS thus far are *N*-terminal acetylation, phosphorylation, O-GlcNAcylation, lysine acetylation, lysine ubiquitination, tyrosine nitration and glutamate arginylation **(Baskakov, 2021; Chen *et al*., 2019; Pancoe *et al*., 2022; Schaffert and Carter, 2020)**.

We have recently published a comprehensive study of the relative levels of PTMs in the soluble αS monomer pool between MSA, PD, and DLB patients vs. healthy controls **(Zhang *et al*., 2023)**. While many of the PTMs identified have been previously studied in chemical detail by our laboratory and others,**(Moon *et al*., 2021; Pancoe *et al*., 2022)** lysine acetylation stood out as a PTM that is very common and highly physiologically relevant in other proteins, but had received relatively little attention to date in the context of αS. Given that other αS PTMs have found great significance as biomarkers (e.g. pS_129_ – a hallmark of PD **(Fujiwara *et al*., 2002)**) and drug targets (e.g. kinase inhibitors (**Pagan *et al*., 2019**)), we wished to investigate these acetyl lysine (^Ac^K) sites more thoroughly.

Lysine acetylation is a reversible PTM that can be introduced at specific sites by lysine acetyltransferases (KATs) or non-enzymatically added by reaction with abundant cytosolic acetyl coenzyme A. Lysine deacetylation is catalyzed by lysine deacetylases (KDACs), which include Zn^2+^-dependent histone deacetylases (HDACs) and NAD^+^-dependent sirtuins ***(Ruijter et al., 2003; Sauve et al*., 2006)**. In addition to our comprehensive PTM study in patient samples, there has been some previous evidence for the role of lysine acetylation in synucleinopathies. It has been suggested that activity imbalances between KATs and KDACs on histone or non-histone proteins are pathologically relevant to PD. In fact, activators of some sirtuins and inhibitors of specific KDACs/KATs have shown potential as therapeutics (**Wang *et al*., 2020**). Identified as a substrate of these enzymes, αS was found acetylated on Lys6 and Lys10 in mouse brain. Sirtuin-2 was found to deacetylate those sites and enhance the toxicity of αS **(de Oliveira *et al*., 2017)**. It is notable that in this work, semi-synthetic, acetylated αS was used for the deacetylation assay, however for other experiments glutamine was used to mimic lysine acetylation, which is a common strategy of choice in the field of biochemistry or biophysics, due to easier access to the site-specifically, homogenously modified construct.

Recently, many more disease-relevant lysine acetylation sites have been identified in patient tissue. Eight ^Ac^K sites were identified by Goedert and Scheres in mass spectrometry (MS) studies accompanying a cryo-EM structure of αS fibrils from MSA patients (Lys21/23/32/45/58/60/80/96, Figure 1A) **(Schweighauser *et al*., 2020).** Ten ^Ac^K sites, many overlapping those found by Goedert and Scheres, were identified by MS in our previously noted studies of soluble αS from patients (Lys12/21/23/34/43/45/58/60/96/102, Figure 1A) **(Zhang *et al*., 2023)**. In our accompanying mechanistic studies of the PTMs, authentic constructs of phosphorylated αS were produced through semi-synthesis because phosphorylation occurred at a few key sites with established semi-synthetic routes, but lysine acetylation was investigated only though glutamine mimics due to challenges in systematically investigating a large number of PTM sites where there was less literature to identify key targets **(Zhang *et al*., 2023)**. Thus, there have not been studies of the effect of authentic lysine acetylation in αS at the sites identified from patient tissue. Acetylation is likely to affect monomer binding to membranes (Figure 1B/C), where Lys residues interact with negatively charged phospholipid head groups, and fibril formation, where Lys residues interact with other αS residues or co-factors in structures of recombinant protein fibrils (Figure 1D) or fibrils isolated from MSA patients (Figure 1E).

**Figure 1.**
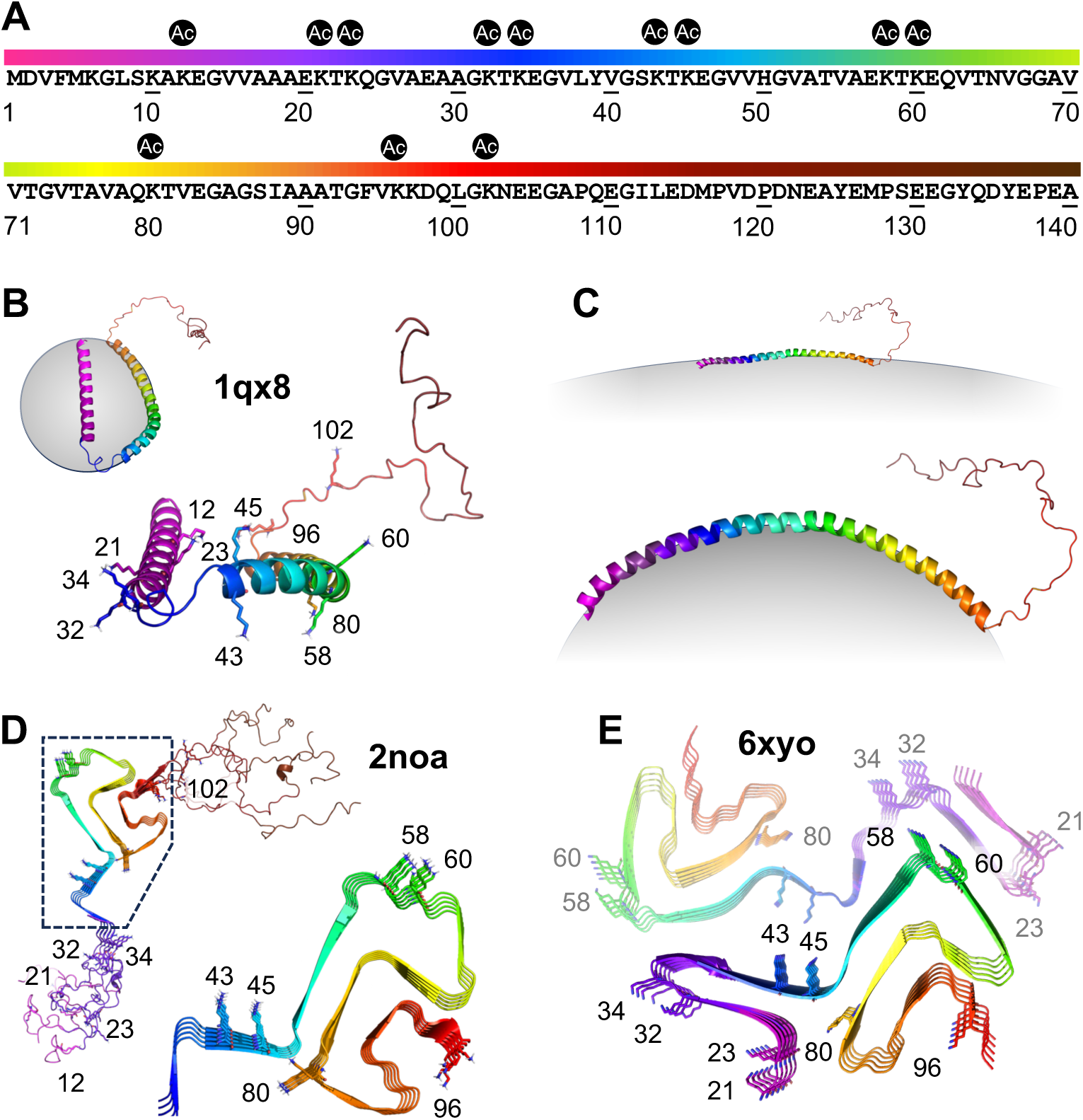
Neurodegeneration-relevant Lys acetylation sites in αS. (A) αS sequence with positions 12, 21, 23, 32, 43, 45, 58, 60, 80, 96 and 102 marked. (B) Solution NMR structure of micelle-bound αS (PDB: 1qx8) **(Ulmer *et al*., 2005)**. (C) Proposed structure of vesicle-bound αS based on electron paramagnetic resonance and fluorescence data, where the first ∼100 residues form an amphipathic membrane-bound helix that matches vesicle curvature **(Jao *et al*., 2004; Middleton and Rhoades, 2010)**. (D) Solid-state NMR structure of recombinant αS fibrils (PDB: 2noa) **(Tuttle *et al*., 2016)**. (E) Cryo-EM structure of MSA patient αS fibrils (PDB: 6xyo) **(Schweighauser *et al*., 2020)**.

In this work, we set out to study lysine acetylation at all 12 disease-relevant sites of αS (Figure 1A). We began by comparing the efficiency of producing acetylated αS through either native chemical ligation (NCL) or non-canonical amino acid mutagenesis (ncAA mutagenesis). We found that ncAA mutagenesis provided comparable yields, and was therefore superior for scanning many acetylation sites due to the ease of generating new constructs. Once the 12 αS ^Ac^K variants were expressed and purified, we studied their binding to membranes as well as their aggregation propensities. We performed NMR, fluorescence correlation spectroscopy (FCS), and transmission electron microscopy (TEM) experiments on acetylated variants that showed perturbed membrane binding or aggregation. NMR and FCS experiments were enabled by our ncAA mutagenesis approach which made it facile to produce isotopically or fluorescently labeled αS. We went on to characterize the seeding ability of select ^Ac^K constructs in neurons, determine a cryo-EM structure of fibrils with a particular ^Ac^K site of interest, and to test HDAC selectivity in deacetylating these sites. The combination of the site-specific incorporation approach and a variety of biological characterization methods provides a systematic understanding of lysine acetylation, identifying a few key ^Ac^K sites as significant for further investigation and potential therapeutic intervention.

## Results and Discussion

### Comparison of ncAA Mutagenesis and NCL

Protein semi-synthesis is a powerful approach to site-specifically incorporate modifications of interest into a protein sequence **(Thompson and Muir, 2020)** and it has been a method of choice for many αS PTM studies **(Moon *et al*., 2021)**, including Lys acetylation **(de Oliveira *et al*., 2017)**. To test this approach to synthesizing acetylated αS, we chose ^Ac^K_80_ as an example, and combined solid-phase peptide synthesis (SPPS),**(Merrifield, 1963)** by which *N*,-acetyllysine is incorporated, with the expression of protein fragments and a three-part NCL sequence using acyl hydrazides **(Zheng *et al*., 2013)** (Figure 2A, additional details in Appendix 1 – figure 1).

**Figure 2.**
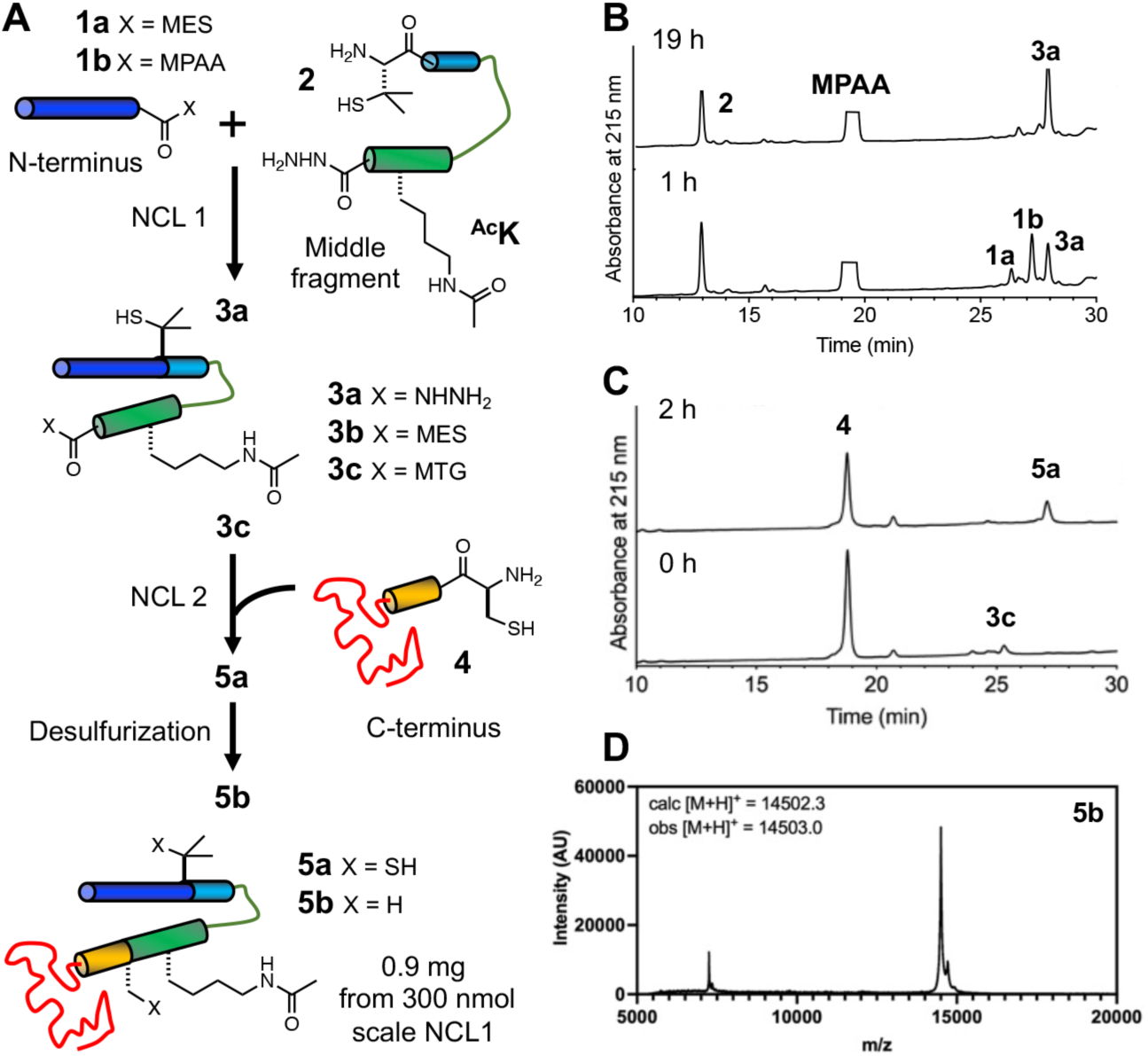
Semi-synthesis of αS-^Ac^K_80_. (A) Acetylation is introduced through peptide synthesis, and the peptide is combined with expressed peptide fragments using NCL. (B) Analytical HPLC trace for the first ligation. **1a**: αS_1-76_-MES, **1b**: αS_1-76_-MPAA, **2:** αS_77-84_-Pen_77_^Ac^K_80_-NHNH_2_, **3a:** αS_1-84_-Pen_77_^Ac^K_80_-NHNH_2_. (C) Analytical HPLC trace for the second ligation. **3b:** αS_1-84__Pen_77_^Ac^K_80_-MES, **3c:** αS_1-84__Pen_77_^Ac^K_80_-MTG, **4:** αS_85-140_-C_85_, **5a:** αS-Pen_77_C_85_^Ac^K_80_. (D) MALDI MS of HPLC-purified αS-^Ac^K_80_ (**5b**).

*N*-terminal thioester fragment αS_1-76_-MES (**1a**) and C-terminal fragment αS_85-140_-C_85_ (**4**) were each recombinantly expressed as a fusion with Mxe GyrA intein. The *N*-terminal thioester was generated by adding excess sodium 2-mercaptoethane sulfonate (MESNa) to cleave the intein by N,S-acyl shift **(Muir *et al*., 1998)** (reported yield 24.1 mg/L **(Pan et al., 2020a)**). Endogenous methionyl aminopeptidase in *E. coli* processes the *N*-terminus of the 85-140 peptide to expose the *N*-terminal cysteine, **(Xiao *et al*., 2010)** which further reacts with aldehydes or ketones *in vivo* to form thiazolidine derivatives **(Liu *et al*., 2016)**. The thiazolidine derivatives were deprotected with methoxyamine to give a free *N*-terminal cysteine (4.40 mg/L, SI Figure S1B). The middle acyl hydrazide peptide αS_77-84_-Pen_77_^Ac^K_80_-NHNH_2_ (**2**, Pen: penicillamine **(Haase *et al*., 2008)**) was synthesized through SPPS (Yield: 12.4 mg, 12 μmol, 48%, Appendix 1 – figure 2A).

αS_1-76_-MES (**1a**) and αS_77-84_-Pen_77_^Ac^K_80_-NHNH_2_ (**2**) were ligated overnight under routine NCL conditions (NCL1) in the presence of 4-mercaptophenylacetic acid (MPAA). (Yield: 1.46 mg, 172 nmol, 57%, Appendix 1 – figure 2B). The purified product (**3a**) was activated by oxidation to form a MES thioester (**3b**) (Yield: 1.29 mg, 126 nmol, 73%, Appendix 1 – figure 2C). The second ligation (NCL2) between αS_1-84_-Pen_77_^Ac^K_80_-MES (**3b**) and αS_85-140_-C_85_ (**4**) to form αS-Pen_77_C_85_^Ac^K_80_ (**5a**) as performed in the presence of methyl thioglycolate to allow for desulfurization without intermediate purification (Figure 2C) **(Huang *et al*., 2016)**. The product, αS-^Ac^K_80_ (**5b**), was obtained in 43% yield (0.90 mg, 62 nmol, Figure 2D). Although we successfully completed this synthesis, it is notable that we encountered solubility issues of the intermediate fragments (**3b**, **5a**) and the product (**5b**), after lyophilization.

Experiencing difficulties in sample handling and considering the inefficiency of applying NCL to scan 12 lysine acetylation sites distributed throughout the protein, we then sought to access site-specifically acetylated αS through ncAA mutagenesis (Figure 3A). We recombinantly expressed αS (plasmid sequence in Appendix 1 – figure 3) with lysine acetylation at site 80 in *E. coli* through amber codon suppression. We used a previously reported pair of aminoacyl tRNA synthetase (chAcK3RS with IPYE mutations) and cognate tRNA to incorporate *N*^ε^-acetyllysine at a position dictated by an amber stop (TAG) codon **(Bryson *et al*., 2017)**. In addition to 10 mM *N*^ε^-acetyllysine, 50 mM nicotinamide, an inhibitor to endogenous deacetylases, was added to the media before inducing αS expression. The protein was expressed as an intein fusion as reported before for easy removal of truncated protein through affinity purification **(Batjargal et al., 2015)**. After traceless intein cleavage with 2-mercaptoethanol, the ^Ac^K-containing protein was purified by reverse phase high performance liquid chromatography (RP-HPLC) and exchanged into appropriate buffers for biophysical assays (Figure 3B).

**Figure 3.**
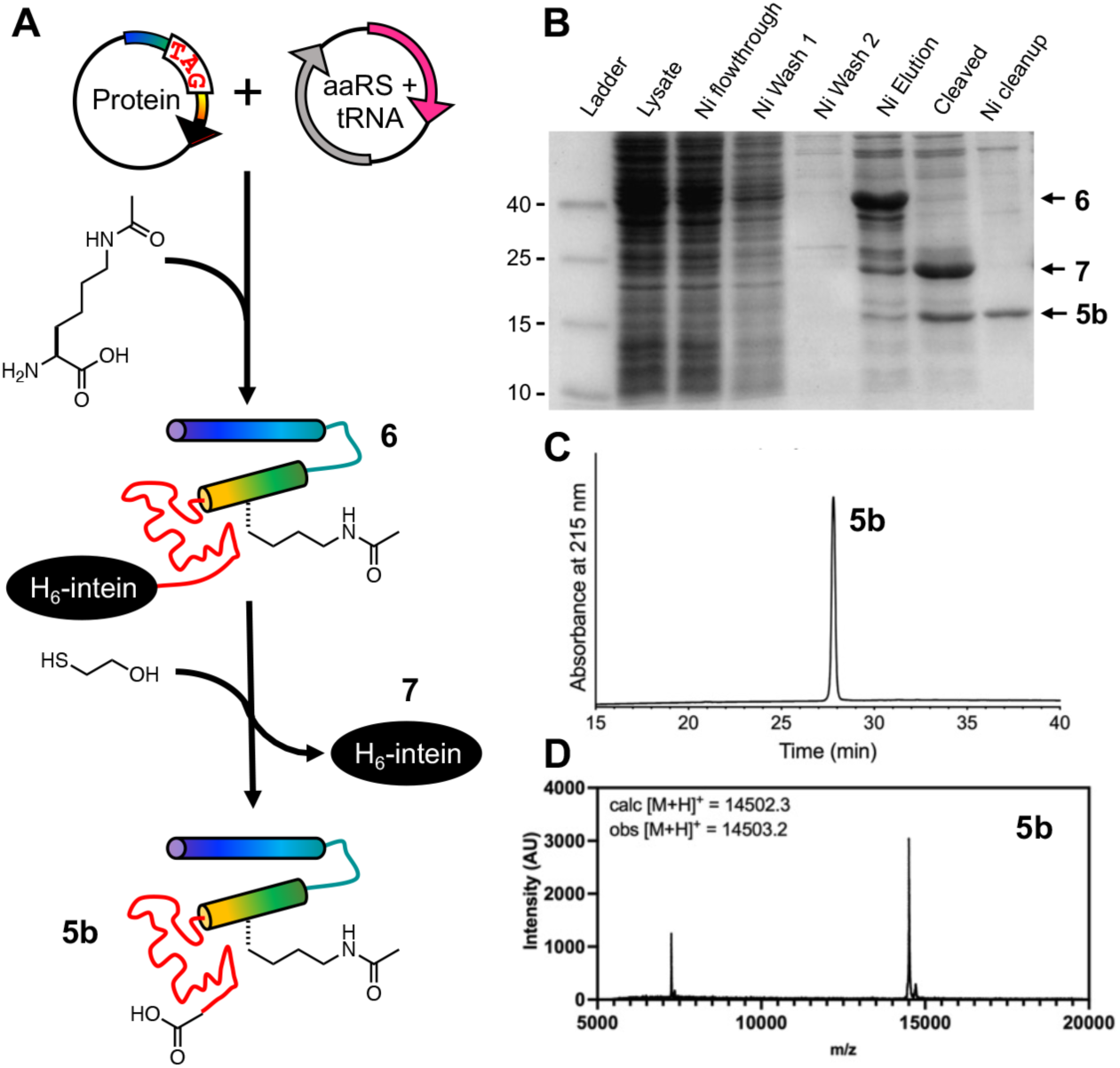
Expression of αS-^Ac^K_80_ through ncAA mutagenesis. (A) An orthogonal aminoacyl tRNA synthetase (aaRS)/tRNA pair site-specifically incorporates acetyllysine in recombinant αS. Intein tagging at the C-terminus allows for traceless purification of the full-length product. (B) SDS-PAGE gel (Coomassie stain) showing Ni-affinity purification of recombinant αS-^Ac^K_80_. Purified αS-^Ac^K_80_ (**5b**) characterized with (C) analytical HPLC and (D) MALDI MS.

We obtained 0.65 mg of pure αS-^Ac^K_80_ per L of bacterial culture (Figure 3C, 3D), a yield comparable to that obtained by NCL, but because no lyophilization or handling of ligation intermediates is required, we did not encounter the solubility problems observed in the NCL process. Therefore, we deemed the ncAA mutagenesis approach at least comparable to NCL for producing a specific construct. Since we wished to study 12 sites distributed throughout the αS sequence, ncAA mutagenesis was also advantageous because we avoided having to generate constructs for several different ligation sites and could simply perform site-directed mutagenesis to insert TAG codons for each new αS-^Ac^K_n_ variant.

Bolstered by our success with ^Ac^K_80_, we generated TAG mutants at sites 12, 21, 23, 32, 34, 43, 45, 58, 60, 96, or 102. We expressed and purified these proteins, observing successful ncAA mutagenesis at each site (Appendix 1 – figures 4-14), however, the yield varied significantly between different sites (0.11-1.5 mg/L of culture). This is an interesting result in light of the large number of sites that were tested in the same protein and the fact that αS is an intrinsically disordered protein, so protein folding should not affect incorporation. Examination of the local RNA sequence context of the amber (TAG) codon did not explain the varied suppression efficiency, based either on previously identified sequence impacts **(Pott *et al*., 2014)** or by comparing the sites within αS (Appendix 1 – figure 15). Given the pseudo-repeat nature of the αS sequence, many sites feature similar sequences, and a comparison of 21 and 58 is particularly striking with a 10-fold difference in expression levels despite near identity in the flanking sequences. While these observations are notable for users of ncAA technology, in the context of this study, our approach allowed us to acquire sufficient amounts of the 12 different authentically modified αS constructs for biophysical experiments.

Thus, in spite of low expression yields for some sites, ncAA mutagenesis was a preferred method for this work, due to the better efficiency in scanning 12 different modification sites and the ease of handling aggregation-prone protein fragments. The expression-based strategy also allows for low-cost access to isotopically labeled, PTM-modified αS constructs, as we have demonstrated previously **(Pan *et al*., 2021)**.

### Effects on αS Helicity on Micelles and Aggregation

αS is known to bind to lipid surfaces and form helical structures, part of its physiological role in modulating neurotransmitter vesicle trafficking **(Ramirez *et al*., 2023)**. More specifically, on micelles, an NMR structure showed that micelle-bound αS forms a broken helix, where two helical strands are connected with a loop region (Figure 1B, PDB: 1xq8) **(Ulmer *et al*., 2005)**. A helical wheel model, created based on this structure, shows that lysine residues are aligned on the membrane surface, and that they are likely involved in enhancing binding by interactions with negatively charged lipid head groups **(Meade *et al*., 2019)**.

With each acetylated αS variant, we first examined the effects of Lys acetylation on the secondary structure of αS in the presence of micelles by wavelength scan circular dichroism (CD) spectroscopy. Each acetylated αS was compared to unmodified wild type (WT) αS in phosphate-based buffer, pH 7.4, with a large excess of sodium dodecyl sulfate (SDS). We normalized the molar ellipticity at 222 nm of each acetylated construct to that of WT to compare the effect on helicity at each site. We found that significant reduction of helicity was caused only by acetylation at site 43 and that acetylation at other sites had only minor effects on helicity (Figure 4, Appendix 1 – figure 16). This study implies that only this site could potentially perturb αS function in neurotransmitter trafficking. However, since SDS micelles are crude mimics of neurotransmitter vesicles, additional investigations with vesicles composed of biologically relevant lipids (see below) and cell-based studies of neurotransmission should be used to purse these findings further.

**Figure 4.**
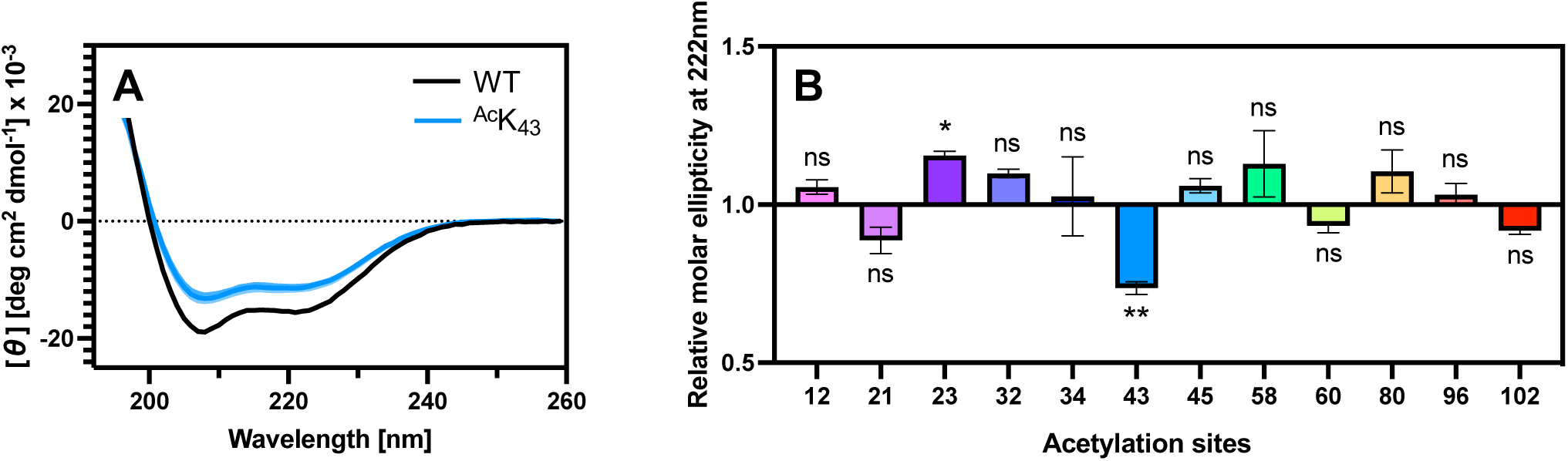
Effects of lysine acetylation on micelle-bound αS. (A) CD spectra for WT and ^Ac^K_43_ αS. spectra for other constructs are shown in Appendix 1 – figure 16. (B) Molar ellipticity at 222 nm was normalized to WT value to quantify helicity on SDS micelles. Mean with SD, R=3

We then investigated whether lysine acetylation at different sites has impacts on αS in pathological contexts. To do this, we first performed *in vitro* aggregation experiments and assessed site-specific effects. A plate-based approach was taken to efficiently perform the assay, and each aggregation reaction was seeded by mixing with αS WT pre-formed fibrils (PFFs) that constituted 10% of the total monomer concentration. This type of seeded aggregation assay primarily reports on the elongation phase of aggregation rather than the nucleation phase (**Pancoe *et al*., 2022**). The monomer samples were prepared by mixing αS WT with acetylated αS at either 10% or 25% of the total monomer concentration. These concentrations were chosen because our quantitative studies of other PTMs in patient samples indicated that most were present in this range, rather than stoichiometrically (**Zhang *et al*., 2023; Zhao *et al*., 2022)** (see additional discussion in Conclusions). Aggregation was carried out at 37 ℃ with shaking and kinetics and thermodynamics (final fibril amounts) were monitored.

To examine the effects on aggregation kinetics, we took advantage of the change in fluorescence of the amyloid binding dye, thioflavin T (ThT), during aggregation to monitor the process *in situ*. We found that the effects differ between different modification sites (Figure 5, Appendix 1 – figures 17-18). For Lys acetylation at 12, 23, 43, 80 and 102 we observed differential slowing effects – the effects were particularly significant at sites 12, 43 and 80, and the effects at 12, 23 and 43 were dose dependent. While we observed acceleration of aggregation for site 32 both at 10% and 25%, the effects were similar between the different dosages.

**Figure 5.**
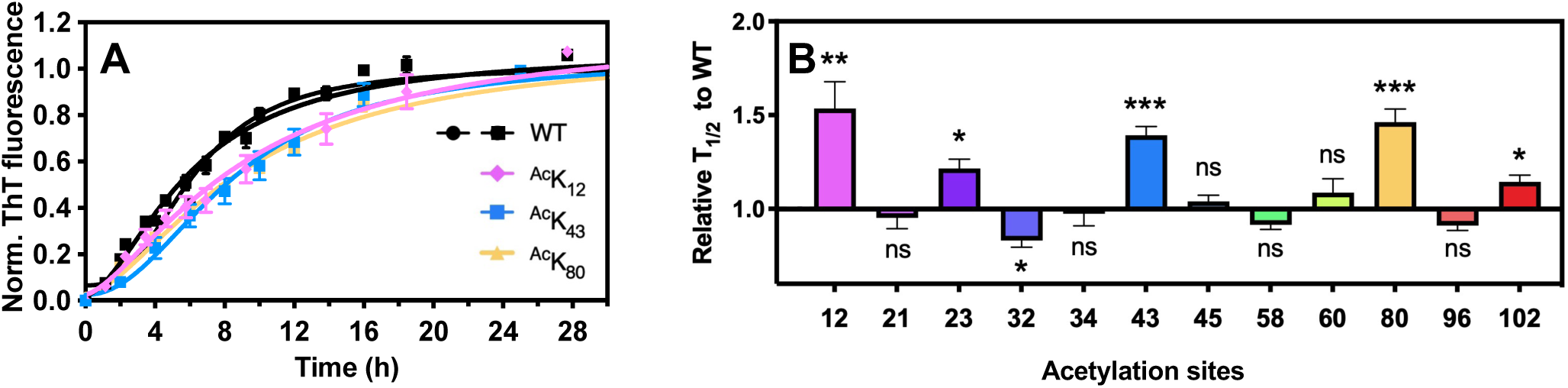
Effects of lysine acetylation on *in vitro* aggregation. (A) Aggregation kinetics were monitored by fluorescence intensity change of ThT. Two WT traces are shown, one trial conducted with ^Ac^K_12_ (squares) and one trial conducted with ^Ac^K_43_ and ^Ac^K_80_ (circles). ^Ac^K_12_ data were scaled by WT trials for clarity. Original data are shown in Appendix 1 – figure 18. (B) Time to reach 50% fibrilization (T_1/2_) for each condition was normalized to that of WT. Seeded aggregation was performed with αS monomers where acetylated αS was mixed with αS WT at 25%:75% ratio. SEM, R=6

To confirm that these effects were not the result of reduced monomer incorporation, we isolated fibrils at the endpoint of 10% or 25% aggregations and SDS-PAGE gels were run and stained with the Coomassie Brilliant Blue dye to quantify total monomer incorporation into the fibrils. We found that there were no consistent reductions in monomer incorporation, and in fact there were some moderate apparent enhancements of incorporation. However, these were generally not consistent between the 10% and 25% aggregation experiments, except in the case of ^Ac^K_34_ (Appendix 1 – figures 19-20). Taking all of the aggregation kinetics and monomer incorporation data into account, we chose to investigate the kinetically perturbed sites 12, 43, and 80 further, since the cellular process will be unlikely to reach equilibrium and our previous study had shown that the Gln mimic mutation at position 34 did not alter aggregation in cells **(Zhang *et al*., 2023)**.

### Fibril Seeding in Neurons

To investigate the impact of Lys acetylation on aggregation in more physiologically relevant contexts – in cultured neurons – we followed the approach that we have done previously with arginylated αS.**(Pan *et al*., 2022; Zhao *et al*., 2022**) We prepared PFFs with the following compositions: αS WT or αS WT mixed with 25% acetylated αS, ^Ac^K_12_, ^Ac^K_43_ or ^Ac^K_80_. Mouse primary hippocampal neurons were grown for 8 days on a coated plate, to which 50 ng/µL PFFs were added, following established protocols.(**Haney *et al*., 2016; Luk *et al*., 2009; Marotta *et al*., 2021)** After 2 weeks, intracellular αS aggregates were quantified by staining with an antibody that recognizes phosphoserine 129 (pS_129_), a commonly used pathological marker (Figure 6, Appendix 1 – figure 21). Compared to the WT PFFs, all the acetylated PFFs tested resulted in significantly reduced aggregation seeding: the pS_129_ signal (AU ± SEM; arbitrary units, standard error of the mean) of PFF-seeded αS aggregates was 2347 ± 107.1 (WT), 1730 ± 83.67 (^Ac^K_12_), 1854 ± 70.79 (^Ac^K_43_), 1698 ± 54.41 (^Ac^K_80_) with respect to DAPI. Notably, acetylation at these sites also slowed seeded aggregation in the *in vitro* fibrilization experiment, but did not reduce aggregates quantified at the endpoint. This supports the idea that aggregation in a cellular context is unlikely to reach saturation. However, it is important to note that the experiments in neurons featured acetylation on the fibril seeds, whereas the *in vitro* experiments featured acetylation on the monomer pool, and are thus reflective of different aspects of the aggregation process.

**Figure 6.**
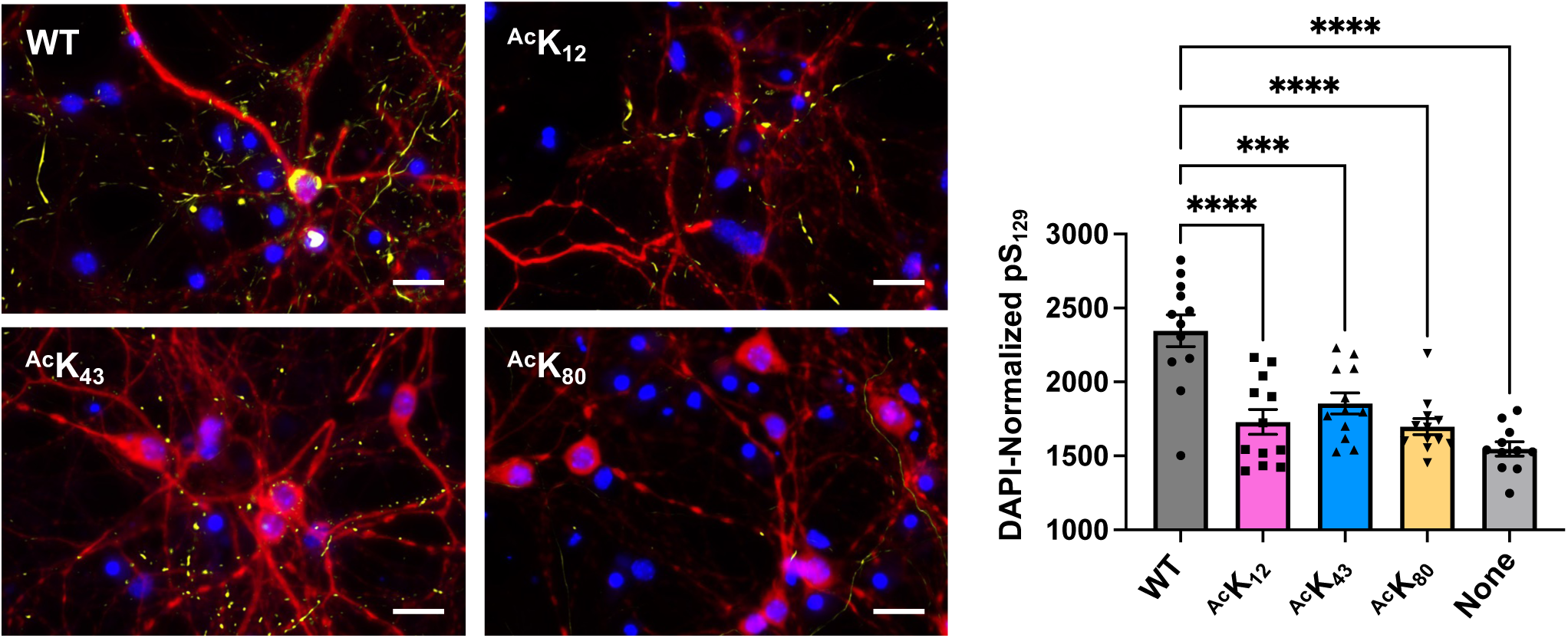
Effects on aggregation seeding in primary neuron cells. Left: representative images of neuron cultures treated with unmodified or 25% acetylated αS PFFs, stained with an anti-pS129 antibody (yellow), DAPI (blue), and an anti-MAP2 antbody (red). Scale bar = 25 µm. Larger fields of view shown in Appendix 1 – figure 21. Right: quantification of DAPI-normalized anti-pS_129_ area of intracellular aggregates seeded by different αS PFFs. Mean with SE, R= 11-12. *** = 0.001 < p-value < 0.0001; **** = 0.00001 < p-value < 0.0001

### Structural Characterization of αS Monomers

Having demonstrated that acetylation at K_12_, K_43_ or K_80_ significantly reduced αS aggregation *in vitro* and in cells, we wished to gain information on the structural impact of acetylation at these sites. First, to give insights into the effects on monomer conformation, we acquired proton-nitrogen correlation spectra (^1^H,^15^N – HSQC) for ^Ac^K_12_, ^Ac^K_43,_ or ^Ac^K_80_, an experiment that is facile with ncAA mutagenesis, but challenging to perform via NCL due to the high cost of isotopically-labeled amino acids for SPPS. To access ^15^N-labeled αS, we expressed the acetylated αS and αS WT in M9 minimal media containing ^15^N-labeled ammonium chloride (Appendix 1 – figures 22-24). This afforded comparable protein yields to expressions in LB media. It is notable, however, that sub-stoichiometric isotopic labeling was observed at some Lys sites, depending on batches of expression, which could be due to deacetylation in *E. coli* cells followed by incorporation at Lys codons. Overlaying the HSQC spectra for αS-WT and αS- ^Ac^K_12_, ^Ac^K_43_ or ^Ac^K_80_, peak shifts were observed only in signals from surrounding residues, suggesting that the structural change was local and there is no major impact of lysine acetylation on monomer structure (Appendix 1 – figures 25-26).

### Biophysical Characterization of Lipid Binding

We next wished to learn the effects of Lys acetylation at K_12_, K_43_ or K_80_ on the native function of αS by investigating its lipid binding mode. To quantify conformational changes of αS upon vesicle binding, we acquired ^1^H,^15^N – HSQC spectra for WT, ^Ac^K_12_, ^Ac^K_43_ or ^Ac^K_80_ in the presence of small unilamellar vesicles (SUVs) that are composed of 60:25:15 1,2-dioleoyl-sn-glycero-3-phosphocholine/1,2-dioleoyl-sn-glycero-3-phosphatidylethanolamine/1,2-dioleoyl-sn-glycero-3-phospho-L-serine (DOPC/DOPE/DOPS). The NMR NH chemical shifts were similar for all constructs and consistent with spectra previously reported for WT αS (**Eliezer *et al*., 2001**). There was no notable backbone chemical shift perturbation at the surrounding residues of each acetylation site (Appendix 1 – figure 27).

In the presence of vesicles, a reduction of intensity for residues 1-100 was observed for all the constructs, which is caused by binding of this portion of αS to the slowly tumbling lipid vesicles and is again consistent with previous observations **(Bodner *et al*., 2009; Bussell and Eliezer, 2004**). αS- ^Ac^K_80_ had a similar intensity change to WT (∼40%, Figure 7A, Appendix 1 – figure 28), whereas αS-^Ac^K_43_ had a smaller intensity change, suggesting weaker vesicle binding (∼20%, Figure 7A, Appendix 1 – figure 28). This is consistent with the acetylation effects observed with SDS micelles (Figure 4). αS-^Ac^K_12_ had an intermediate intensity reduction (∼30%, Figure 7A, Appendix 1 – figure 28), a more significant effect of K_12_ acetylation on vesicle binding than what was observed with SDS micelles (Figure 4). It is possible that this is due to the differences in curvature and headgroup between the micelles and the vesicles, which is known to result in different αS binding modes (**Cholak *et al*., 2020; Georgieva *et al*., 2010; Middleton and Rhoades, 2010; Rhoades et al., 2006; Trexler and Rhoades, 2009**). It is also possible that this is due to the increased sensitivity of NMR to subtle differences in binding.

**Figure 7.**
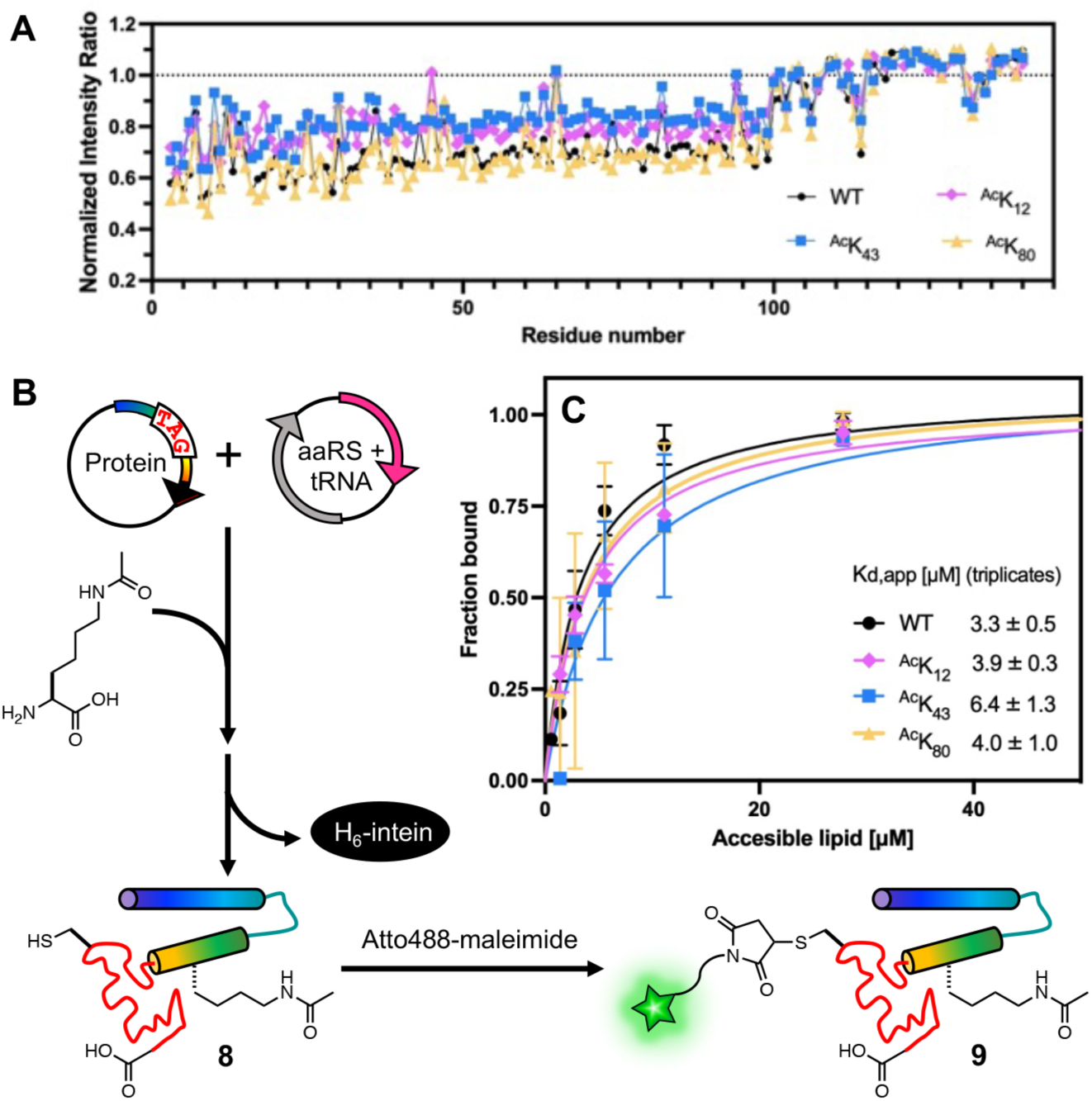
Effects of lysine acetylation on vesicle binding affinity. (A) NMR intensity ratio for each residue calculated from ^1^H-^15^N HSQC spectra collected with ^15^N-labeled αS variants in the presence or absence of SUVs, normalized by the average ratio for residues 101-140. (B) αS with a TAG codon at the acetylation site of interest and a Cys mutation at a labeling site (**8**) was co-expressed with an aaRS/tRNA plasmid for acetyllysine incorporation. After intein cleavage, labeling with an Atto488 dye was performed through Cys-maleimide chemistry to give an acetylated, labeled protein (**9**) for FCS. (C) Vesicle binding affinity determined by fluorescent correlation spectroscopy measurements. For each construct, measurements were performed on three separate days. Mean with SD, R=3

While NMR is a very valuable technique for characterizing vesicle binding with a non-perturbing label, to measure affinity, we turned to FCS, a well-established method for rigorously determining vesicle apparent dissociation constants (K_d,app_) **(Rhoades *et al*., 2006)**. To enable this experiment, we expressed the acetylated αS constructs at site 12, 43, or 80 or a non-acetylated construct (“WT”), bearing a Cys mutation at site 114 (**8**, αS–^Ac^K_12_C_114_, αS–^Ac^K_43_C_114_, αS–^Ac^K_80_C_114,_ and αS–C_114_) to allow for fluorescent labeling (Figure 7B, Appendix 1 – figures 29-32). The fluorophore Atto488-maleimide was reacted with purified Cys mutants overnight at 4 ℃ or for a few hours at room temperature to yield labeled constructs (**9**, αS–^Ac^K_12_C^Atto488^_114_, αS–^Ac^K_43_C^Atto488^_114_, αS–^Ac^K_80_C ^Atto488^_114,_ and αS–C ^Atto488^_114_), and the conversion was almost quantitative (Appendix 1 – figures 29-32). We prepared synthetic lipid vesicles containing 50:50 1-palmitoyl-2-oleoyl-sn-glycero-3-phospho-L-serine/1-palmitoyl-2-oleoyl-glycero-3-phosphocholine (POPS/POPC). The diffusion times of free αS and of the vesicles were obtained first, and in assessing the αS-vesicle binding, we added the same quantity of αS to varied concentrations of vesicles and then determined the protein fractions bound by fitting a two-component autocorrelation function. The fraction bound values at each vesicle concentration were used to fit a binding curve for each αS construct. We found that acetylation at site 43 leads to two-fold weaker binding and acetylation at site 12 or 80 did not significantly affect binding (Figure 7C, Appendix 1 – figures 33-34; K_d,app_^WT^ = 3.2 ± 0.5 μM, K ^AcK12^ = 3.9 ± 0.3 μM, K_d,app_^Ac^K_43_ = 6.3 ± 1.3 μM, K_d,app_^AcK80^ = 4.0 ± 1.0 μM).

Slightly reduced binding due to acetylation at site 43 correlates with the reduced helicity we observed in the CD wavelength scan and the differences in NMR peak intensities in the presence of vesicles. The NMR experiments showed that ^Ac^K_43_ reduced vesicle binding more significantly than ^Ac^K_12_ (moderate) or ^Ac^K_80_ (little to none). The FCS experiments supported this, showing that ^Ac^K_43_ led to weaker binding than ^Ac^K_12_ or ^Ac^K_80_, which were similar to WT. Previous NMR experiments suggested that the *N*-terminal helix of αS (residues 6-25) drives association with lipid membranes and the 26-97 region modulates the affinity, depending on lipid composition **(Fusco *et al*., 2014)**. The different effect between ^Ac^K_43_ and ^Ac^K_12_ or ^Ac^K_80_ suggests that K_43_ is more important in modulating the binding affinity.

Taken together, our results show that among all the disease-relevant acetylation sites, ^Ac^K_12_, ^Ac^K_43_, and ^Ac^K_80_ each inhibit aggregation, but that ^Ac^K_43_ also inhibits membrane binding (as does ^Ac^K_12_, to a lesser degree). Thus, in the case of ^Ac^K_43_, the potential benefits of reduced amyloidogenicity may be offset by compromising function in neurotransmitter release.

### Structural Characterization of αS Fibrils

To get preliminary insights into fibril structure effects, we performed TEM imaging on fibrils formed from acetylated αS (αS-^Ac^K_12_, ^Ac^K_43_ or ^Ac^K_80_), mixed with αS WT, at 25% of the total monomer concentration (Appendix 1 – figure 35). Interestingly, we observed mixed morphology for PFFs prepared with αS-^Ac^K_12_, with some very narrow fibrils. Both 25% ^Ac^K_12_ and 25% ^Ac^K_43_ PFFs have minimal helical twist, making them difficult to characterize by cryo-EM. On the other hand, for PFFs prepared with 25% αS-^Ac^K_80_, we observed a slightly more twisted fibril morphology, so we attempted to solve a structure using helical reconstruction cryo-EM methods. We were able to solve a structure to 2.88 Å resolution, and comparison to previously published αS WT fibril structures (Appendix 1 – figures 36, 43, table 1) shows that the backbone fold is similar to the two-stranded polymorph typified by PDB ID 6a6b (Figure 8, inset, 6a6b), (**Li *et al*., 2018b**) similar to PDB IDs 6cu7 (**Li *et al*., 2018a**) and 6h6b (**Guerrero-Ferreira *et al*., 2018**), which all have the “Greek key” protein fold first reported in solid state NMR studies of single stranded fibrils under PDB ID 2n0a **(Tuttle *et al*., 2016)**. However, the density clearly shows that K_80_ is not acetylated in this structure, implying that the WT fibrils are forming this 6a6b-type fibril, while the ^Ac^K_80_ fibrils are forming a separate, minority population that we are not able to resolve.

**Figure 8.**
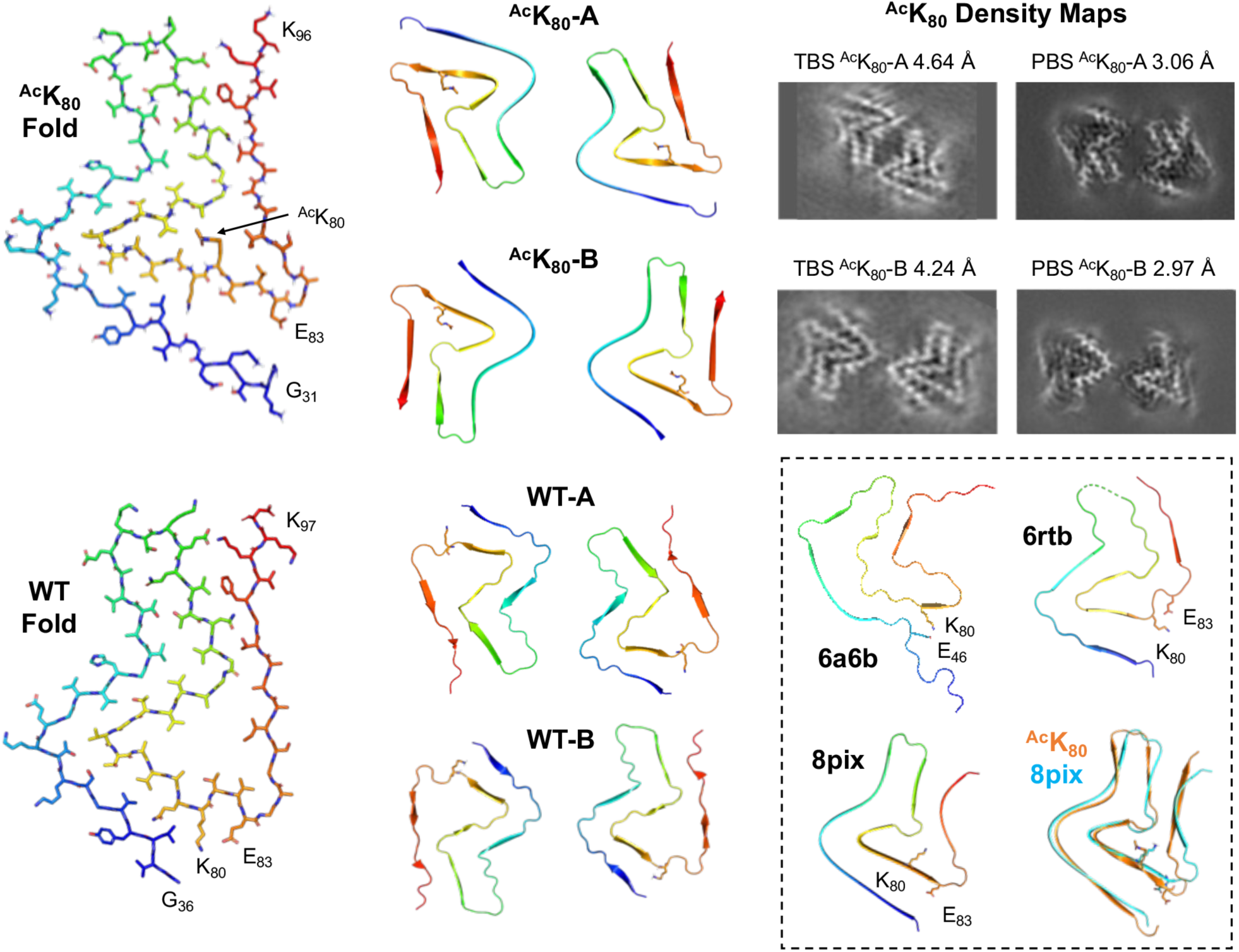
Structural impact of K80 acetylation on fibril morphology. ^Ac^K_80_ Fold and WT Fold show the fold of a single αS molecule in the fibrils, viewed down the fibril axis (from ^Ac^K_80_-A PBS and WT-A TBS structures). ^Ac^K_80_-A and ^Ac^K_80_-B show the two fibril polymorphs, with similar protein folds, but different strand-strand packing (from PBS structures). WT-A and WT-B show the two fibril polymorphs, with similar protein folds, but different strand-strand packing (from TBS structures). ^Ac^K_80_-A Density Maps show that the same fibril polymorphs were obtained for fibrils made in TBS and PBS. Inset: The interactions of K80 are shown in three previously αS fibril polymorphs designated by their PDB IDs.(**Frey *et al*., 2024; Guerrero-Ferreira *et al*., 2018; Li *et al*., 2018b**) The overlay shows the similarity of the ^Ac^K_80_ fold to the 8pix fold.

**Table 1.** Cryo-EM particle numbers for fibril samples.

|  | 25% <sup>Ac</sup>K<sub>80</sub> | WT TBS | WT PBS | <sup>Ac</sup>K<sub>80</sub> TBS | <sup>Ac</sup>K<sub>80</sub> PBS |
| --- | --- | --- | --- | --- | --- |
| Template-Free Filament Tracing Particles | 4,942,459 | 650,880 | 2,031,379 | 871,724 | 6,247,663 |
| Extracted Particles | 3,364,884 | 1,163,693 | 269,189 | 332,878 | 4,158,103 |
| Retained Particles | 228,881 | 764,292 |  | 199,843 | 705,709 |
| Re-extracted Particles | 338,172 | 2,341,469 |  | 834,159 | 893,660 |
| Initial Model Particles | 338,081 | 2,341,469 |  | 834,159 | 893,660 |
| 3D Classification Particles | 296,473 |  |  |  |  |
| 3D Classification Particles A |  | 696,830 |  | 48,610 | 102,778 |
| 3D Classification Particles B |  | 1,317,407 |  | 312,253 | 200,042 |

In order to more clearly observe the structural impact of K_80_ acetylation, we prepared fibrils with 100% ^Ac^K_80_ αS in Tris-buffered saline (TBS) for cryo-EM studies. For these fibrils, we were able to solve structures of two different polymorphs, both composed of two strands (Figure 8, ^Ac^K_80_-A and ^Ac^K_80_-B, Appendix 1 – figures 37, 39, 40, 43, table 2). The protein fold is essentially the same in both polymorphs, but they differ in strand-strand packing. Since it is well-documented that differences in buffer composition and aggregation methods can lead to differences in fibril morphology, we also prepared ^Ac^K_80_ αS fibrils in phosphate-buffered saline (PBS), the same conditions used in our aggregation kinetics studies. Gratifyingly, the ^Ac^K_80_ αS fibrils prepared in PBS exhibited the same two polymorphs seen for TBS ^Ac^K_80_ αS fibrils, with identical protein folds and strand-strand packings (Figure 8, Appendix 1 – figure 41). We were also able to solve cryo-EM structures of WT αS fibrils generated under the same conditions in TBS. We observed two WT polymorphs (Figure 8, WT-A and WT-B, Appendix 1 – figures 37, 38, 40, 43, table 2) which exhibited similar folds and strand-strand packings to the ^Ac^K_80_ polymorphs, but with a notable change in morphology around K_80_. Acetylation of K_80_ disrupts a salt-bridge interaction with E_83_ that can be clearly seen in the WT-B polymorph (Figure 8, WT Fold) and neutralizes the sidechain charge, allowing it to pack in a hydrophobic pocket formed by A_69_ and V_71_ (Figure 8, ^Ac^K_80_ Fold). This leads to a twist of the backbone in the T_75_-A_90_ segment, generating a modest change in the protein fold (local RMSD ∼5 Å, Appendix 1 – figure 42). Given that the ^Ac^K_80_ protein fold is fairly similar to the WT protein fold, it is not surprising that they exhibit similar strand-strand packings and that K_80_ acetylation has a moderate impact on aggregation rates. We can use these structures to consider ^Ac^K_80_ effects in the context of other structural studies of αS fibrils.

Our WT-A and WT-B structures resemble those first reported under PDB IDs 6rtb and 6rto (Figure 8, inset, 6rtb; Appendix 1 – figure 44) **(Guerrero-Ferreira *et al*., 2019**). These polymorphs have been observed by several investigators for WT αS fibrils formed at near-neutral pH, along with the commonly observed 6a6b/2n0a form (Figure 8, inset, 6a6b) **(Li *et al*., 2018b).** Our ^Ac^K_80_ αS fibril structures, ^Ac^K_80_-A and ^Ac^K_80_-B, resemble those recently reported for WT αS fibrils formed at pH ≤6.5 under PDB IDs 8pix and 8pic (Figure 8, inset, 8pix) (**Frey *et al*., 2024**). In the two fibril polymorphs commonly populated at pH 7, K_80_ makes key stabilizing salt bridge interactions. For the 6a6b polymorph, K_80_ makes a salt bridge with E_46_; for the 6rtb polymorph, it makes a salt bridge with E_83_. Disruption of these salt bridges by acetylation would destabilize either fold, favoring the ^Ac^K_80_ fold that we observe. The fact that a similar fold (8pix) has been seen at lower pHs can be rationalized by assuming that acidification leads to protonation of the E_46_ or E_83_ sidechains, weakening their interactions with K_80_ just as acetylation does to drive a change fibril polymorph. While pH 6.5 is significantly above the pK_a_ of a typical glutamate sidechain, it is possible that the pK_a_s are perturbed in the local environment of the fibril, and full deprotonation would not be required to destabilize interactions with K_80_. Thus, one can rationalize our observation of a polymorph like 8pix for our ^Ac^K_80_ at pH 7, when it had only been previously observed at lower pHs. It is further notable that the ^Ac^K_80_ fold is robust to changes in buffer (PBS vs. TBS). The fibril polymorphs that we observe are clearly different than those obtained from MSA patient samples (Figure 1E), and this is likely a consequence of the combination of additional PTMs and non-proteinaceous cofactors present beyond the single K_80_ acetylation in our fibrils. A comprehensive study of the interplay of such PTMs will be necessary to understand how they drive fibril polymorphism, and studies of single PTMs like ours are just a starting point toward such an investigation. However, our study does imply that increasing acetylation of certain key Lys residues can drive fibril formation toward conformations that are not compatible with PD/DLB or MSA polymorphs which would disrupt propagation (Appendix 1 – figure 45). Similar strategies have been proposed around increasing levels of other PTMs.

It is worth comparing the effects that we observe for ^Ac^K_80_ acetylation to the structural and biophysical effects observed with other αS PTMs which have been structurally characterized, such as Y_39_ phosphorylation (pY_39_) or S_87_ phosphorylation (pS_87_) and *N*-Acetyl glucosamine glycosylation (gS_87_). For pY_39_ αS, a 4-fold increase in aggregation rate has been observed for αS 100% phosphorylation, with nuanced effects at lower phosphorylation percentages. We have shown that pY_39_ leads to only modest changes in monomer conformation, based on NMR and single molecule FRET studies **(Pan *et al*., 2021; Pan *et al*., 2020b**). In contrast, Zhao *et al*’s cryo-EM structure shows a dramatic rearrangement of the fibril polymorph **(Zhao *et al*., 2020)**. This indicates that the effect of Y_39_ phosphorylation is primarily at the fibril level, similar to our findings here for ^Ac^K_80_ acetylation. Both pS_87_ and gS_87_ modifications inhibit aggregation much more drastically than pY_39_. Two different structures of gS_87_ αS fibrils have been reported, both showing a significant deviation from reported WT αS polymorphs (**Balana *et al*., 2024; Hu *et al*., 2024**). Despite their differences, the structures both provide a clear rationale for the effects of S_87_ glycosylation. Intriguingly, Hu *et al*. highlight the way in which gS_87_ changes the structure of the 80-89 region of αS to disrupt the E_46_-K_80_, destabilizing the 6a6b fold (**Hu *et al*., 2024**). The pS_87_ structure is different from either gS_87_ structure, and although the residue cannot be observed in the structure, it again demonstrates that a PTM can dramatically alter fibril morphology. Thus, like ^Ac^K_80_ studied here, these PTMs seem to primarily exert their influence on αS aggregation through changes in fibril structure, which is sensible given the disordered nature of the αS monomer.

### Deacetylase Site Specificity

Since acetylation of K_12,_ K_43_ and K_80_ can reduce αS aggregation, and at least ^Ac^K_80_ forms fibrils that are structurally incompatible with PD/DLB and MSA fibril polymorphs, we considered the anti-aggregation potential of increasing acetylation at key Lys residues by inhibiting a KDAC. Doing so would require that the KDAC had some specificity for these residues. Using previously published methods, **(Decroos *et al*., 2014; Dowling *et al*., 2008; Osko *et al*., 2021**) we expressed and purified recombinant human HDAC8, a Zn-dependent HDAC known to act on non-histone proteins, including cytosolic targets like tubulin in HeLa cells (**Vanaja *et al*., 2018**). We treated samples of each of the acetylated αS variants with HDAC8 and monitored deacetylation through a matric assisted laser desorption ionization (MALDI) MS assay using ^15^N-labeled αS as a standard. After 24 h, all of the constructs showed significant levels of deacetylation, but ^Ac^K_34_, ^Ac^K_43_, ^Ac^K_45_, and ^Ac^K_80_ retained an average of 44% acetylation, 3-fold greater levels than all other sites (Figure 9). These preliminary results indicate that inhibition of HDAC8 could increase acetylation levels of αS at specific lysine residues which we have shown to retard fibril elongation *in vitro* and in PFF-seededing neurons.

**Figure 9.**
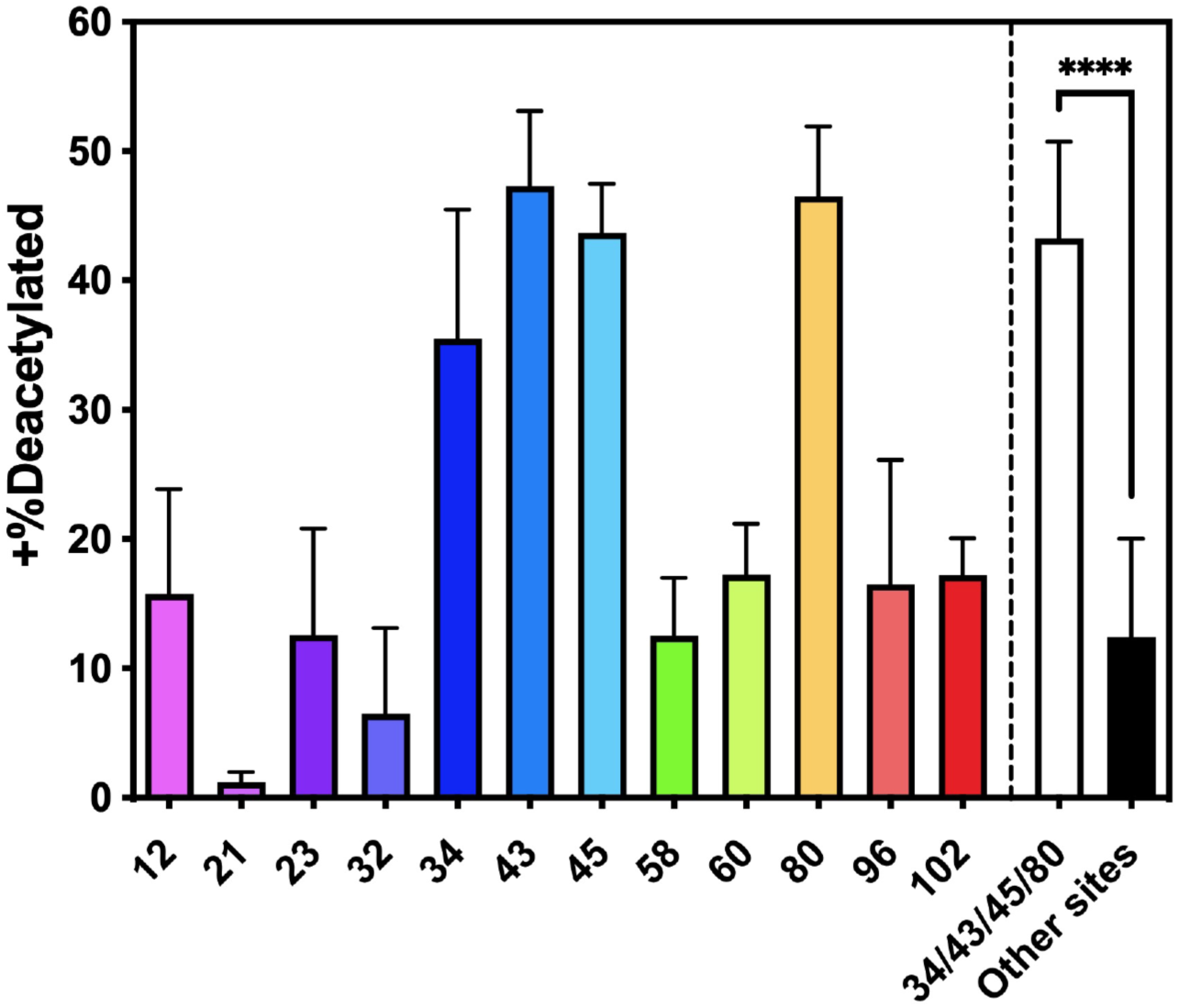
Site-specificity of HDAC activity. Samples of each of the acetylated αS variants were mixed with HDAC8 and after 24 h, acetylation levels were checked with a MALDI MS assay using ^15^N-labeled αS as a standard. Mean with SD, R=3

## Conclusion

In this study, we incorporated ^Ac^K site-specifically at all 12 currently known disease-relevant sites through ncAA mutagenesis and characterized the effects of this PTM on the physiological and pathological roles of αS using a variety of techniques. The aggregation assays showed that many of the Lys acetylations observed in patient samples have no effect, demonstrating that there is no non-specific effect on protein solubility or electrostatic interactions, at least for single Lys modifications. At sites 12, 43, and 80, Lys acetylation significantly slowed the elongation of fibrils *in vitro* and reduced PFF-based seeding in cells, showing that it can interfere with both phases of aggregation. Therefore, increasing acetylation at these sites through the use of KAT stimulators or KDAC inhibitors has potential therapeutic benefits. However, acetylation at Lys 12 or 43 perturbs membrane binding moderately, so increasing acetylation at these sites in αS could disrupt its native function in neurotransmitter vesicle trafficking. Thus, Lys 80 seems like the most promising site for targeted acetylation to drive αS toward conformations that are incompatible with known disease-related fibril polymorphs. Indeed, our previous cell-based studies using Gln mimics have shown that Lys 80 modification reduces aggregation (**Zhang *et al*., 2023**) By determining cryo-EM structures of ^Ac^K_80_ fibrils, we have provided a structural explanation for its inhibitory effects. Collectively, our results imply that strategies that can specifically enhance acetylation at Lys 80, without affecting Lys 12 or Lys 43, would be the most favorable approach to reduce αS aggregation pathology.

It should be noted that it is not clear at this point exactly what the acetylation levels at K_12_, K_43_ and K_80_ are in synucleinopathy patients. Taking advantage of our capability to produce authentically acetylated αS, we determined the extent of acetylation within human protein samples by quantitative liquid chromatography MS (Appendix 1 – figures 46-54). The ^Ac^K αS standards allowed us to correct for changes in trypsinization and ionization efficiency of acetylated peptides, the latter of which turned out to be very low for the ^Ac^K_80_ peptide due to its large size (a result of the missed K_80_ cut site due to acetylation, Appendix 1 – table 4). The level of acetylation was variable – no clear trend was observed between healthy control and patients – nor between patients of different diseases (Appendix 1 – table 3). Nevertheless, the MS data suggest that the 10 and 25% acetylation that we used for aggregation experiments are in the (patho)physiological range. Given the results reported here, it will be valuable to generate antibodies to acetylated peptides for the ^Ac^K_12_, ^Ac^K_43_ and ^Ac^K_80_ epitopes to more easily quantify the levels of acetylation in both soluble and fibrillar αS for immunofluorescence microscopy and Western blotting studies.

More broadly, our experiments show the value of a ncAA mutagenesis approach in systematically investigating a PTM that occurs at many locations in a protein. Since the yields were similar between NCL and ncAA mutagenesis, the ability to scan many sites by simple site-directed mutation to a TAG codon clearly makes ncAA mutagenesis the method of choice for our application. We efficiently scanned 12 different modification sites and fluorescently labeled proteins for binding studies. The ncAA approach was also crucial to generating isotopically labeled, acetylated αS for solution-phase NMR experiments and MS analysis. The isotopic labeling approach could be used in future solid-state NMR experiments to give detailed structural insight into slowed aggregation and distinct fibril morphology.

Our future experiments will include assessing the site-specificity of KATs and other KDACs for sites in αS in a similar fashion to the HDAC8 experiments here, studies enabled by our ability to easily produce ^Ac^K αS constructs. We can study modulation of HDAC8 and these other enzymes for their ability to specifically increase acetylation at Lys 80 without altering acetylation at Lys 12 or 43. We will also investigate effects on αS aggregation in cellular models using glutamine mutations that mimic Lys acetylation as well as small molecule modulators of HDAC8 or other enzymes to increase authentic acetylation levels. Such studies could verify our postulated effects on neurotransmitter trafficking, permit study of elongation effects due to (pseudo)acetylation of the monomer pool in a cellular context, and demonstrate the true viability of an HDAC8 inhibition strategy for modulation of αS pathology, in all three cases going beyond the *in vitro* results presented here. Furthermore, combining ncAA mutagenesis and NCL would allow us to study more complex PTM effects in αS, such as the combinatorial effects between multiple lysine acetylations or crosstalk between acetylation and other PTMs. As we have noted, it is likely that the PD/DLB and MSA fibril polymorphs are the result of an interplay of PTMs and cofactor interactions. While studies such as ours provide valuable insight into the effects of single PTMs and imply that increasing a PTM such as acetylation could inhibit aggregation, they are just a starting point to understanding how combinations of PTMs drive pathophysiological fibril polymorphs.

## Materials and Methods

### General Information

Reagents for peptide synthesis, including 2-(*1H*-benzotriazol-1-yl)-1,1,3,3-tetramethyluronium hexafluorophosphate (HBTU), *N*,*N*-diisopropylethylamine (DIPEA), and Fmoc-amino acids, were purchased from EMD Millipore (Burlington, MA, USA) or ChemImpex International (Wood Dale, IL, USA). Reagents for native chemical ligation (NCL): NaNO_2_, *tris*(2-carboxyethyl)phosphine (TCEP), and mercaptophenyl acetic acid (MPAA) were purchased from Sigma-Aldrich (St. Louis, MO, USA). *E. coli* BL21(DE3) cells and *E. coli* Dh5α cells were purchased from New England Biotechnologies (Ipswich, MA, USA). DNA oligomers were purchased from Integrated DNA Technologies, Inc (Coralville, IA, USA). DNA extraction and Miniprep kits were purchased from Qiagen (Hilden, Germany). Buffers were made with MilliQ filtered (18 MΩ) water (Millipore; Billerica, MA, USA). Preparation of the pTXB1-αS-intein-H_6_ plasmid containing α-synuclein (αS) with a C-terminal fusion to the *Mycobacterium xenopi* GyrA intein and C-terminal His_6_ tag was described previously (**Batjargal *et al*., 2015**). This plasmid was used as a starting point for the preparation of αS (mutants) -intein constructs. For overexpression of αS in HEK cells, the expression vector pcDNA5/TO was purchased from Thermo Fisher Scientific (Waltham, MA). pTECH-chAcK3RS (IPYE) was a gift from David Liu via Addgene (plasmid # 104069; http://n2t.net/addgene:104069; RRID:Addgene_104069; Watertown, MA, USA). Acetyllysine was purchased from ChemImpex. Nicotinamide was purchased from Alfa Aeser (Tewksbury, MA, USA). Atto 488 maleimide was purchased from Sigma-Aldrich. Matrix-assisted laser desorption/ionization mass spectrometer (MALDI-MS) data were collected with a Bruker Ultraflex III MALDI-MS instrument or a Bruker Microflex MALDI-MS (Billerica, MA, USA). UV/Vis absorbance spectra were obtained with a Hewlett-Packard 8452A diode array spectrophotometer (currently Agilent Technologies; Santa Clara, CA). Gel images were obtained with a Typhoon FLA 7000 (GE Lifesciences; Princeton, NJ, USA). Thioflavin T (ThT) absorbance spectra were collected on a Tecan SPARK plate reader (Mannedorf, Switzerland). Proteins were purified on a 1260 Infinity II preparative high-performance liquid chromatography (HPLC) system (Agilent Technologies). NCL reactions were monitored on a 1260 Infinity II Analytical HPLC system (Agilent Technologies) using a Jupiter C4 column (Phenomenex; Torrance, CA, USA). Water + 0.1% trifluoroacetic acid (TFA) (solvent A) and acetonitrile + 0.1% TFA (solvent B) were used as the mobile phase in HPLC.

### Protein semi-synthesis for generation of αS -^Ac^K_80._

To synthesize αS acetylated at K_80_, an N-to-C three-part native chemical ligation (NCL) was performed between the fragments αS_1-76_, αS_77-84_-^Ac^K_80_, and αS_85-140_. All fragments and intermediate products were purified by reverse-phase high-performance liquid chromatography (RP-HPLC) over a C4 column.

N-terminal thioester fragment αS_1-76_-MES (**1a**) and C-terminal fragment αS_85-140_-C_85_ (**4**) were constructed through deletion polymerase chain reaction (PCR) of previously published αS fragment constructs. They were each recombinantly expressed as a fusion with a polyhistidine-tagged GyrA intein from *Mycobacterium xenopi*. The N-terminal thioester was generated by adding excess sodium mercaptoethane sulfonate (MESNa) to cleave the intein by N,S-acyl shift.(**Muir *et al*., 1998**) (reported yield 24.1 mg/L.(**Pan *et al*., 2020a**)) Endogenous methionyl aminopeptidase processes the N-terminus of the 85-140 peptide to expose the N-terminal cysteine, (**Xiao *et al*., 2010**) which further reacts with aldehydes or ketones *in vivo* to form thiazolidine derivatives (**Liu *et al*., 2016**). The thiazolidine derivatives were deprotected with methoxyamine to give a free N-terminal cysteine (4.40 mg/L, SI Figure S1b).

The middle peptide αS_77-84_-Pen_77_^Ac^K_80_-NHNH_2_ (**2**, Pen: penicillamine (**Haase *et al*., 2008**)) was synthesized as a C-terminal acyl hydrazide through Fmoc-based, solid-phase peptide synthesis (Yield: 12.4 mg, 12 μmol, 48%, SI Figure S1a).

αS_1-76_-MES (**1a**) and αS_77-84_-Pen_77_^Ac^K_80_-NHNH_2_ (**2)** were ligated overnight under routine NCL conditions: 6 M Guanidine, 200 mM phosphate, 3 mM αS_1-76_-MES, 21 mM αS_77-84_-Pen_77_^Ac^K_80_-NHNH_2_, 100 mM MPAA, 50 mM TCEP, pH 7.0, 100 ul, 37 °C. (Yield: 1.46 mg, 172 nmol, 57%, SI Figure S1d). The purified product αS_1-84_-Pen_77_^Ac^K_80_-NHNH_2_ (**3a**) was activated to MES thioester (**3b**): 6 M guanidine, 200 mM phosphate, 0.5 mM **3a**, 5 mM NaNO_2_, pH 3.0, -15 °C, then 50 mM MESNa added and pH adjusted to 7.0, incubated at room temperature (Yield: 1.29 mg, 126 nmol, 73%, Figure S1c). The second ligation between αS_1-84_-Pen_77_^Ac^K_80_-MES (**3b**) and αS_85-140_-C_85_ (**4**) was performed in the presence of methyl thioglycolate (MTG) (**Huang *et al*., 2016**) to allow for desulfurization without intermediate purification (SI Figure S1e) Ligation: 6 M guanidine, 200 mM phosphate, 1.2 mM **3b**, 1.4 mM **4**, 120 mM MTG, 40 mM TCEP, pH 7.0, 37 °C; Desulfurization: 0.3 mM final concentration of ligation product (assuming full conversion), additional TCEP 250 mM, reduced glutathione 100 mM, VA-044 20 mM, pH 7.0, 37 °C. The product αS-^Ac^K_80_ (**5b**) was obtained in 43% yield (0.90 mg, 62 nmol, Figure S1f).

#### Construction of expression plasmids

The following primers were designed for site-directed mutagenesis to introduce TAG (= Z) codons at each lysine acetylation site. Site-directed mutagenesis for TAG mutations were performed on the plasmid encoding αS-MxeGyrA-His_6_.

Primer sequences:

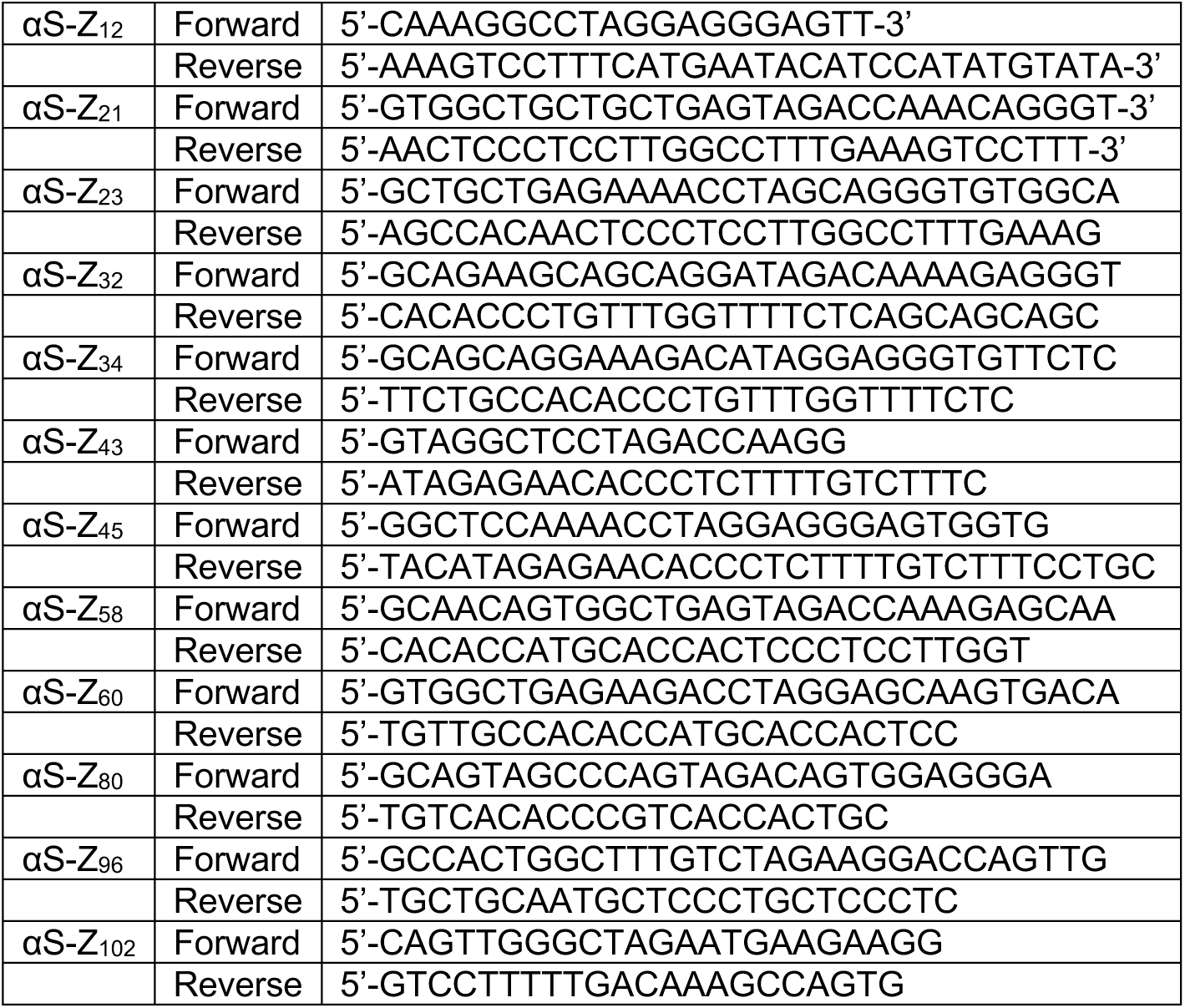

#### Production of recombinant αS constructs

To generate acetylated αS, each plasmid encoding αS with a TAG mutation at the site of interest and a machinery plasmid for acetyllysine incorporation, pTECH-chAcK3RS (IPYE), were co-transformed by heat shock at 42 °C into BL21 (DE3) competent cells. Cells were plated and incubated on ampicillin/chloramphenicol (Amp/Chlor) plates. Single colonies were picked to inoculate primary cultures in LB media supplemented with 0.1 mg/mL Amp and 0.025 mg/mL Chlor. Primary cultures were incubated overnight or until they were cloudy at 37 °C. Secondary cultures in LB media were inoculated and grown at 37 °C with shaking at 250 rpm until the optical density reached ∼ 0.6. The culture was subsequently cooled to 18 °C. 50 mM nicotinamide and 10 mM ε-acetyl lysine were added to the culture and incubated for ∼ 5 min before inducing the expression of the gene of interest with 1 mM isopropyl ß-D-1-thiogalactopyranoside (IPTG). To generate isotopically labeled, acetylated αS, protein expression was performed as above, except that cells were grown in M9 minimal media (**Anderson, 1946**) that contains ^15^N-ammonium chloride. Induced cells were then grown in the shaker-incubator at 18 °C overnight. After centrifugation (4000 rpm, 20 min, 4 °C), cell pellets were re-suspended in buffer (40 mM Tris pH 8.3, with one Roche protease inhibitor tablet) and sonicated in a cup in an ice bath (5 min, 1 s ON, 2 s OFF). The resulting lysate was centrifuged (14,000 rpm, 30 min, 4 °C), and supernatant containing the αS variant was purified over a Ni-NTA affinity column. Intein cleavage was carried out by incubation with 200 mM β-mercaptoethanol (βME) on a rotisserie overnight at room temperature. Cleaved αS variant was dialyzed into 20 mM Tris, pH 8 buffer before purification over a second Ni-NTA column to remove the free intein from the sample. The αS proteins were purified by RP-HPLC over a C4 column (acetylated proteins) or by fast-protein liquid chromatography (FPLC) using a Hi-Trap Q 5 mL column (glutamine mutants), dialyzed into 1x phosphate buffered saline (PBS) and spin-concentrated. For purification of acetylated Cys mutants for fluorescent labeling, TCEP was added to a final concentration of 1 mM prior to HPLC injection and samples were dialyzed into 20 mM Tris 50 mM NaCl pH 7.4 after purification. Upon flash-freezing, protein stocks were kept at -80 °C until further use. Isotopically labeled αS samples were lyophilized after HPLC purification.

#### Fluorescent labeling

To label αS-^Ac^K_X_C_114_, the protein stocks in 20 mM Tris, 50 mM NaCl, pH 7.4 were incubated with 2-10 eq. TCEP, then 10 eq. Atto 488-maleimide dye was added and incubated at room temperature for 2-4 hours or at 4 °C for overnight until product formation was observed by MALDI-MS. The product was purified by HPLC over a C4 column and dialyzed into 20 mM Tris, 50 mM NaCl, pH 8.

#### Circular Dichroism (CD)

αS samples were filtered through 100 kDa-cutoff spin concentrators and the concentrations were determined by UV-Vis absorbance. Based on the quantification, all the αS samples were first diluted to 15 μM with 1x PBS. Samples for CD acquisition were then prepared in triplicate by dilution of the protein stock solutions using 4.3 mM Na_2_HPO_4_, 1.47 mM KH_2_PO_4_, pH 7.4, and mixing with 100 mM sodium dodecyl sulfate (SDS) in 4.3 mM Na_2_HPO_4_, 1.47 mM KH_2_PO_4_, pH 7.4 to yield samples composed of 5 μM αS with 10 mM SDS in 45.7 mM NaCl, 0.9 mM KCl, 4.3 mM Na_2_HPO_4_, 1.47 mM KH_2_PO_4_, pH 7.4.

CD spectra were acquired on a JASCO J-1500 spectrometer in quartz cuvettes with a path length of 1 mm at 25 °C. Spectra were collected over the range of 190-260 nm using a 1 nm data pitch, 2 nm bandwidth, 8 s data integration time and 50 nm/min scanning speed. At each wavelength, spectra were background subtracted for buffer blank and corrected for concentration, path length, and number of residues as described by the following equation, where 𝜃_sample_ and 𝜃_blank_ refer to raw ellipticity values for the sample and buffer blank, respectively, ε is the path length of the cuvette, *c* is the protein concentration, and *n* is the number of residues.

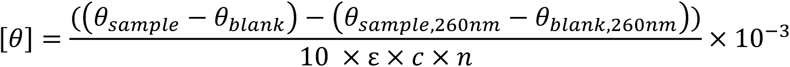

To determine the effects on helicity, the [𝜃_+++_] value acquired for each acetylated αS was normalized by the [𝜃_+++_] value of αS WT.

#### Protein aggregation kinetics and percentage incorporation into fibrils

Aggregation of αS monomer seeded by preformed fibrils (PFFs) of wild type (WT) αS was performed with 100% WT αS or with mixtures of 10% or 25% acetylated αS or αS Gln mutants. The seeding experiments were carried out by agitation at 1400 rpm at 37 °C and monitored by ThT. αS WT monomer in buffer (1x phosphate-buffered saline (PBS), pH 7.4) was prepared as a 40 μM stock solution and a 50 μL aliquot was added to each well of a 96-well half area clear bottom plate (6 replicates per one construct). The plate was sealed with a plastic film and incubated at 37 °C for 30 min before aggregation. PFF seeds were prepared by resuspending fibrils in PBS to make a 4 μM stock solution and freshly sonicating in an Eppendorf tube in an ice bath (2 min, 1 s ON, 1 s OFF). A 50 μL aliquot of sonicated seeds was added to monomers in each well (10% seeds). Samples were shaken on an IKA MS3 orbital shaker set to 1400 rpm at 37 °C. At each time point, ThT fluorescence was measured on a Tecan SPARK plate reader (excitation: 450 nm, emission: 485 nm, emission bandwidth: 5 nm, integration time: 40 μs). The extent of aggregation was determined based on normalized fluorescence intensity at 485 nm calculated from the minimum intensity and maximum intensity of each replicate. The data points were fit using GraphPad Prism software with the nonlinear regression model using the following equation:

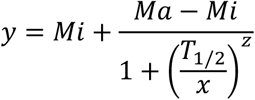

where Mi and Ma are the minimal and maximum y values, respectively, T_1/2_ is the time at the mid-point of aggregation, and z is a parameter that determines the steepness of the curve. For the plots in Figure 4, Aggregation time was normalized to T_1/2_ of the WT control, performed in parallel, and minimum and maximum fluorescence values were also normalized.

After the final time point, samples were pelleted at maximum speed on a tabletop centrifuge for 90 min. The supernatant was removed, and pellet was resuspended in the original volume of buffer. Samples were supplemented with SDS to a 25 mM final concentration, boiled for 20 min, and chilled on ice. Monomeric samples for calibration were prepared by 2-fold serial dilutions in water. All samples were analyzed by SDS-PAGE (4-15 or 4-20% acrylamide). Gels were stained with Coomassie Brilliant Blue dye. Quantification of the intensity of bands was done using the ImageJ software (National Institutes of Health; Bethesda, MD, USA. Values reported for aggregation kinetics and monomer incorporation are the average and standard error of mean taken from independent replicates.

#### In situ aggregation in cultured primary hippocampal neurons

Primary hippocampal neurons were isolated from mouse neonate brain and plated on a Thermo Fisher Scientific 96-well plate at a density of 350 cells/µL in 100µL of neurobasal (NB) media with 5% Fetal Bovine Serum. After 24 hours of incubation, the media was replaced by aspiration with fresh NB media.

PFFs were formed by agitation for 3 days at 3500 rpm at 37 °C for both WT αS and post-translationally modified variants for the following conditions: 25% ^Ac^K_12_, 25% ^Ac^K_43_, 25% ^Ac^K_80_. At 8 days in vitro (DIV), PFF constructs were sonicated in (QSonica Microson XL-2000) for 20 cycles (1s ON, 1s OFF) and used to treat the neurons for a final concentration of 50 ng/µL. The neurons were cultured for an additional 14 days following treatment of the fibril seeds, during which the medium was exchanged every 7 days. At 21 DIV, the cells were fixed with a final concentration of 4% paraformaldehyde (Electron Microscopy Sciences) + 4% sucrose (Sigma-Aldrich) in PBS, washed 3x with PBS (Sigma-Aldrich), and permeabilized with 0.1% Triton X-100 (Thermo Fisher). After 3x washes with PBS, the plate was then incubated overnight in fluorescent western blot blocking buffer (Rockland Immunochemical) and probed with 1:3000 phosphoserine 129 antibody (m81A (**Waxman and Giasson, 2008**)) for detection of intracellular αS aggregates and neuronal cytoskeleton antibody (MAP2 (**Volpicelli-Daley et al., 2011**)) for detection of neuronal processes in blocking buffer at room temperature for 4 hours. Samples were washed 3x with PBS and then incubated with 1:1000 fluorophore-conjugated secondary antibodies in blocking buffer at room temperature for 1 hour. The plate was then immediately washed 3x with PBS and imaged in 75 µL of PBS (IN Cell Analyzer 2200/2000). Quantification of 81A signal was done using ImageJ. All ^Ac^K variants were found to have significantly less *in situ* recruitment compared to WT by One-Way ANOVA and Tukey post-hoc (GraphPad Prism 9.3.1).

#### Preparation of synthetic vesicles

For NMR experiments, 60:25:15 1,2-dioleoyl-sn-glycero-3-phosphocholine:1,2-dioleoyl-sn-glycero-3-phosphatidylethanolamine:1,2-dioleoyl-sn-glycero-3-phospho-L-serine (DOPC:DOPE:DOPS) small unilamellar vesicles (SUVs) were generated as described previously(Dikiy and Eliezer, 2014) to mimic the size and lipid composition of native synaptic vesicles. In brief, mixtures of DOPC, DOPE, and DOPS dissolved in chloroform (Avanti Polar Lipids, Alabaster, AL) were dried under nitrogen the lipid film was resuspended in NMR buffer (10 mM Na2HPO4, 100 mM NaCl, 10% v/v D2O, pH 6.8), immersed in a bath sonicator for 15 min at a time until clear, and further clarified by ultracentrifugation at 130,000 × g for 2 h. The supernatant was stored at 4 °C as SUV stock solution and was used for NMR experiments within 1 day.

For fluorescence correlation spectroscopy, lipid vesicles were prepared by extrusion through porous membranes. A mixture in 50:50 molar ratio of 1-palmitoyl-2-oleoyl-sn-glycero-3-phosphoserine (POPS) and 1-palmitoyl-2-oleoyl-sn-glycero-3-phosphocholine (POPC) were drawn from chloroform stock and dried under nitrogen gas to form a film inside a glass vial. Films were desiccated under vacuum and re-hydrated in 20 mM 3-(*N*-morpholino)propanesulfonic acid (MOPS), 147 mM NaCl, 2.7 mM KCl, pH 7.4. Ten freeze-thaw cycles consisting of cooling in liquid nitrogen for 40 s and warming in a 60 °C water bath for 2 min were performed to aid the formation of uniformly sized vesicles. With syringes, vesicles were then extruded 31 times through stacked 50 nm pore membranes held in place inside an extruder. Vesicles were determined by dynamic light scattering (DLS) to be monodisperse and distributed uniformly around 80 nm in diameter, consistent across different concentrations of all samples. All lipid vesicles were prepared fresh and used within 48 h of extrusion.

#### Heteronuclear single quantum coherence NMR spectroscopy (HSQC)

Lyophilized WT or lysine-acetylated αS mutants were dissolved in NMR buffer (10 mM Na_2_HPO_4_, 100 mM NaCl, 10% v/v D_2_O, pH 6.8) and mixed in a 1:1 volume ratio with SUV stock solution or NMR buffer. Final protein concentrations were ca. 100 µM and final SUV lipid concentrations were ca. 3 mM, assuming complete conversion of lipids to SUVs. NMR ^1^H-^15^N heteronuclear single quantum coherence (HSQC) experiments were acquired at 10 °C using a 500 MHz Bruker Avance spectrometer equipped with a cryogenic probe using TopSpin 3.2 software. Data processing and analysis were performed using NMRbox (**Maciejewski et al., 2017**), NMRPipe (**Delaglio et al., 1995**), and NMRFAM-SPARKY (**Lee et al., 2015**) software. Amide resonance peaks were assigned based on previously published chemical shift assignments for WT αS monomer (**Eliezer *et al*., 2001**). NMR peak intensity ratios were calculated as the ratio of peak intensity in the presence of SUVs to that in the absence of SUVs for matched samples. To correct for protein concentration variations in the samples, the intensity ratios were normalized by the average ratio for the C-terminal 40 residues, which do not feature appreciable membrane interactions at these lipid concentrations.

#### Fluorescence correlation spectroscopy (FCS)

Fluorescence correlation spectroscopy (FCS) experiments to study the binding of αS-^Ac^K_43_C^488^_114_ and αS-^Ac^K_80_C^488^_114_, including preparation of synthetic lipid vesicles, collection of FCS data, and data analysis, were carried out as described previously for arginylated αS.(Pan *et al*., 2020a) Eight-well chambered coverglasses (Nunc, Rochester, NY, USA) were prepared by plasma cleaning followed by incubation overnight with polylysine-conjugated polyethylene glycol (PEG-PLL), prepared using a modified Pierce PEGylation protocol (Pierce, Rockford, IL, USA). PEG-PLL coated chambers were rinsed with and stored in Milli-Q water until use. FCS measurements were performed on a lab-built instrument based on an Olympus IX71 microscope with a continuous emission 488 nm DPSS 50 mW laser (Spectra-Physics; Santa Clara, CA, USA). All measurements were made at 20 °C. The laser power entering the microscope was adjusted to 4.5 μW. Fluorescence emission collected through the objective was separated from the excitation signal through a Z488rdc long pass dichroic filter and an HQ600/200m bandpass filter (Chroma; Bellows Falls, VT, USA). Emission signal was focused onto the aperture of a 50 μm optical fiber. Signal was amplified by an avalanche photodiode (Perkin Elmer; Waltham, MA, USA) coupled to the fiber. A digital autocorrelator (Flex03Q-12, correlator.com; Bridgewater, NJ, USA) was used to collect 10 autocorrelation curves of 10 seconds for each measurement of free protein in buffer without lipids or 30 autocorrelation curves of 30 seconds for each measurement in the presence of lipid vesicles. Fitting was done using lab-written code in MATLAB (The MathWorks; Natick, MA, USA).

To determine the diffusion time of each protein construct, each αS variant labeled with Atto 488 was measured in buffer without lipid. The average of 10 autocorrelation curves was fit to a 1-component autocorrelation function:

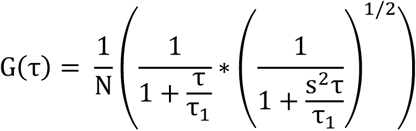

where G(τ) is the autocorrelation function, N is the number of molecules in the focal volume, τ_0_ is the diffusion time of αS, and s is the radial-to-axial ratio of the excitation volume. The counts per molecule (CPM) for each sample was calculated by dividing the average intensity (Hz) of the measured signal by the number of molecules N. The normalized CPM of each αS was calculated by dividing by the CPM of freely diffusing fluorescent standard Alexa Fluor 488.

#### Vesicle binding affinity

αS constructs labeled with Atto488 were examined in the presence of varying concentrations (0.001 mM to 0.5 mM lipid) of lipid vesicles consisting of 50:50 POPS/POPC. The average of 30 autocorrelation curves was fit to a 2-component equation:

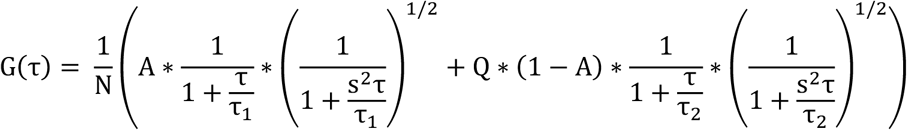

where G(τ) is the autocorrelation function, N is the number of molecules in the focal volume, τ_0_ is the characteristic diffusion time of αS, τ_+_ is the characteristic diffusion time of the vesicles, s is the radial-to-axial ratio of the excitation volume, Q is the ratio of the brightness of vesicle-bound αS relative to αS, and A is the fraction of free αS. When fitting the autocorrelation curves for αS in the presence of lipid vesicles, the diffusion time of bound and unbound αS were respectively fixed to experimentally determined values. The diffusion time of unbound protein, τ_1_, was determined by measurements of the protein in buffer without lipids. Since bound protein diffuses with the vesicles to which they are bound, the diffusion time of the vesicles, τ_2_, was determined by measurements of the protein in the presence of a concentration of vesicles that gave the maximum diffusion time (0.1 mM lipid). In the binding assay, the fraction of αS bound at each lipid concentration was obtained from the fit to each autocorrelation curve. Averages and standard deviations were calculated from at least 3 independent measurements performed on separate days at each lipid concentration. The resulting binding curve was fit to the following equation, from which the K_d,app_ was determined.

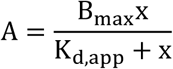

where A is the fraction of αS bound, x is the accessible lipid concentration, B_345_ is the maximum fraction of αS bound, and K_6,477_ is the apparent dissociation constant.

#### Transmission electron microscopy (TEM)

TEM imaging was carried out on an FEI Tecnai T12 instrument (Hillborough, OR, USA) with an accelerating voltage of 100 kV. Fibril samples were prepared by shaking 100 μM αS (WT with 25% ^Ac^K variants) at 1500 rpm for 72 hours, then diluted into water at a final concentration of 0.1 mg/mL. A 5 μL drop of sample was deposited on glow discharged carbon Formvar coated 300-mesh Cu grids and allowed to rest for 1 min at room temperature. 5 μL of stain (2% w/v uranyl acetate in water) was then applied to the grid. The liquid was wicked off with grid paper, and another 5 μL of stain was applied and wicked off. Images were collected at magnifications ranging from 11000x to 42000x.

#### Cryo-electron microscopy (Cryo-EM) data acquisition

PFFs for cryo-EM studies were prepared as following: αS monomer in buffer (PBS: 1x PBS, pH 7.4; TBS: 20 mM Tris 140 mM sodium chloride pH 7.0 at 37 °C) was prepared as 100 μM stock solutions. The monomers were incubated at 37 °C for 30 minutes prior to shaking to ensure that the same temperature is adjusted. Samples were then shaken at 1500 rpm at 37 °C in Eppendorf tubes for overnight.

Prior to vitrification, fibril samples were concentrated 10-fold by centrifugation at 14,000 rpm for 10 minutes using a Beckman Microfuge 18 centrifuge (Beckman Coulter; Indianapolis, Indiana) to a final concentration of approximately 1 mM. A 3.5 μl aliquot was applied to a glow-discharged holey copper grid (Quantifoil R2/1, 300 mesh) equilibrated at 4 °C and 100% humidity. After a 1-second blot, the grids were plunge-frozen into liquid ethane using a Vitrobot Mark IV (Thermo Fisher). Cryo-EM images were acquired on a 200 kV Talos Glacios microscope equipped with a Falcon 4i camera and Selectris energy filter. The magnification was set to 100,000X, resulting in a calibrated pixel size of 1.16 Å/pix. To minimize radiation damage, the total dose applied to the samples was limited to 40 e-/Å² delivered over a 5.01-second exposure time using 1539 movie frames with electron event representation (EER) fractions of 27. A defocus range of -2.5 to -0.8 μm was employed along with a spherical aberration of 2.7 mm. The microscope was operated at 200 kV with a 50 μm objective aperture and a 20 eV energy filter. For further analysis, a total of 5387, 4038, and 5001 images were collected for 25% ^Ac^K_80_, 100% ^Ac^K_80_, and WT αS fibril preparations, respectively.

#### Cryo-EM data processing

Cryo-EM data processing began with CryoSPARC v4.1.1 (Structura Biotechnology Inc.; Toronto, Canada) (**Punjani *et al*., 2017**). EER data files were imported with 27 fractions and dose rate of 1 e^-^/Å^2^/fraction. Patch Motion correction and Patch CTF estimation were applied to the imported data. For initial filament picking, template-free filament tracing was used on all images, targeting filaments with diameters between 50 and 250 nm. Particles of size 768x768 pixels with 12.5 or 50 Å inter-box distance were extracted and subjected to 2D classification (see Table 1 for numbers of template-free filament tracing particles for each fibril sample). The best 3 classes were then selected and used to create a template for a second round of filament tracing on the entire dataset. Following filament tracing, Particles were again extracted from the micrographs with a box size of 512 pixels and underwent 3 rounds of 2D classification with only a subset of particles retained (see Table 1 for numbers of extracted particles and retained for each fibril sample).

CryoSPARC (.cs) files were then converted to RELION STAR files using the PyEM csparc2star.py script. Following this data conversion, a custom Python script developed with ChatGPT precisely determined the start and end coordinates of each fibril. This script processed the csparc2star.py star file output to create coordinate pairs based on the cryoSPARC Fibril ID, notably without transferring the original tilt and psi angles from CryoSPARC to RELION (**Kimanius *et al*., 2021**). The output of this custom script served as the input for the autopick RELION extraction step.

To handle curved fibrils, a special segmentation strategy was implemented: the script traced the coordinates along the fibril, generating a new start and end coordinate pair for individual segments, with the segment’s end coordinate assigned either after spanning 10 particles (around 50 nm in length) or when the end of the fibril was reached. This resulted in shorter, straighter segments for processing. Finally, the createAutopick function from cryolo_boxmanager_tools.py in crYOLO was utilized to generate a STAR file linking the final particle coordinates to their movie files, which was then used to perform particle extraction in RELION. The standard RELION image processing pipeline then followed, including 2D classification, refinement, and 3D classification.

This approach resolves the issue with curved fibrils, but also results in all picked fibrils being 50 nm or shorter in RELION. Consequently, segments from the same fibril receive unique fibril IDs in RELION and may be split into different half-maps. Although this avoids random splitting of individual particles without regard to their origin from the same fibril, it could still cause an overestimation of resolution.

To assess any potential overestimation of resolution, another script was created to reassign the fibril ID of each particle in the RELION STAR file to that of the closest particles in the cryoSPRAC data, effectively ensuring that all particles from an individual fibril have the same fibril ID and are not assigned to different half-maps. This revised STAR file was then used as the input images STAR file for 3D auto-refinement in RELION. A comparison of maps before and after reassigning the fibril IDs shows negligible effects on the resulting structures and on their estimated resolution, indicating that the original procedure did not results in any significant overestimation of the resolution.

For 25% αS-^Ac^K_80_ preparations, particles were re-extracted in Relion 4.0.0 (Kimanius *et al*., 2021) using 3 unique asymmetric units and box size of 256 pixels (see Table 1 for numbers of re-extracted particles). 2D classification was then conducted using Relion with Regularization parameter (T) of 2, in-plane angular sampling of 2 degrees and 20 iterations using the EM algorithm. An initial model was generated from the good 2D classes using the relion_helix_inimodel2d function in Relion (see Table 1 for numbers of initial model particles). 3D auto-refinement was employed to obtain the cryo-EM map with soft-edge mask (see Table 1 for numbers of selected 3D classification particles). A final map with resolution of 2.88 Å was obtained after particle Bayesian polishing and CTF refinement.

For WT and 100% αS-^Ac^K_80_ fibril preparations (from both TBS and PBS), particles were re-extracted in Relion 5.0-beta-1(Burt et al., 2024) using 3 unique asymmetric units and a box size of 256 pixels (see Table 1 for numbers of re-extracted particles). Initial maps were generated using the helical refinement function in CryoSPARC (see Table 1 for numbers of initial model particles for each fibril sample). 3D auto-refinement in Relion was employed to obtain the cryo-EM map with soft-edge masks. However, blurry maps were observed for both WT αS and ^Ac^K_80_ αS in one of two chains, indicating the presence of multiple structures (**Lövestam and Scheres, 2022**). Two runs of 3D classification were then performed with 50 iterations, with alignment and optimization of twist and rise parameters to separate the two structures (see Table 1 for numbers of selected 3D classification particles for each fibril sample). Two maps with similar backbone traces but different alignments were obtained (A and B polymorphs for WT and αS-^Ac^K_80_ for both PBS and TBS preparations, see Figure 8). Final maps were obtained after particle Bayesian polishing and CTF refinement.

#### Model building and refinement

For 25% αS-^Ac^K_80_ preparation, the cryo-EM map was first fitted with the PDB ID 6a6b structure in Chimera (**Li *et al*., 2018b**). The resulting atomic model was then adjusted in Coot according to the map. Next, the model was refined in real space using PHENIX 1.21.1 (**Liebschner *et al*., 2019**). For cross validation, all atoms in the refine model were randomly displaced 0.3 Å and refined against the first half map from Relion 3D auto-refinement. Final statistics for the 25% αS-^Ac^K_80_ preparation given in SI Table S1.

For WT (TBS) and 100% αS-^Ac^K_80_ (PBS) fibril preparations, cryo-EM maps were first fitted with a single chain of the PDB ID 6rt0 structure (**Guerrero-Ferreira *et al*., 2019**) for WT αS and a single chain of the PDB ID 8pix structure (**Frey *et al*., 2024**) for αS-^Ac^K_80_ in Chimera. The resulting atomic models were then adjusted, and layers were added in Coot according to the map. Next, models were refined in real space using PHENIX 1.21.1. For cross validation, all atoms in the refined model were randomly replaced by 0.3 Å and refined against the first half map from Relion 3D auto-refinement. Final statistics for the WT and αS-^Ac^K_80_ fibril polymorphs are given in SI Table S2.

#### Preparation of human HDAC8

Recombinant human HDAC8 was expressed and purified as previously described with minor modifications **(Decroos *et al*., 2014; Dowling *et al*., 2008; Osko *et al*., 2021**). Briefly, the HDAC8-6His-ET20b expression plasmid was grown in BL21-DE3 cells, and a 250 mL culture was incubated overnight in Lysogeny Broth (LB) media supplemented with 100 µg/mL of ampicillin at 37 °C. Aliquots of this culture were used to inoculate 6 x 1 L cultures of 2XYT media supplemented with 100 µg/mL of ampicillin. Cultures were grown at 37 °C until reaching OD = 0.6–0.8; after cooling to 18 °C, cultures were induced by the addition of 0.4 mM isopropyl-β-D-thiogalactopyranoside (IPTG) and 0.5 mM ZnSO_4_ and grown overnight. Cells were pelleted by centrifugation (5244g, 15 min). Purification was achieved exactly as recently summarized (**Osko *et al*., 2021**) with slight modification of the lysis buffer: 50 mM Tris (pH 8.0), 500 mM KCl, 5% w/v glycerol, and the reducing agent 1 mM TCEP in place of 3 mM β-mercaptoethanol (BME).

#### HDAC8 deacetylation assay

Acetylated αS substrates (αS ^Ac^K_X_) and ^15^N-labeled αS WT were buffer-exchanged into HDAC8 reaction buffer (50 mM Tris, 137 mM NaCl, 2.7 mM KCl, 1 mM MgCl_2_). Concentrations of each were quantified and deacetylation reaction was setup in triplicate, with final concentrations of HDAC8 8μM, 1.25 μM αS ^Ac^K_X_, 1.25 μM ^15^N-αS WT. The reaction was performed at room temperature for 24 h. MALDI-TOF-MS spectra were collected for each reaction, at 0h and 24h timepoints. Peak picking and peak area quantification, via fitting to Gaussian curves, for deacetylated product (^14^N-αS WT) and the standard (^15^N-αS WT) were performed on the Bruker flexAnalysis software.

The formula below was used to calculate the increase in %Deacetylated.

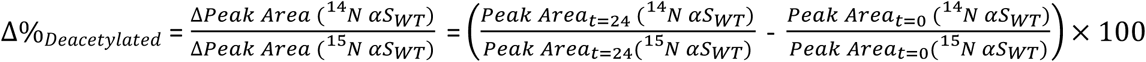

#### Proteomic analysis of acetylated αS standards

Each pure α-synuclein protein (20 µg) was denatured in 40 µL of 8 M urea in PBS, reduced by 5 mM of TCEP with 30 min incubation at 37 °C, alkylated by 15 mM of iodoacetamide (IAA) with 30 min incubation at room temperature in the dark. The solution was diluted to 2 M urea by 50 mM ammonium bicarbonate in H_2_O, digested by trypsin (sequence grade, Promega) at 1:50 trypsin/protein ratio (*w*/*w*) with overnight (∼12 h) incubation at 37 °C. The resulted peptide solution was acidified by formic acid at a final concentration of 5%. The peptide solution was desalted, resuspended in 0.1% formic acid at ∼100 ng/µL. An LC-MS/MS system consisted of an Vanquish Neo UHPLC coupled to a Orbitrap Ascend (Thermo Scientific) was used for peptide analysis. Peptide samples were maintained at 7 °C on sample tray in LC. Separation of peptides was carried out on an Easy-Spray™ PepMap™ Neo nano-column (2 µm, C18, 75 µm X 150 mm) at room temperature with a mobile phase consisting of a linear gradient of A (0.1% FA in H_2_O) and B (acetonitrile containing 0.1% FA) under the following conditions: 0 → 90 →110 → 110.5 → 116 min, 0% → 28% → 36% → 98% → 98% B. The flow rate was 300 nL/min. 200 ng of each sample was injected for LCMS analysis in data-independent mode. The voltage applied to the nano-LC electrospray ionization source was 1.9 kV. The temperature of ion transfer tube (ITC) was set at 275 °C. Spectra were collected in a data-dependent acquisition mode such that each scan cycle (3 sec) involved a single high-resolution (120,000) full MS spectrum of parent ions (MS1 scan from m/z 350–2000) collected in the orbitrap. Parent ions assigned as peptide in charge states +2-6 with intensity higher than 2E4 were included for fragmentation. HCD-induced fragmentation (MS2) scans were recorded in orbitrap (scan from m/z 120–1500). Dynamic exclusion was set as repeat count of 1 within exclusion time of 20 s. All other parameters were left as default values. Relative ionization efficiency of the acetylated peptide over unmodified peptide was calculated based on the peptide intensity in acetylated αS compared to WT αS.

#### Quantitative analysis of αS Lys acetylation in human samples

Raw data were downloaded from previous works (50 raw files from PMID37814027(Zhang *et al*., 2023) and 7 raw files from PMID32461689 (**Schweighauser *et al*., 2020**)). Data was searched using the Byonic software (v4.5.2) against a reverse-concatenated, nonredundant database of the human proteome. Cysteine residues were searched with a static modification for carbamidomethylation (+57.02146 Da). Methionine residues were searched with up to two differential modifications for oxidation (+15.9949 Da). Peptide is allowed to up to two acetylation (+42.0106 Da) on lysine residues. Peptides were required to have two tryptic terminals and up to two missed cleavages were allowed in the database search. The parent and daughter ion mass tolerances for a minimum envelope of three isotopic peaks was set to 10 and 20 ppm, respectively. The false-positive rate was set at 1% or lower. The identified peptides from αS were listed in Supplementary Data 1 from both published datasets. MS1 peak area was used for acetylated and unmodified peptide quantification, only the raw files with identified acetylation sites of interest were included for analysis. Skyline software (v22.2.0.527) was used to export peak areas and data visualization.

## Supporting Information

Additional figures and tables for plasmid mutagenesis, protein semi-synthesis and ncAA mutagenesis, fluorescent labeling, CD, aggregation measurements, neuron seeding, TEM, lipid vesicle preparation, FCS measurements, NMR data acquisition, cryo-EM structure determination, and MS analysis are provided in Supporting Information.

## Acknowledgements

This research was supported by the National Institutes of Health (NIH RF1NS125770 to D.E., E.R., and E.J.P.; RF1NS103873 to E.J.P.; and RF1AG066493 and R35GM136686 to D.E., R01NS111997 and R01HD106051 to B.A.G., and R01GM049758 to D.W.C.). Instruments supported by the National Science Foundation and NIH include NMR (NSF CHE-1827457 and NIH S10OD028556 and S10OD016320) and MALDI MS (NIH S10OD030460). We also thank the Electron Microscopy Resource Lab (RRID: SCR_022375) for the use of their instruments. M.S. thanks the Nakajima Foundation for scholarship funding. J.R. was supported by the NIH Chemistry Biology Interface Training Program (T32GM133398). Z.L. thanks a Washington University School of Medicine BMB Seed Grant (PJ000027587) and support from the Research Education Component through the P30AG066444 grant. Some of the work was performed at the WCM Cryo-EM Core Facility and we acknowledge assistance from Carl Fluck with cryo-EM data collection and analysis.

## Appendix 1

**Figure 1.**
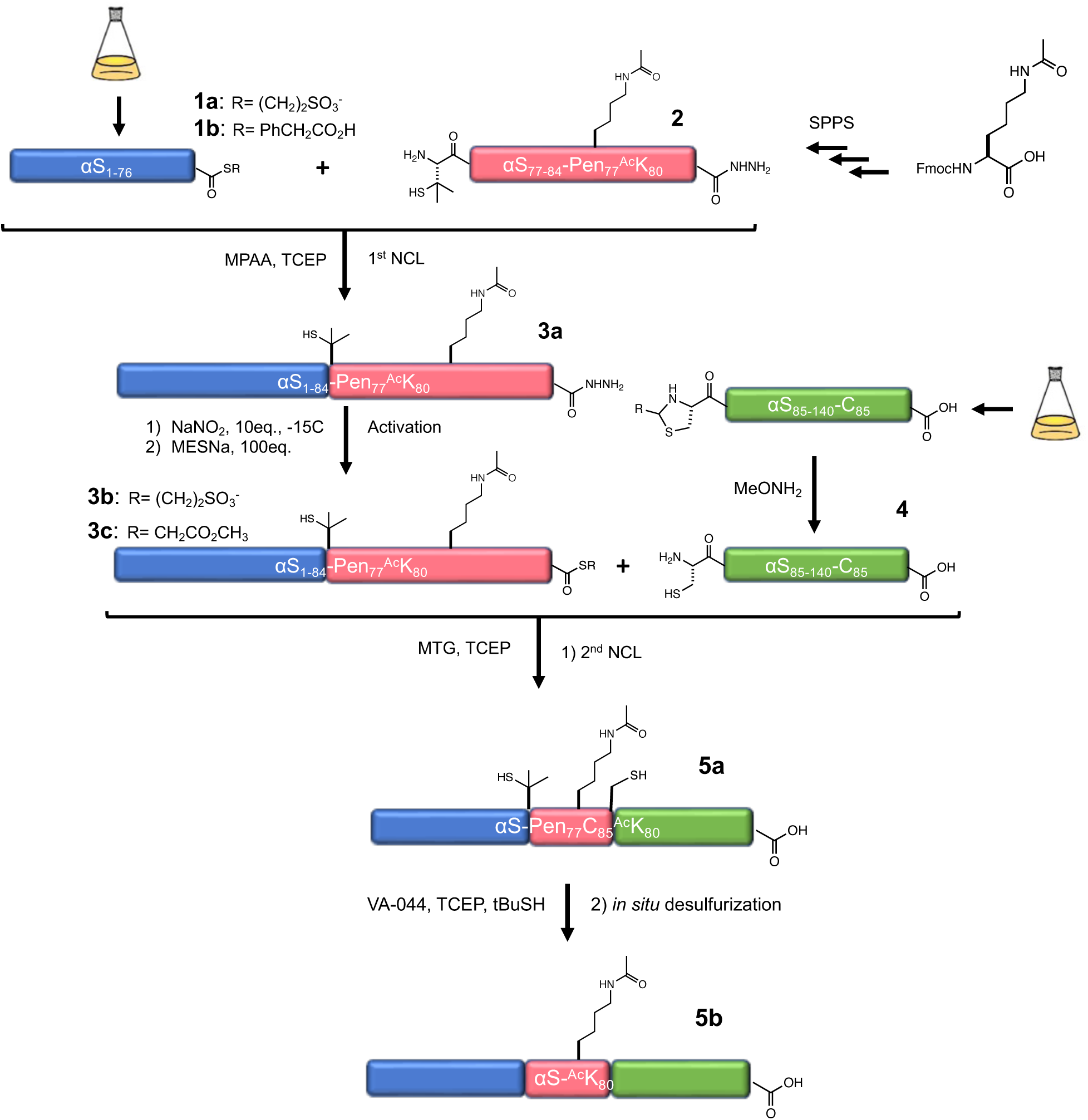
Semi-synthesis scheme for creating αS-^Ac^K_80._

**Figure 2.**
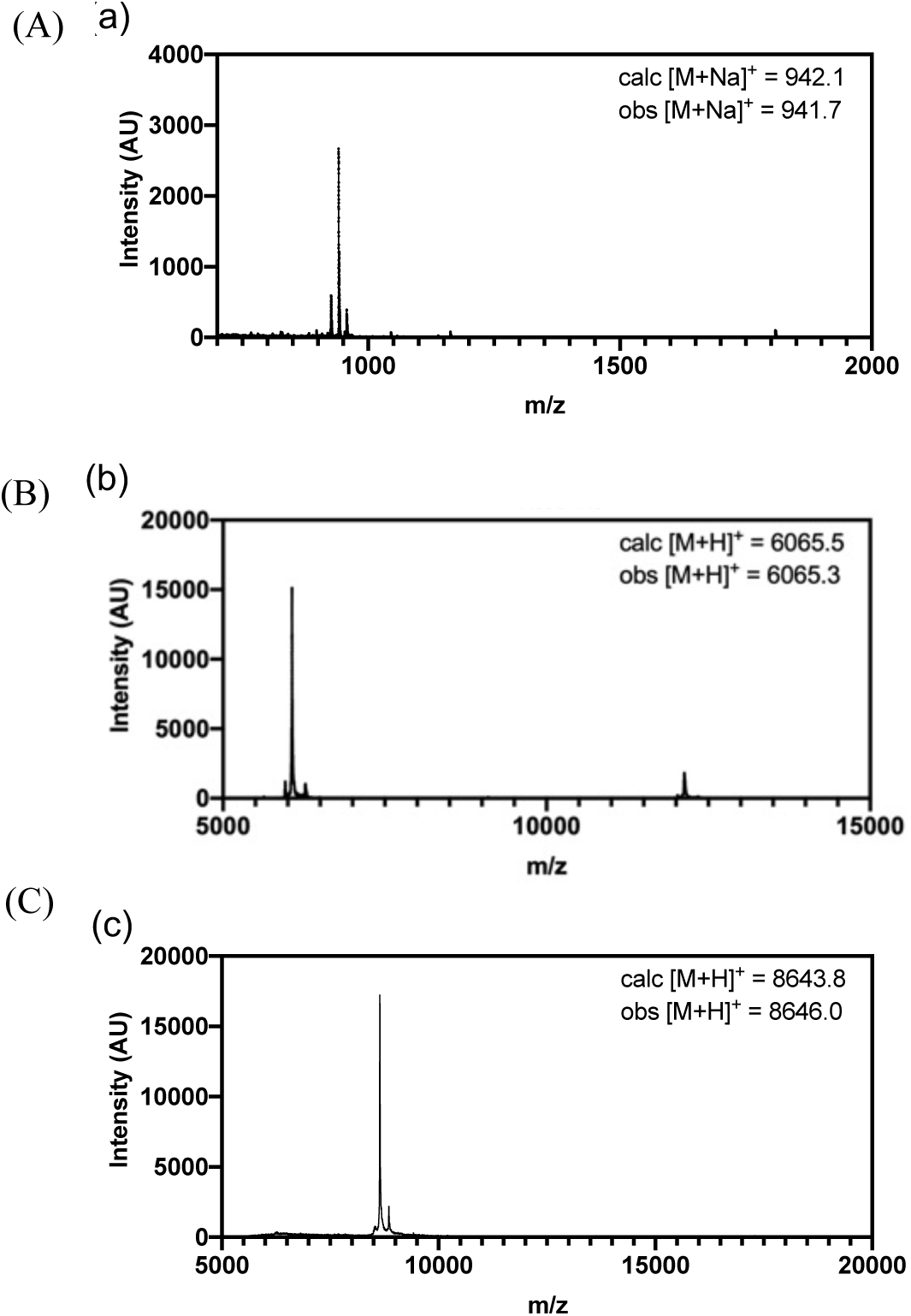
Protein semi-synthesis to incorporate ^Ac^K at position 80. MALDI-TOF-MS characterization of (A) αS_77-84_-Pen_77_^Ac^K_80_-NHNH_2_ (**2**), (B) αS_85-140_-C_85_ (**4**) and (C) αS_1-84_-Pen_77_^Ac^K_80_-MES (**3b**).

**Figure 3.**
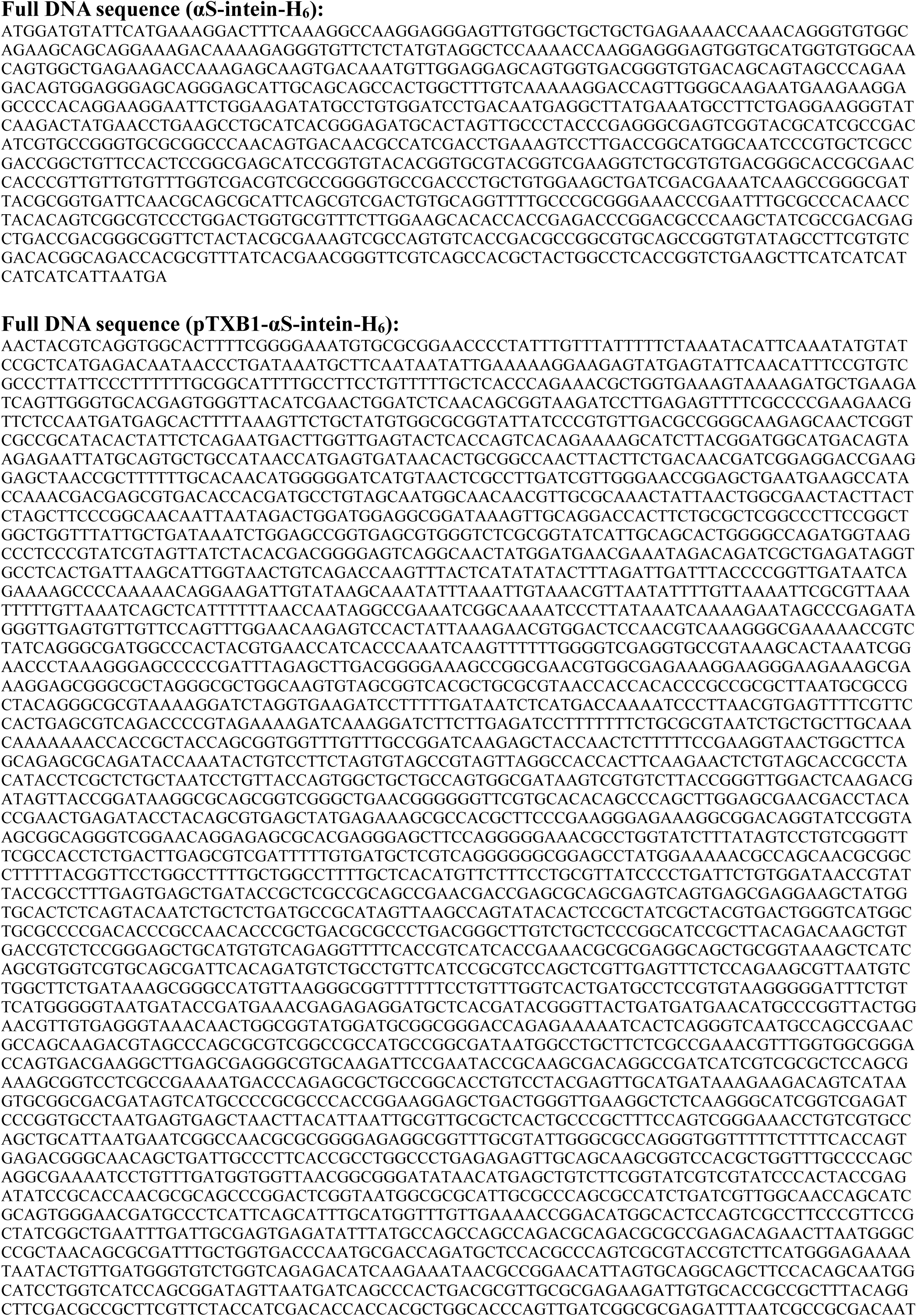
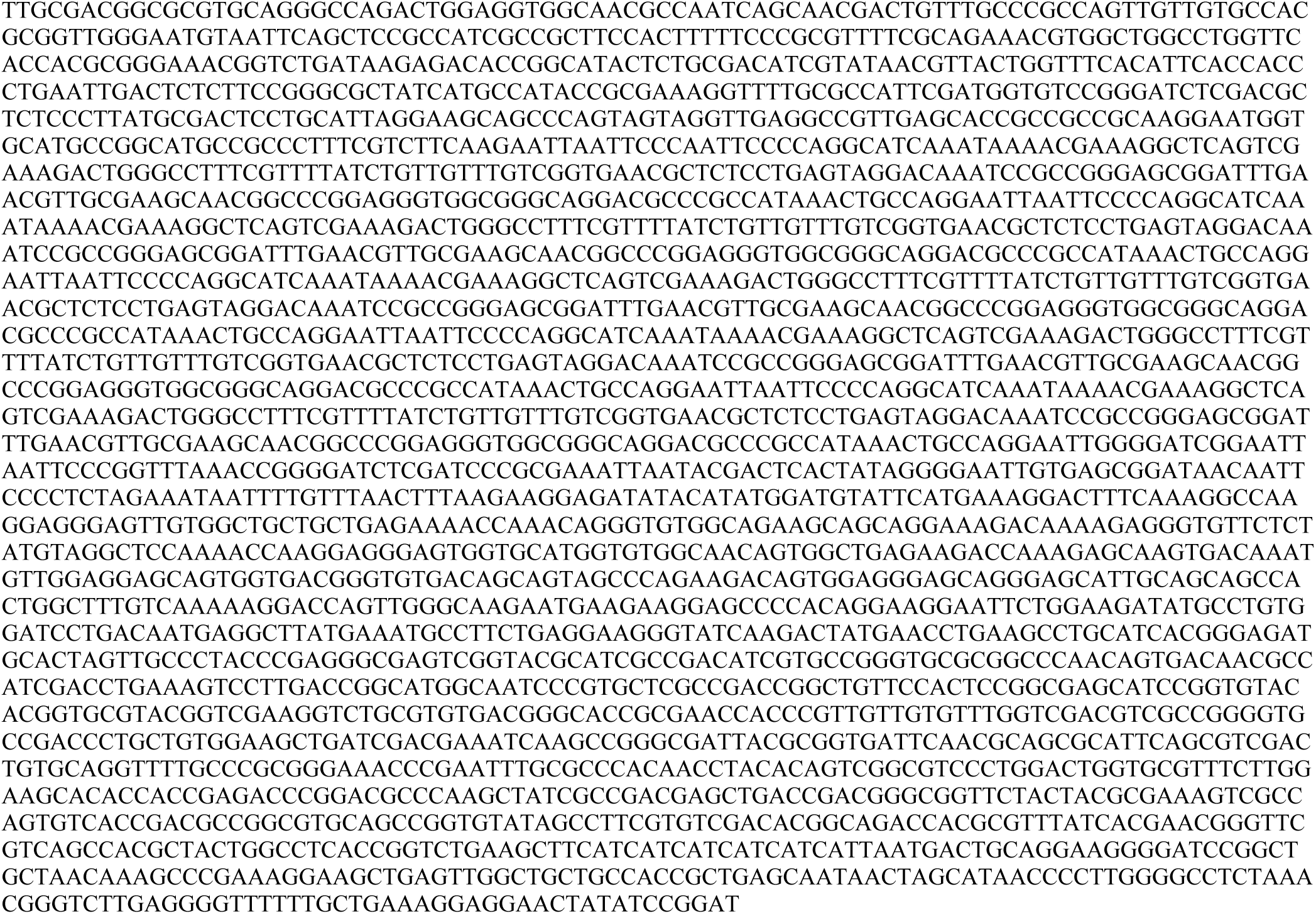
DNA sequences of WT αS expression plasmid.

**Figure 4.**
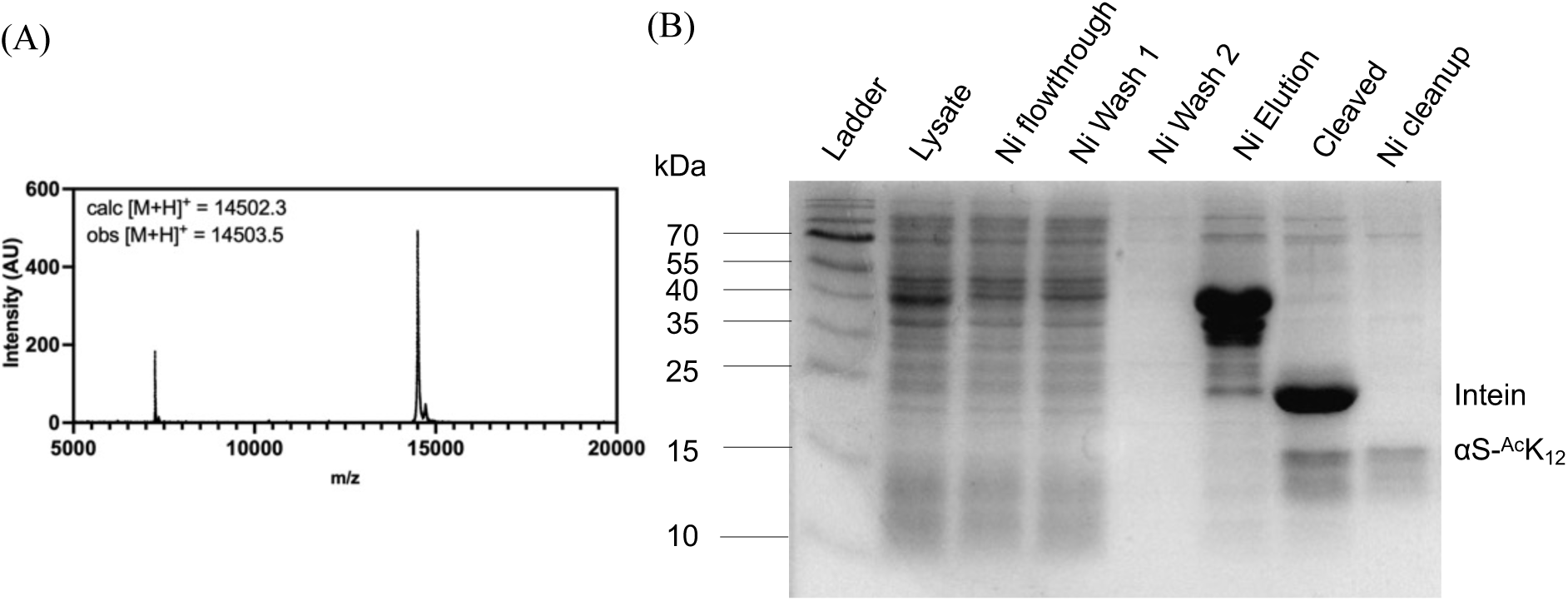
ncAA mutagenesis to incorporate acetyllysine at position 12. (A) MALDI-MS of purified product (B) SDS-PAGE with Coomassie staining to show affinity purification

**Figure 5.**
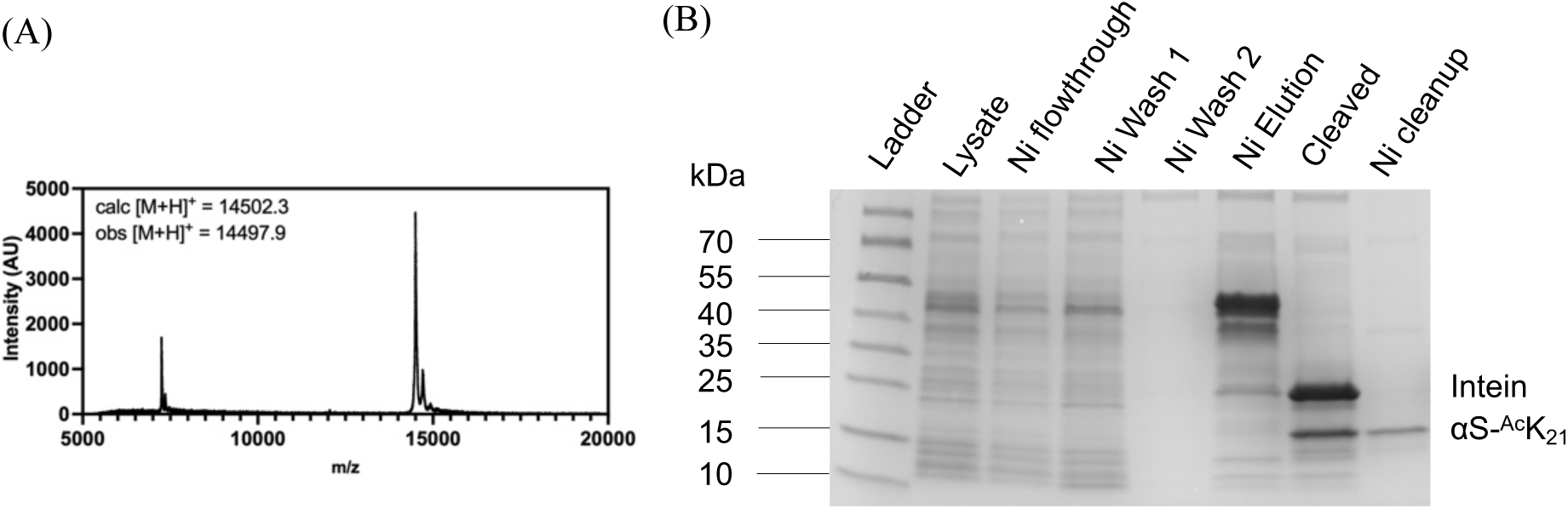
ncAA mutagenesis to incorporate acetyllysine at position 21. (A) MALDI-MS of purified product (B) SDS-PAGE with Coomassie staining to show affinity purification

**Figure 6.**
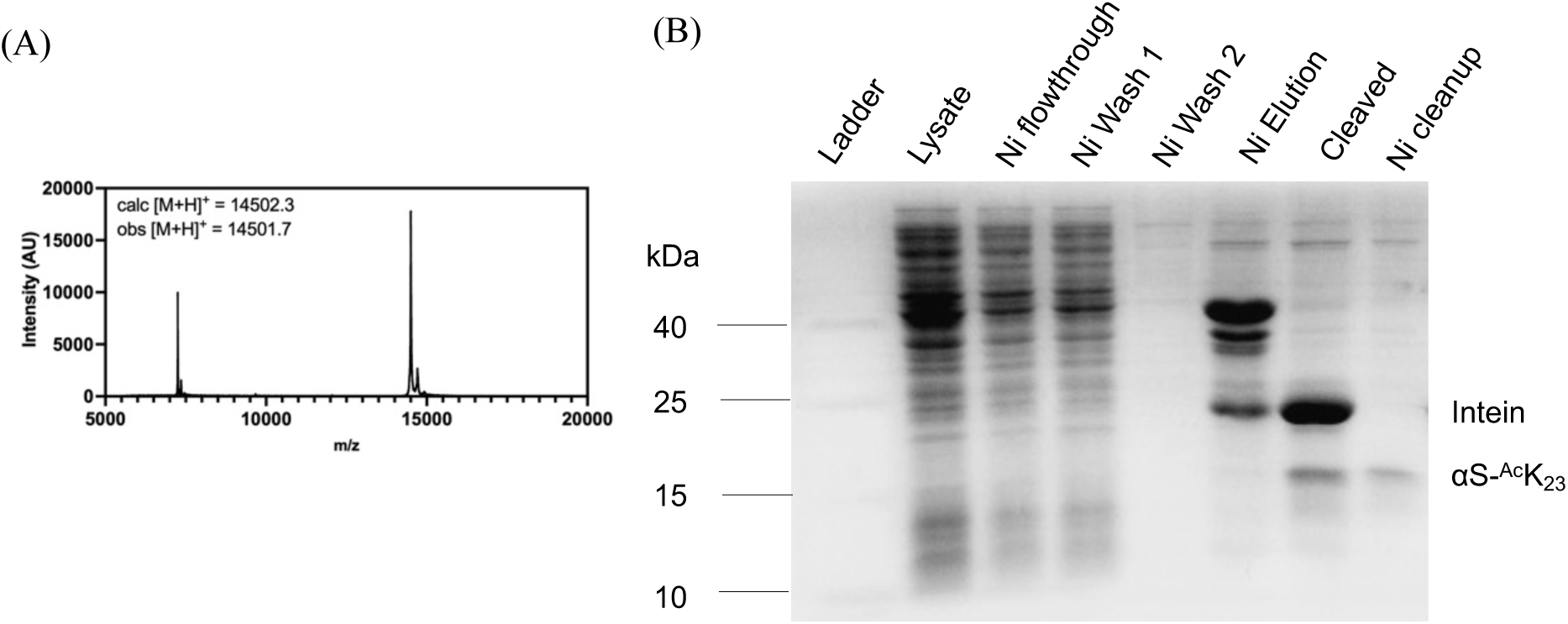
ncAA mutagenesis to incorporate acetyllysine at position 23. (A) MALDI-MS of purified product (B) SDS-PAGE with Coomassie staining to show affinity purification

**Figure 7.**
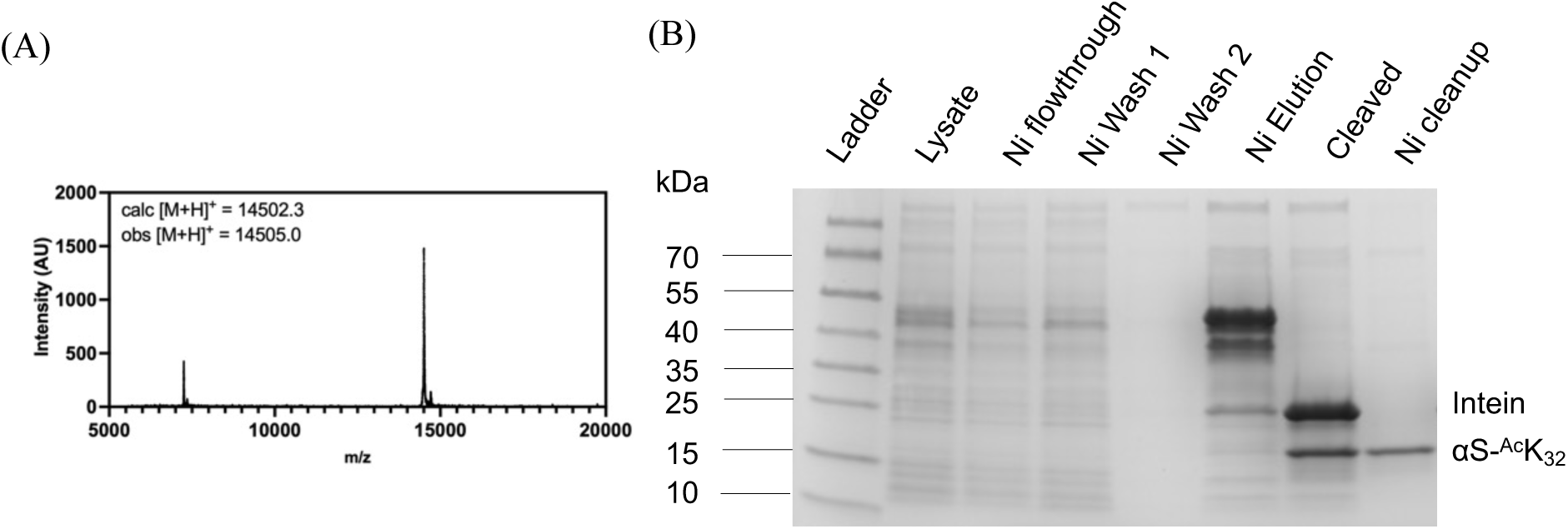
ncAA mutagenesis to incorporate acetyllysine at position 32. (A) MALDI-MS of purified product (B) SDS-PAGE with Coomassie staining to show affinity purification

**Figure 8.**
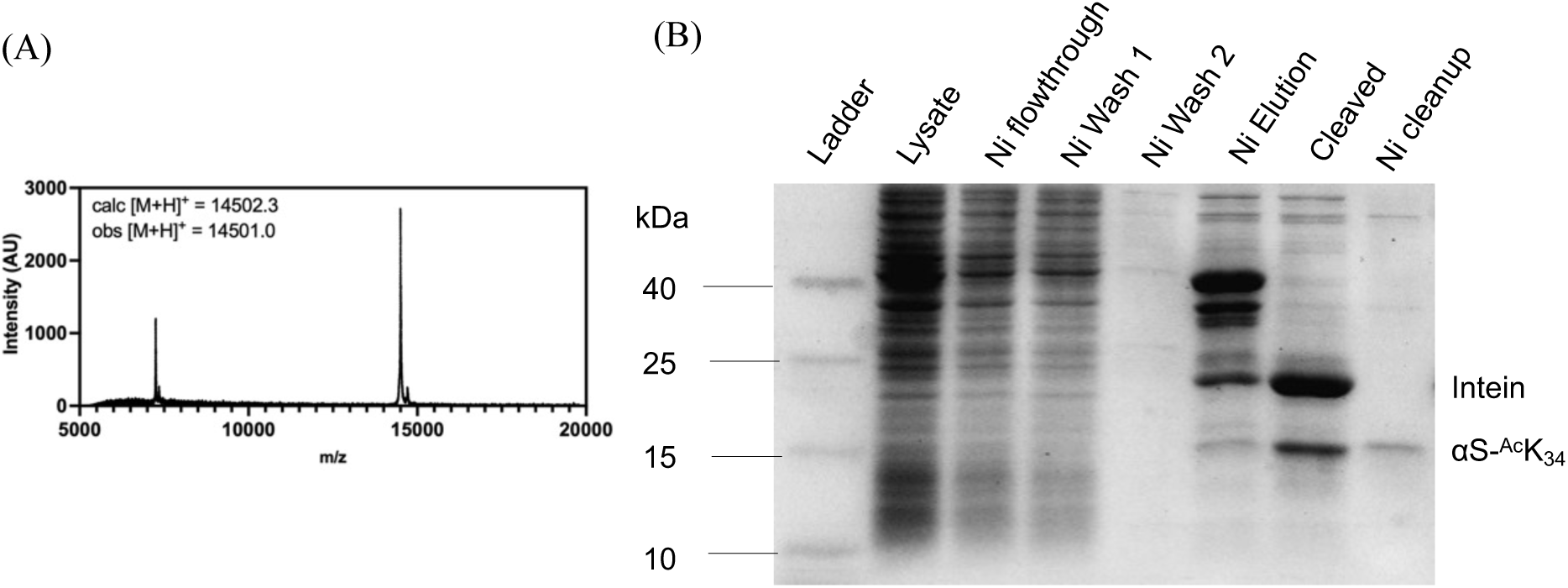
ncAA mutagenesis to incorporate acetyllysine at position 34. (A) MALDI-MS of purified product (B) SDS-PAGE with Coomassie staining to show affinity purification

**Figure 9.**
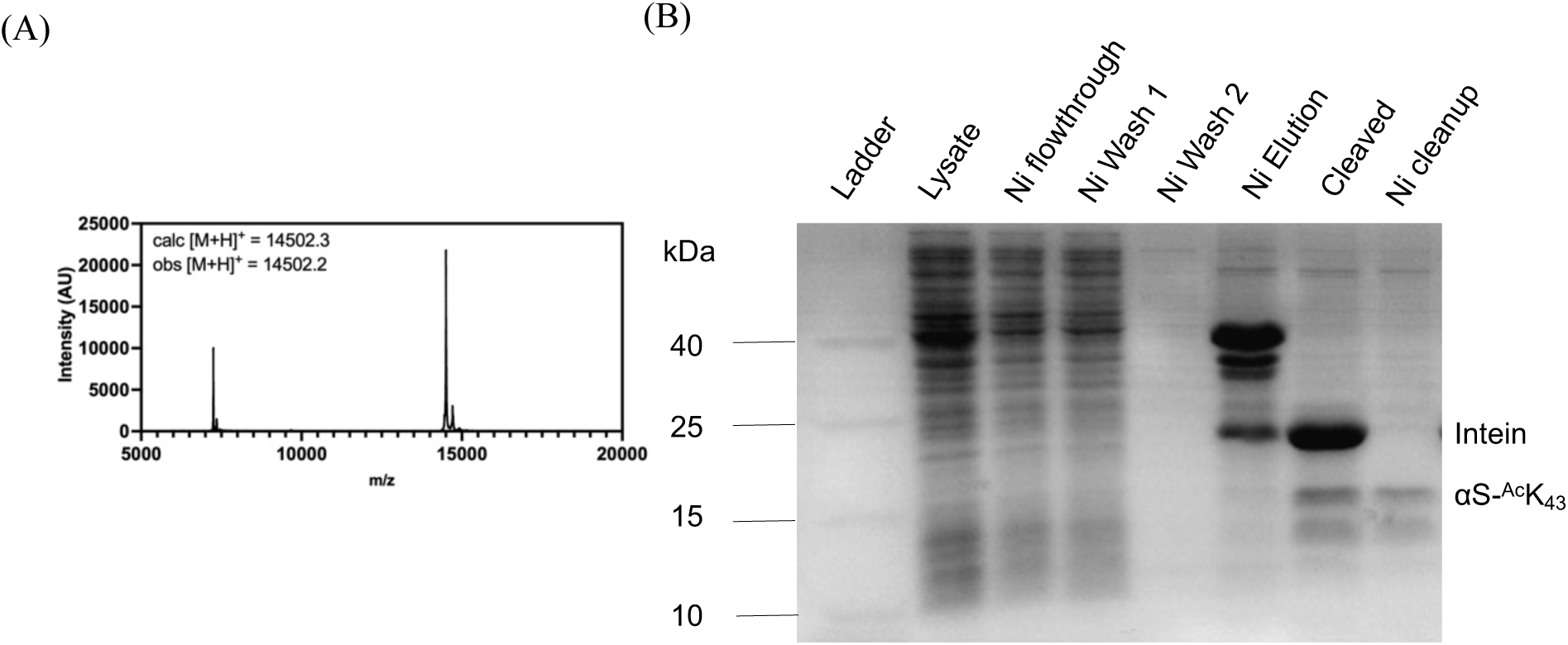
ncAA mutagenesis to incorporate acetyllysine at position 43. (A) MALDI-MS of purified product (B) SDS-PAGE with Coomassie staining to show affinity purification

**Figure 10.**
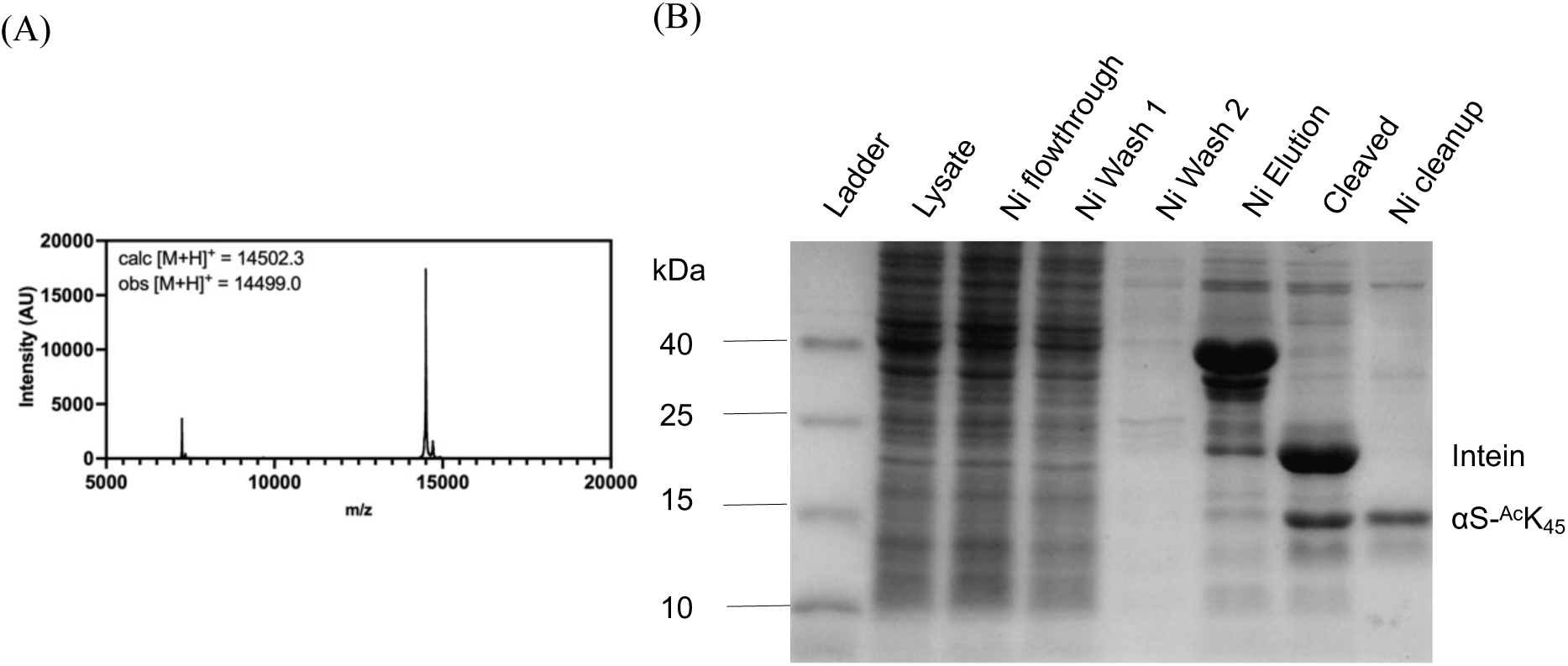
ncAA mutagenesis to incorporate acetyllysine at position 45. (A) MALDI-MS of purified product (B) SDS-PAGE with Coomassie staining to show affinity purification

**Figure 11.**
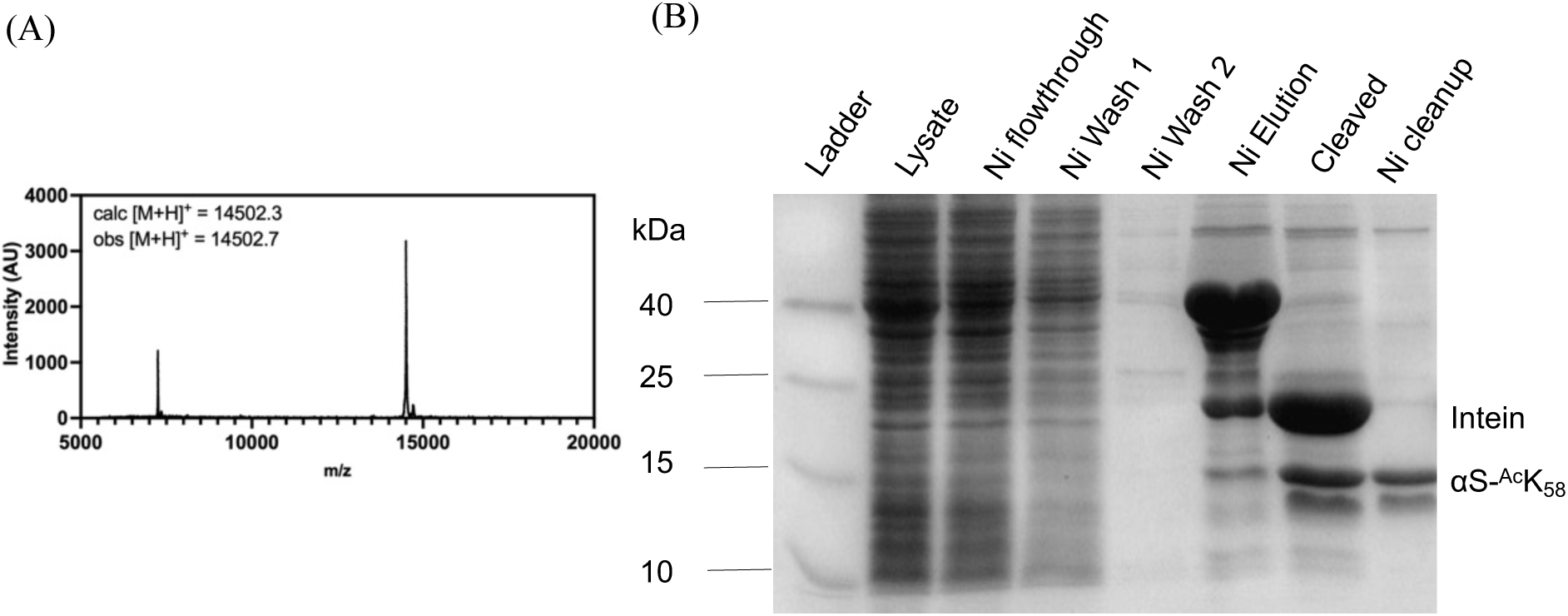
ncAA mutagenesis to incorporate acetyllysine at position 58. (A) MALDI-MS of purified product (B) SDS-PAGE with Coomassie staining to show affinity purification

**Figure 12.**
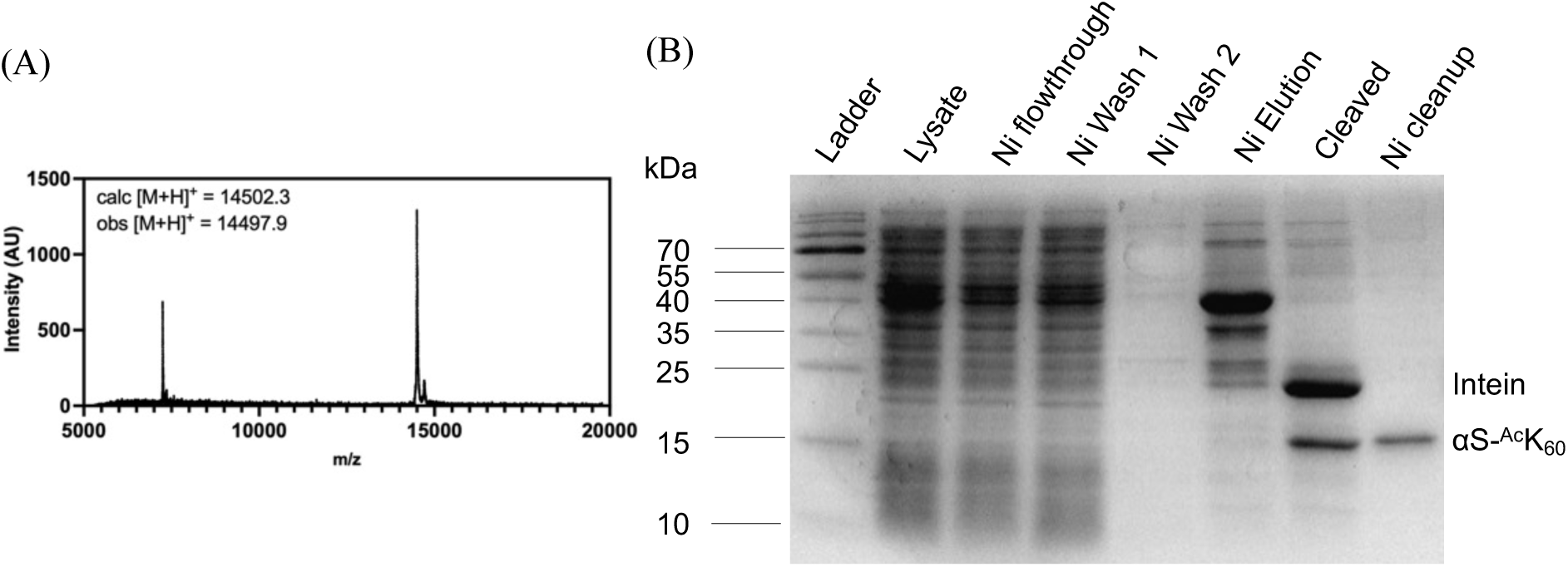
ncAA mutagenesis to incorporate acetyllysine at position 60. (A) MALDI-MS of purified product (B) SDS-PAGE with Coomassie staining to show affinity purification

**Figure 13.**
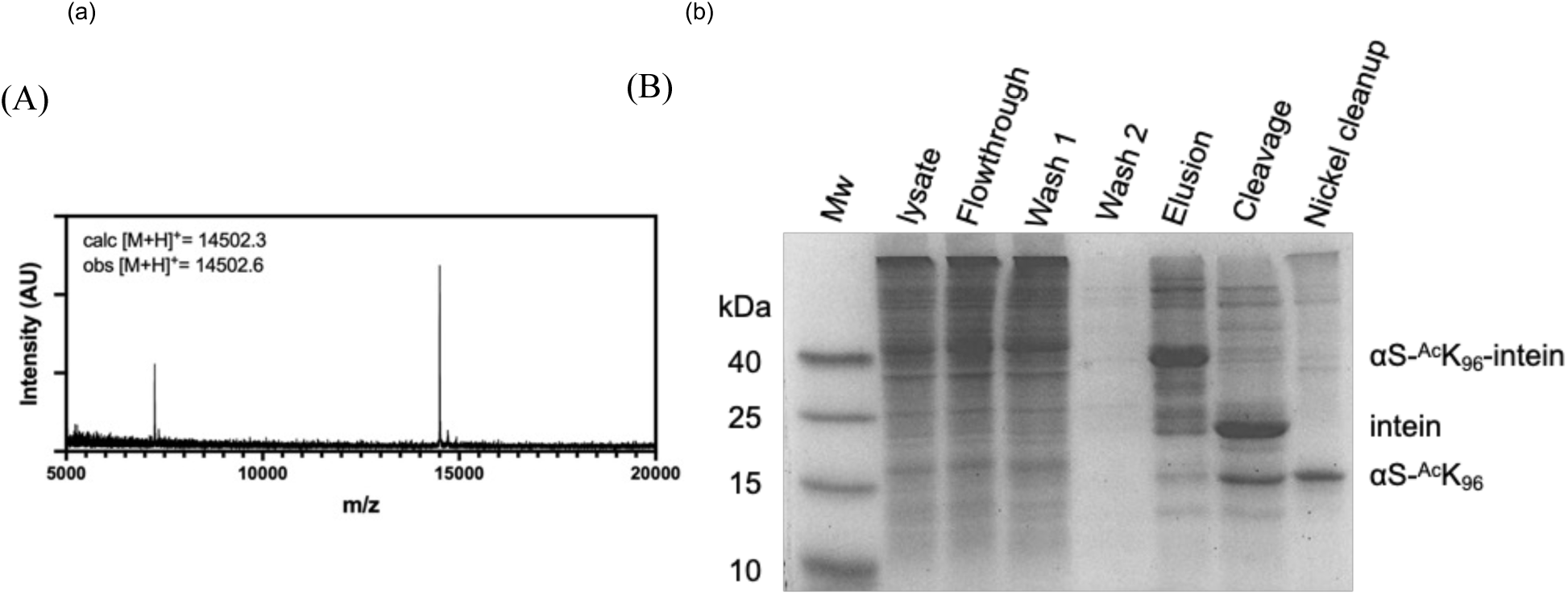
ncAA mutagenesis to incorporate acetyllysine at position 96. (A) MALDI-MS of purified product (B) SDS-PAGE with Coomassie staining to show affinity purification

**Figure 14.**
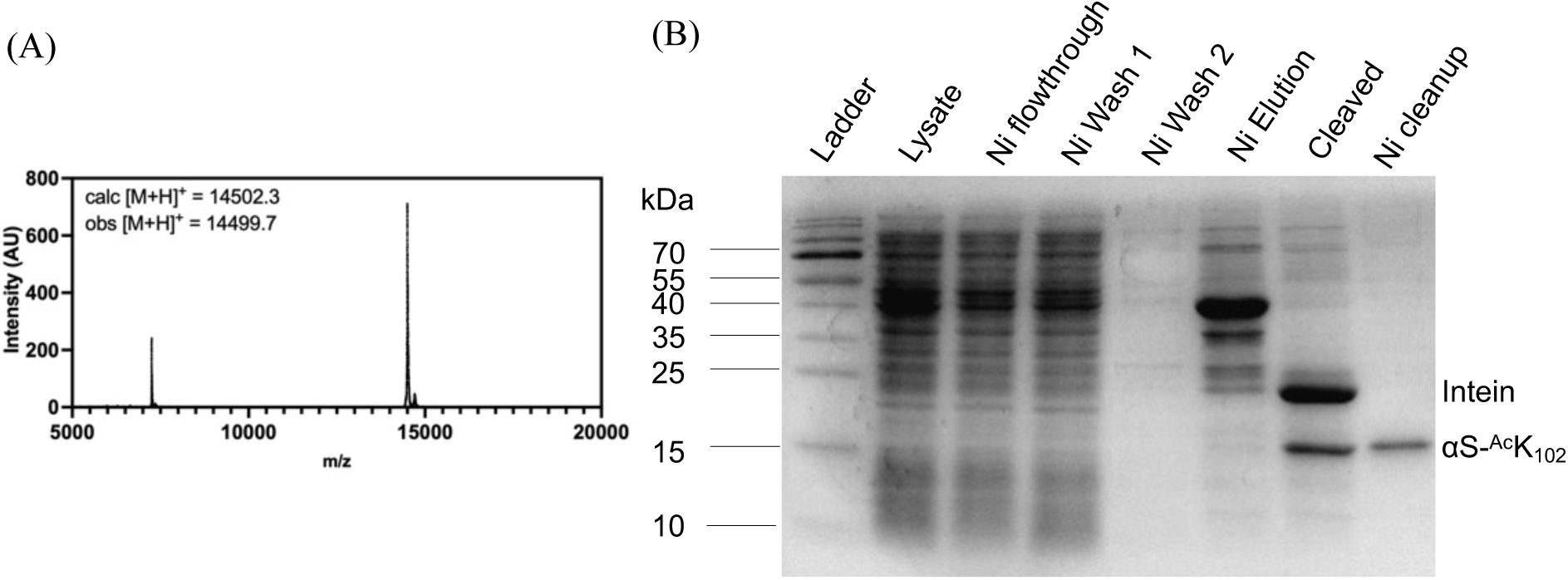
ncAA mutagenesis to incorporate acetyllysine at position 102. (A) MALDI-MS of purified product (B) SDS-PAGE with Coomassie staining to show affinity purification

**Figure 15.**
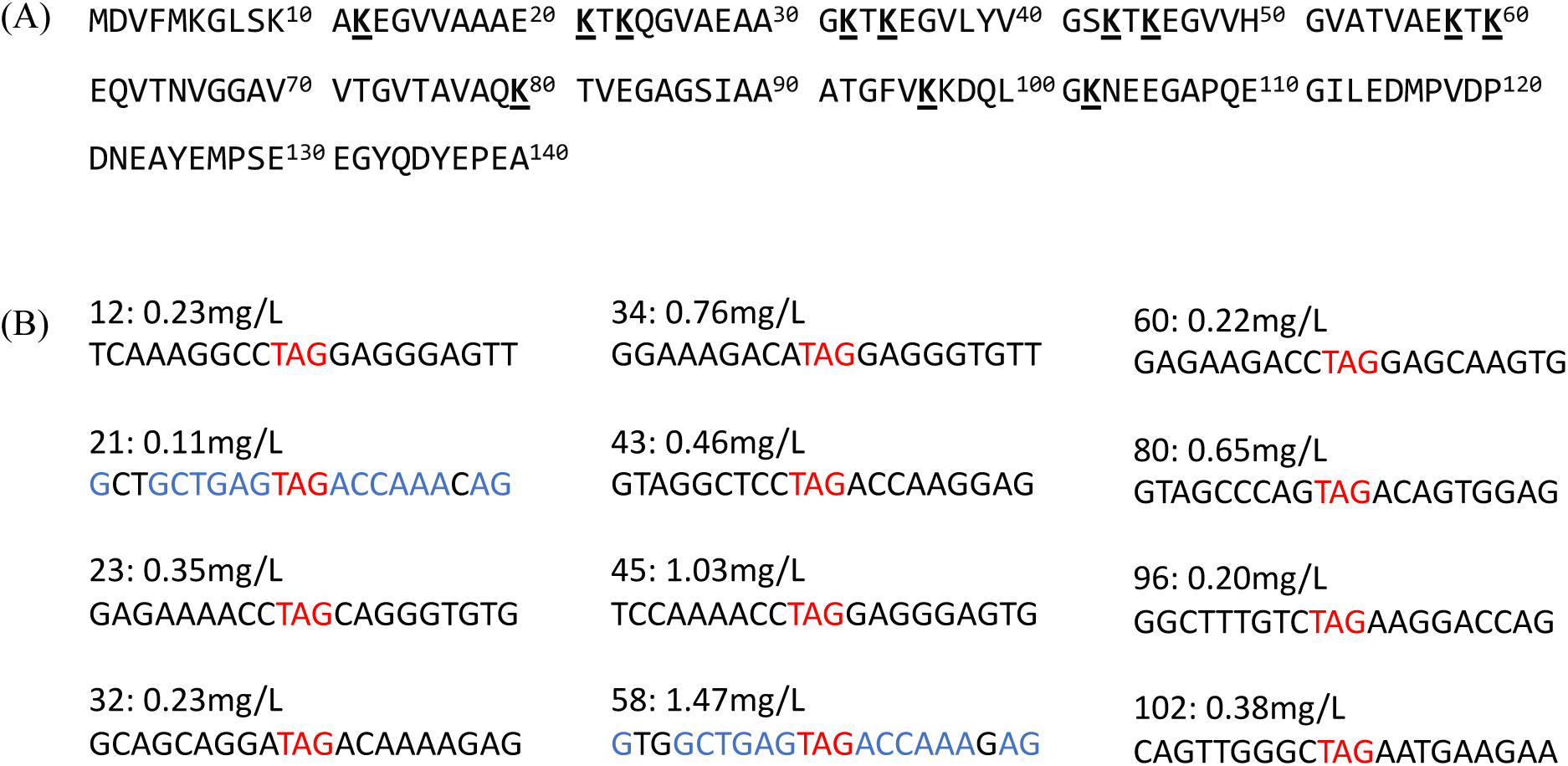
Local sequential contexts of amber codon suppression. (A) Amino acid sequence of αS with acetyl lysine incorporation sites bolded and underlined, (B) Expression yield per liter of *E. coli* culture for each acetylated construct. Sites 21 and 58 have very similar local sequence context yet gave very different suppression yields.

**Figure 16a.**
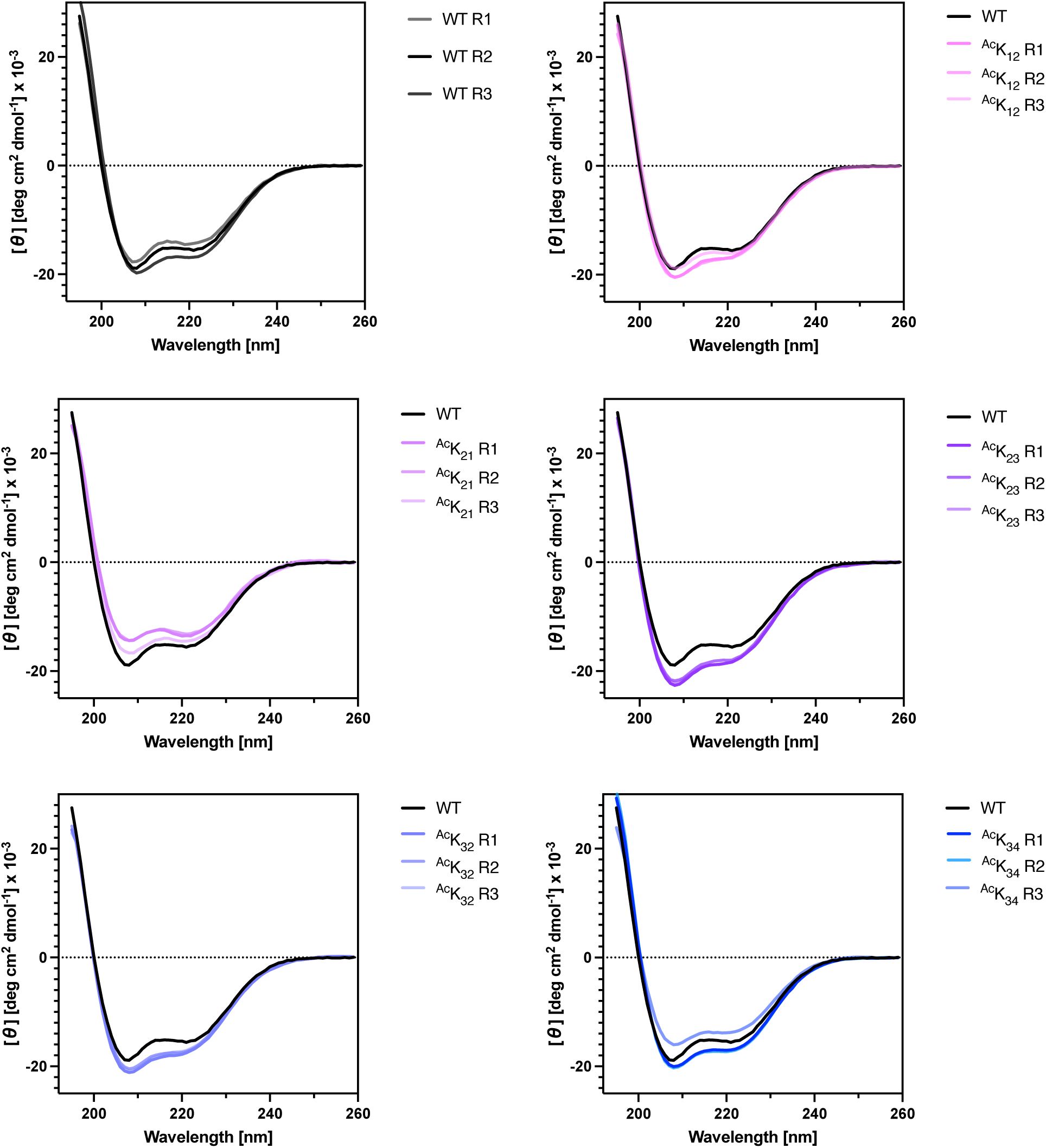
Individual CD wavelength scans for αS-^Ac^K constructs.

**Figure 16b.**
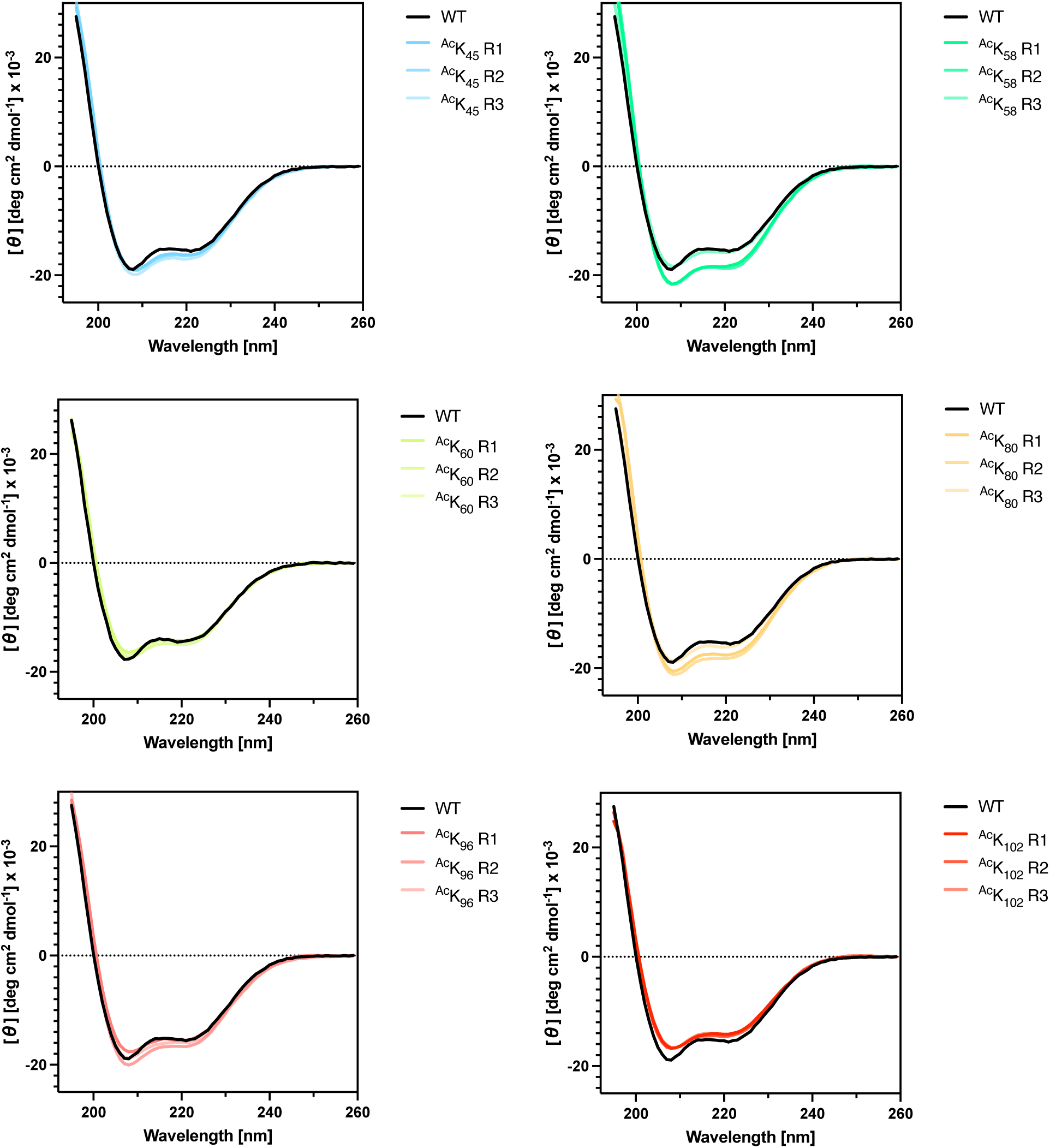
Individual CD wavelength scans for αS-^Ac^K constructs.

**Figure 17a.**
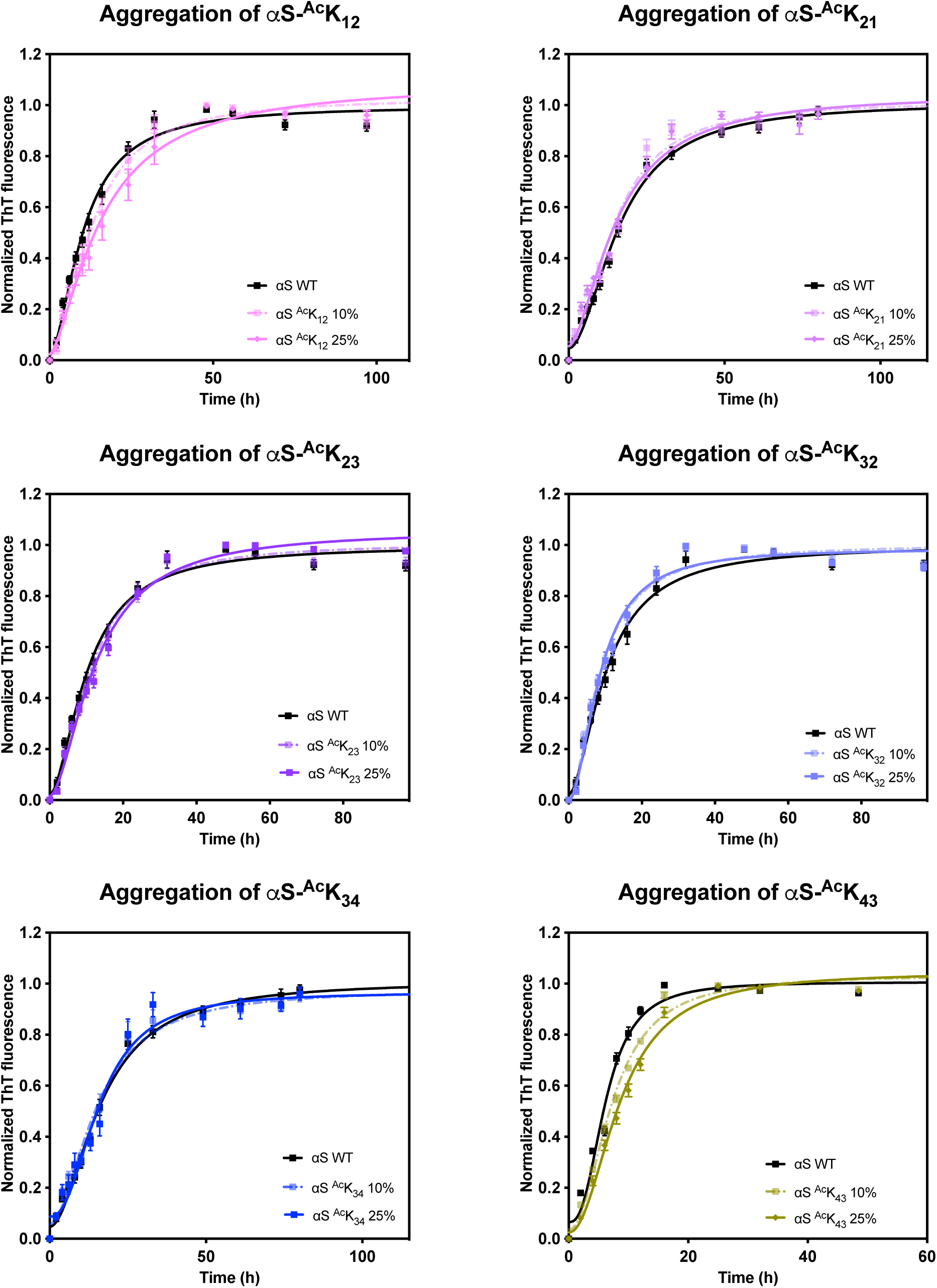
Aggregation kinetics curves for each αS-^Ac^K construct.

**Figure 17b.**
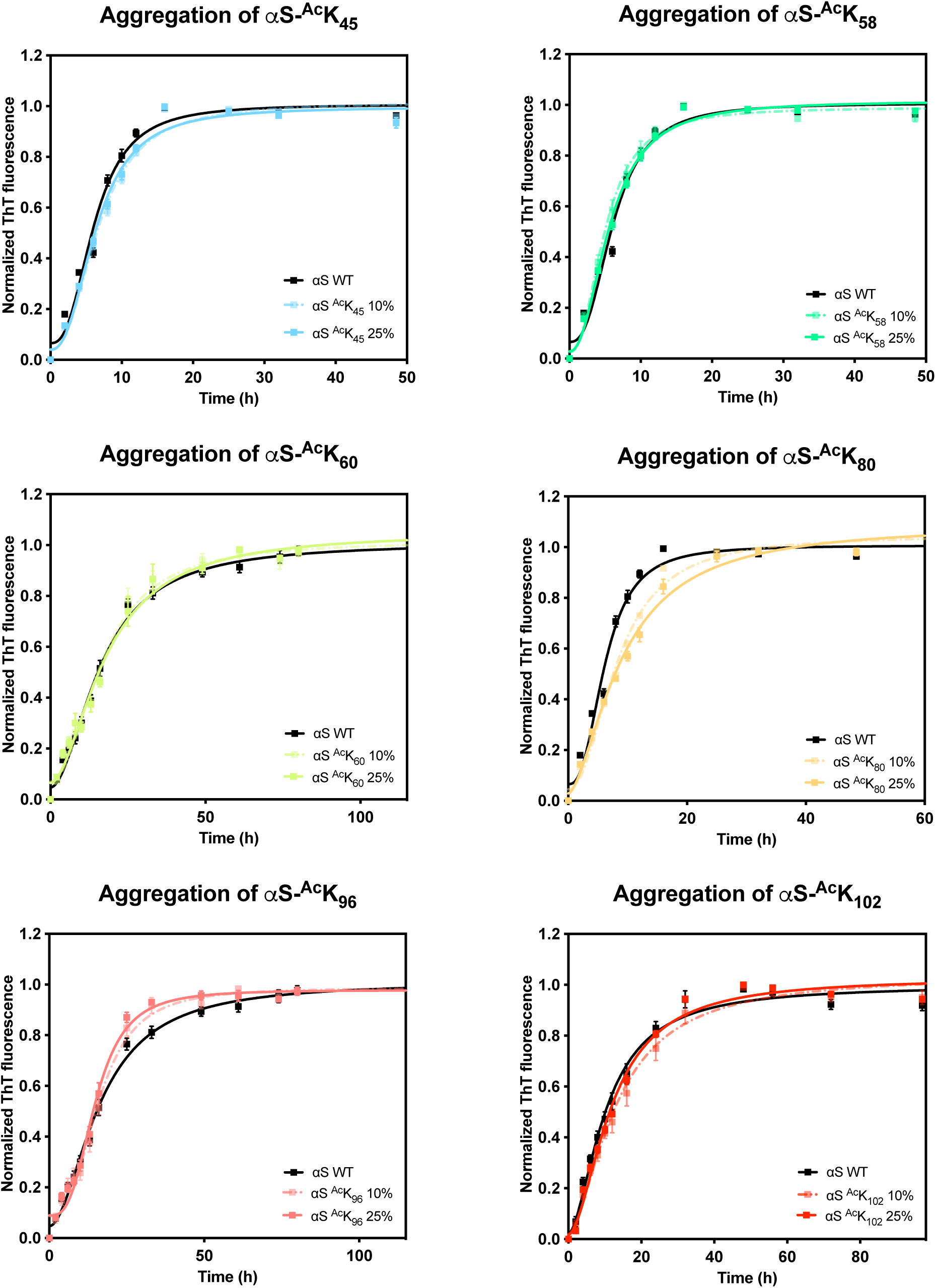
Aggregation kinetics curves for each αS-^Ac^K construct.

**Figure 18.**
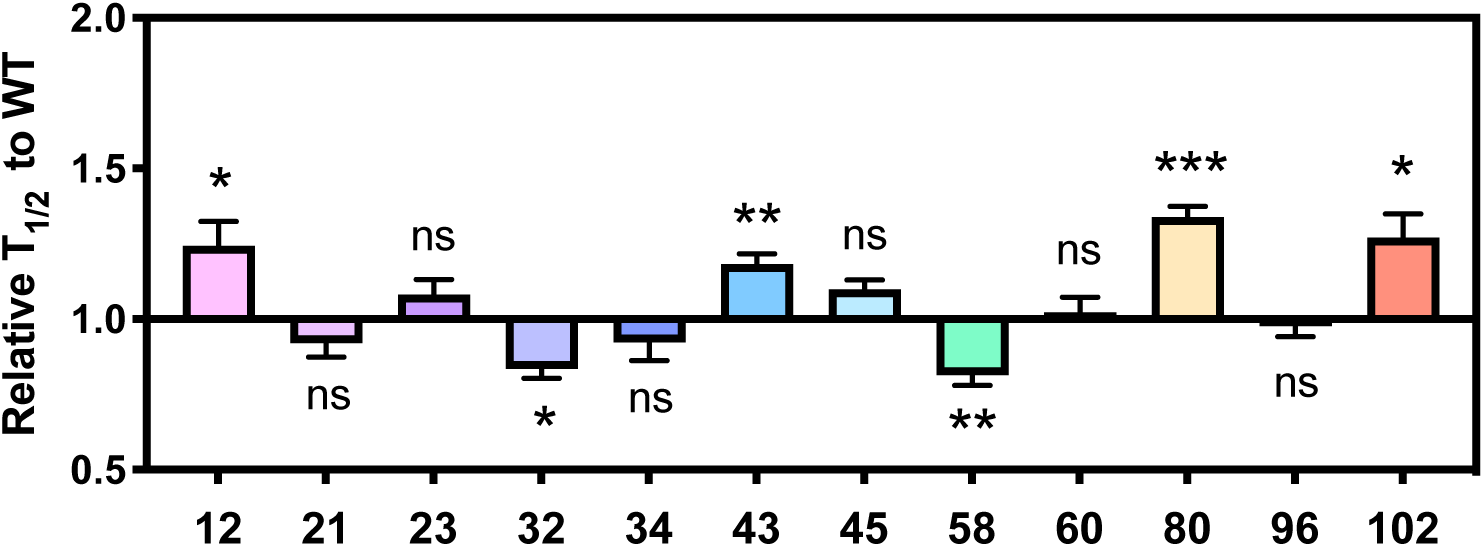
Effects of 10% αS-^Ac^K on aggregation kinetics. The time to reach 50% fibrilization (T_1/2_) for each condition was normalized to that of a 100% WT aggregation. Seeded aggregation was performed with αS monomers where acetylated αS was mixed with αS WT at 10%:90% ratio. Mean, with standard error of six replicates.

**Figure 19a.**
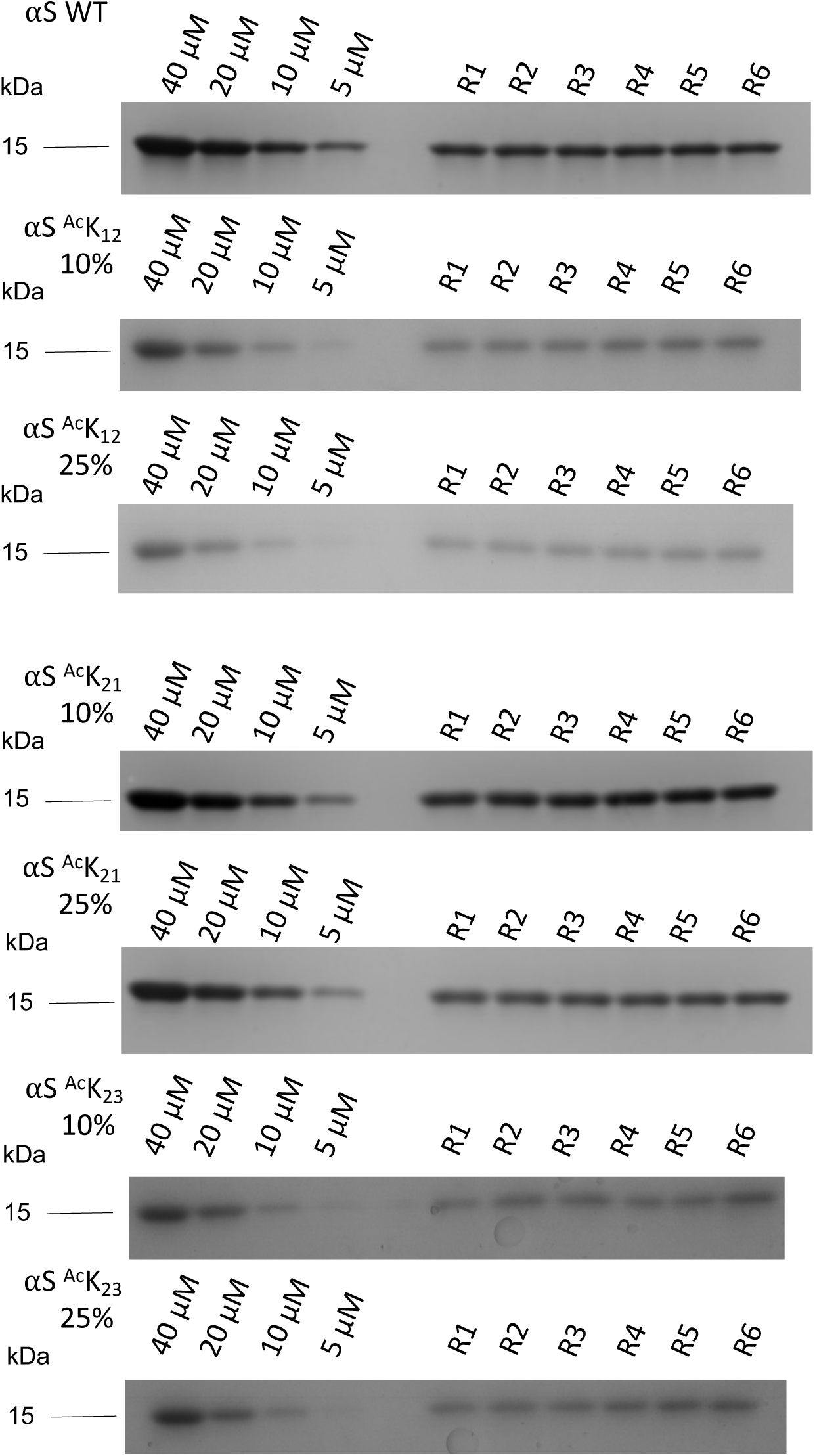
Primary SDS-PAGE gels for quantifying monomer incorporations of αS-^Ac^K constructs. Six replicates shown (R1-6).

**Figure 19b.**
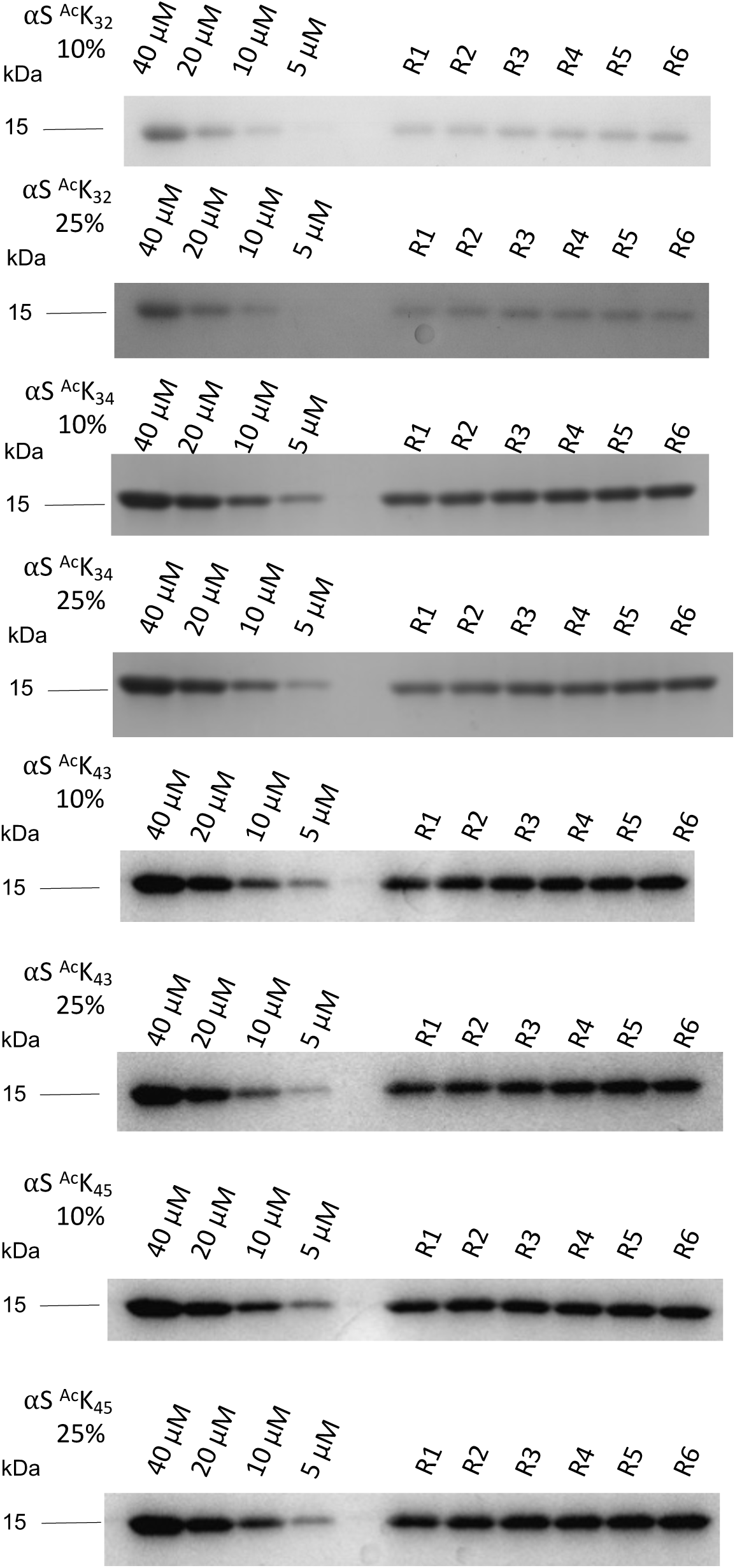
Primary SDS-PAGE gels for quantifying monomer incorporations of αS-^Ac^K constructs. Six replicates shown (R1-6)

**Figure 19c.**
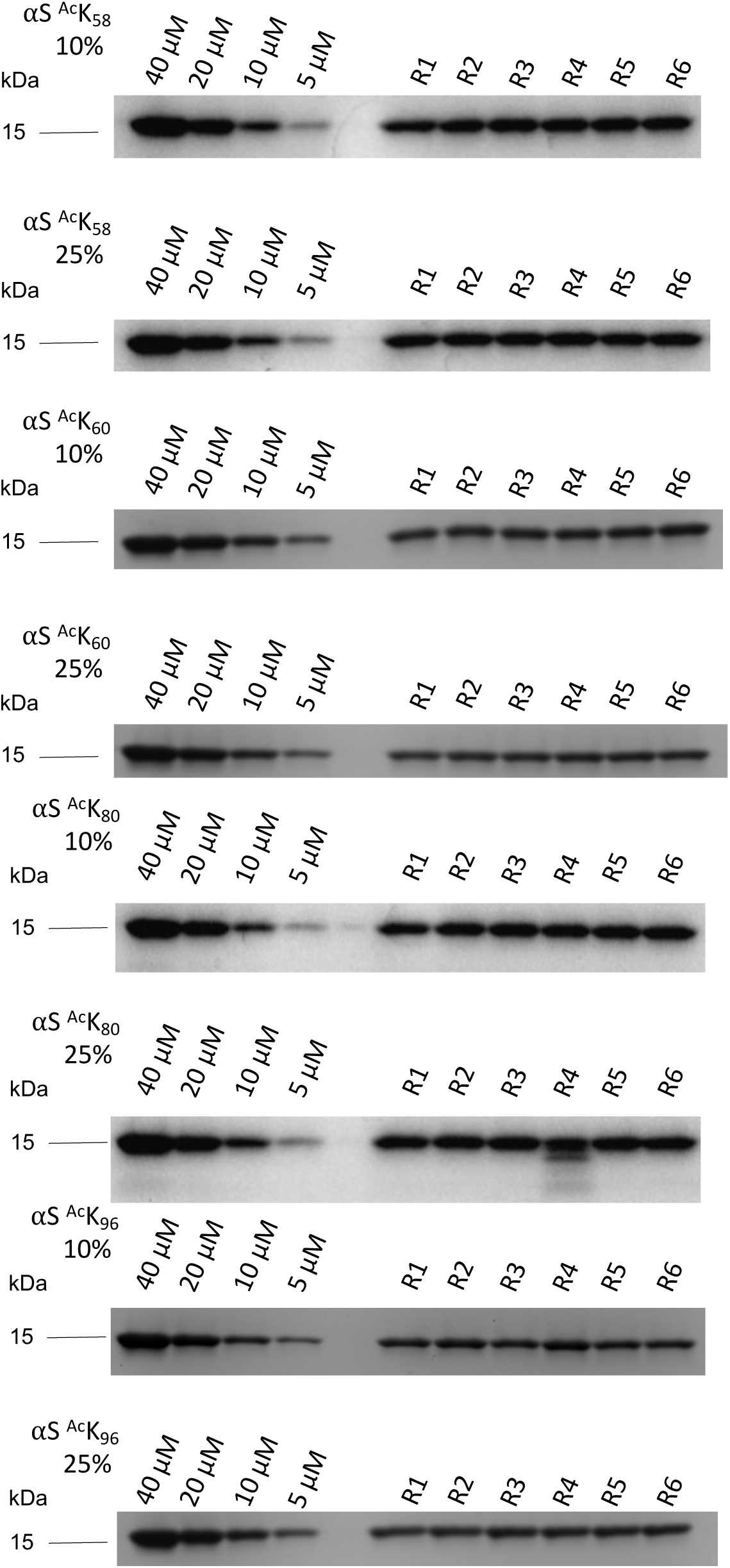
Primary SDS-PAGE gels for quantifying monomer incorporations of αS-^Ac^K constructs. Six replicates shown (R1-6).

**Figure 19d.**
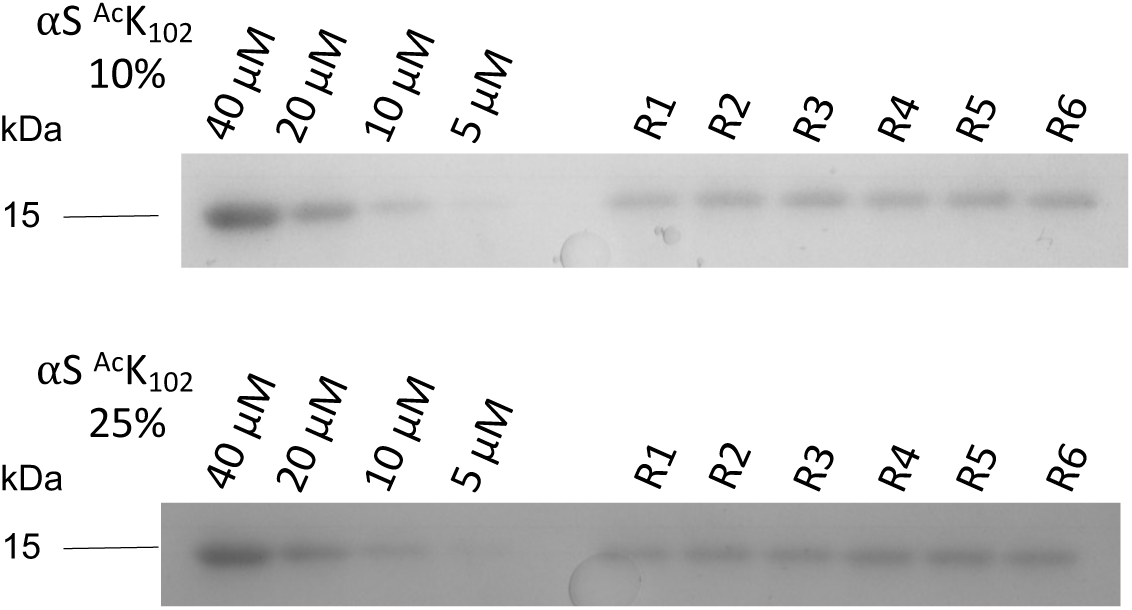
Primary SDS-PAGE gels for quantifying monomer incorporations of αS-^Ac^K constructs. Six replicates shown (R1-6).

**Figure 20.**
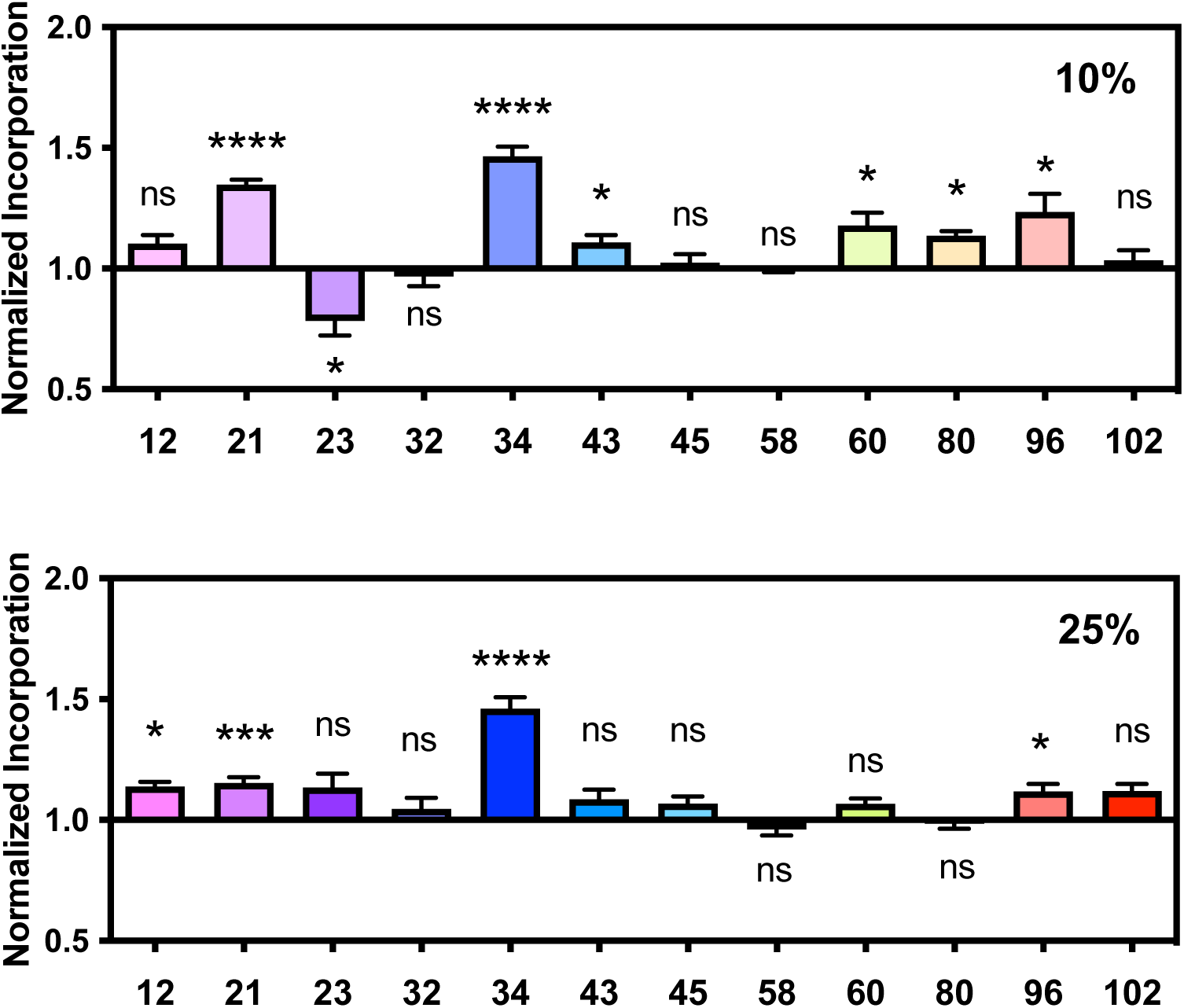
Effects of αS-^Ac^K on total monomer incorporation. Monomers incorporated into fibrils were quantified by SDS-PAGE gels and normalized to WT values. Mean with standard error, R=6

**Figure 21.**
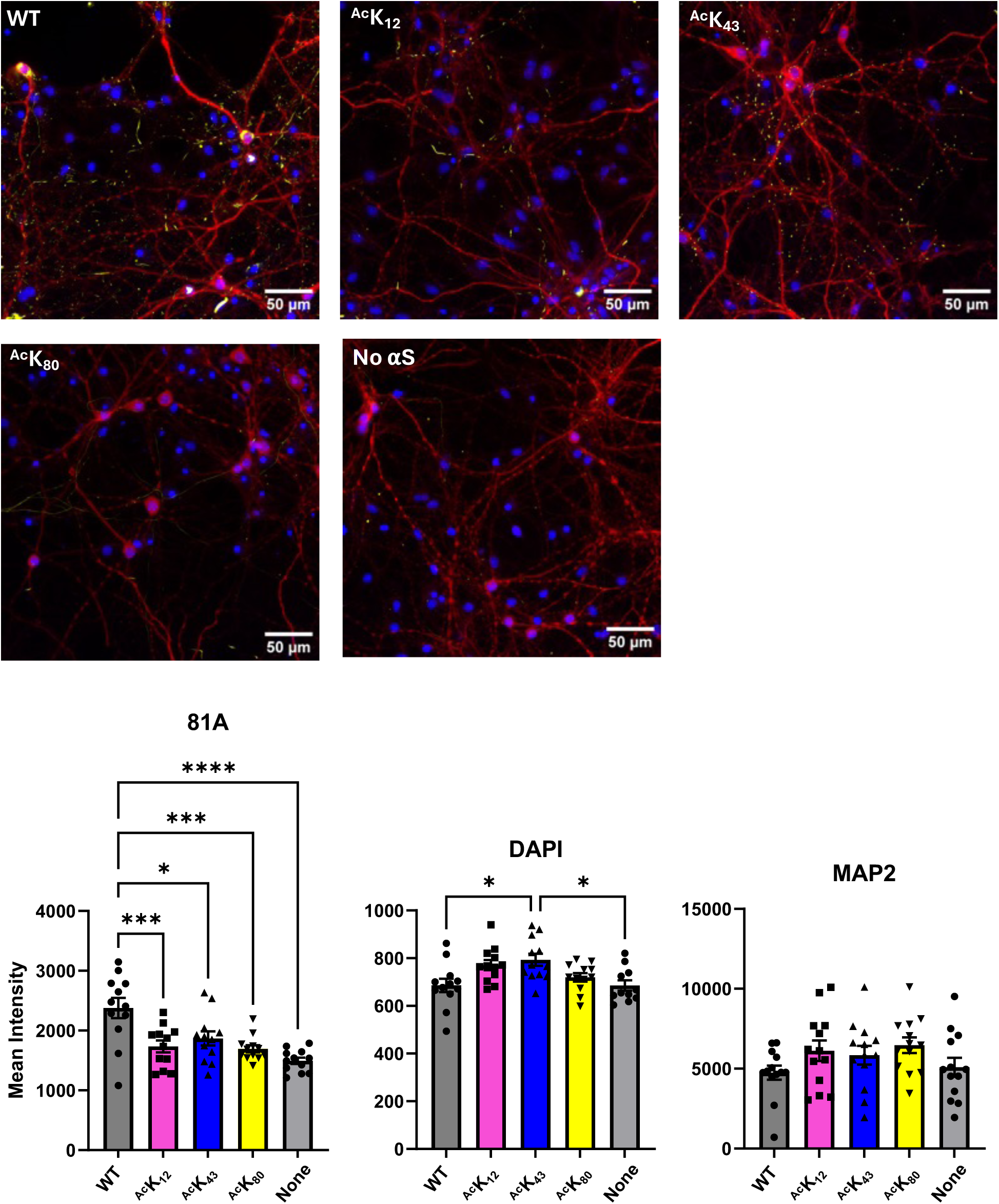
Neuron imaging data Top: Representative images of neuron cultures with additional stains. Yellow = 81A (anti-pS129), Blue = DAPI, Red = MAP2 (Larger fields of view shown than in main text). Bottom: Fluorescence intensity from 81A, DAPI, and MAP2 channels. * = 0.01 < p-value < 0.05; *** = 0.001 < p-value < 0.0001; **** = 0.00001 < p-value < 0.0001

**Figure 22.**
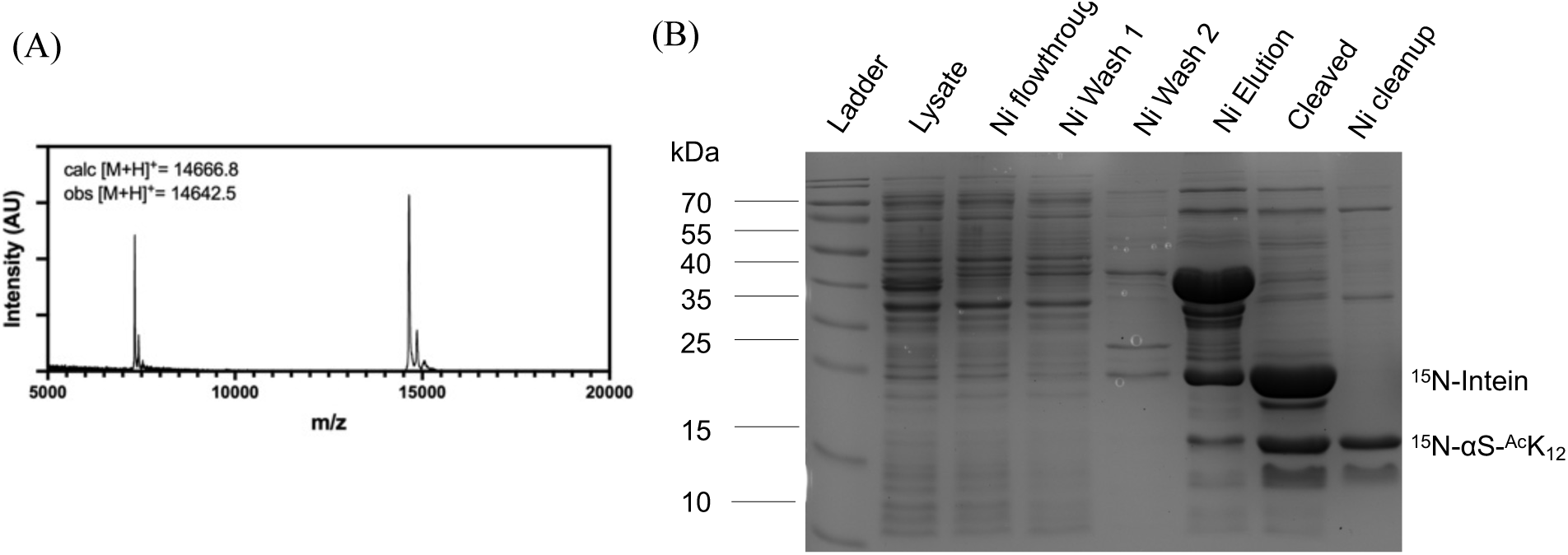
Recombinant ^15^N- αS-^Ac^K_12_ (A) MALDI-MS of purified product (B) SDS-PAGE with Coomassie staining to show affinity purification.

**Figure 23.**
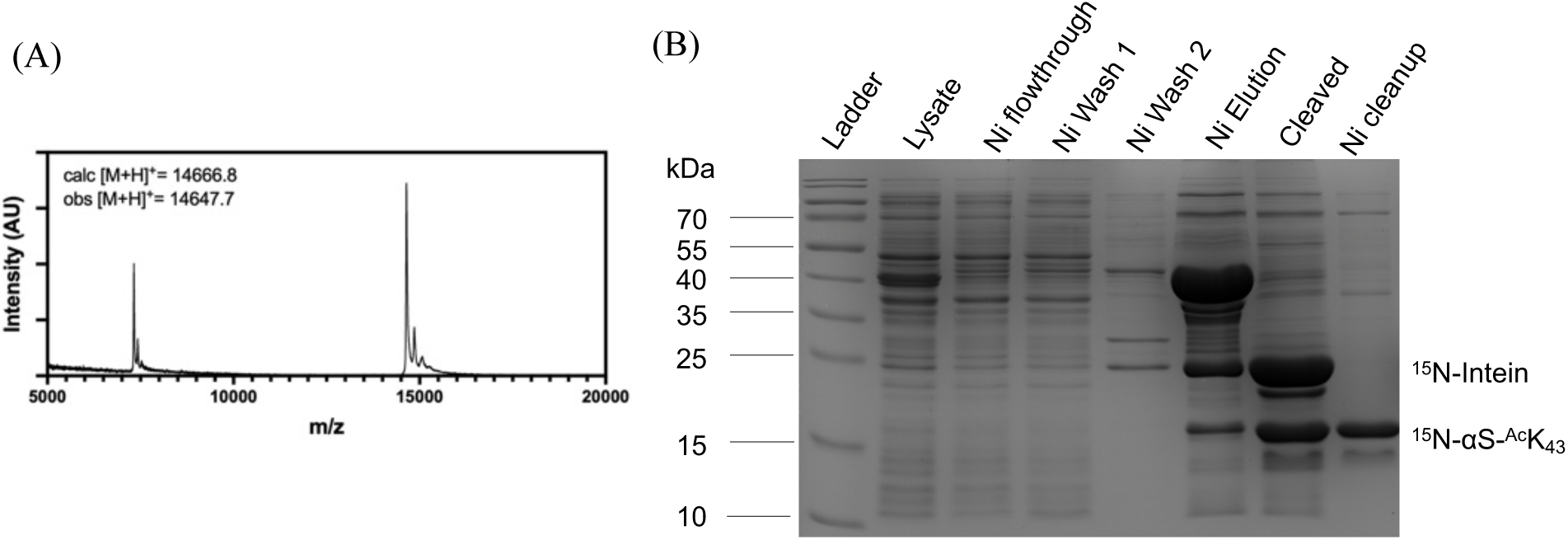
Recombinant ^15^N- αS-^Ac^K_43_ (A) MALDI-MS of purified product (B) SDS-PAGE with Coomassie staining to show affinity purification.

**Figure 24.**
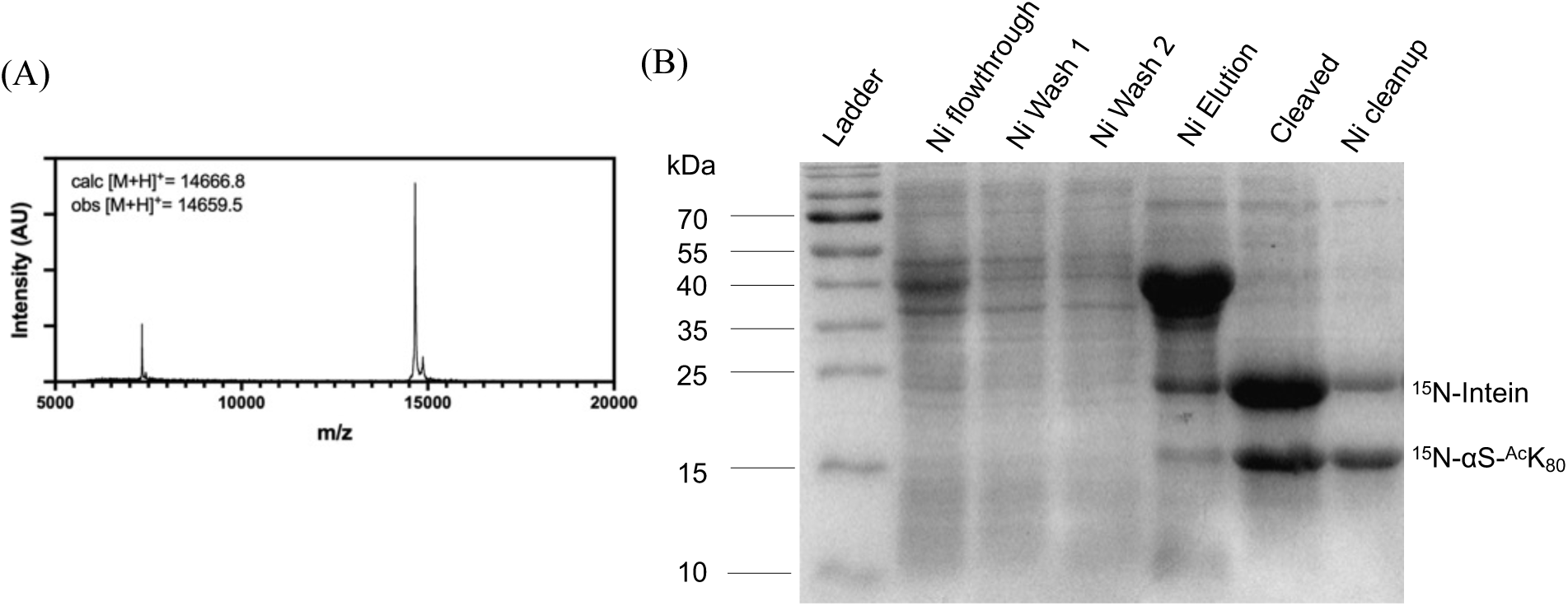
Recombinant ^15^N- αS-^Ac^K_80_ (A) MALDI-MS of purified product (B) SDS-PAGE with Coomassie staining to show affinity purification.

**Figure 25a.**
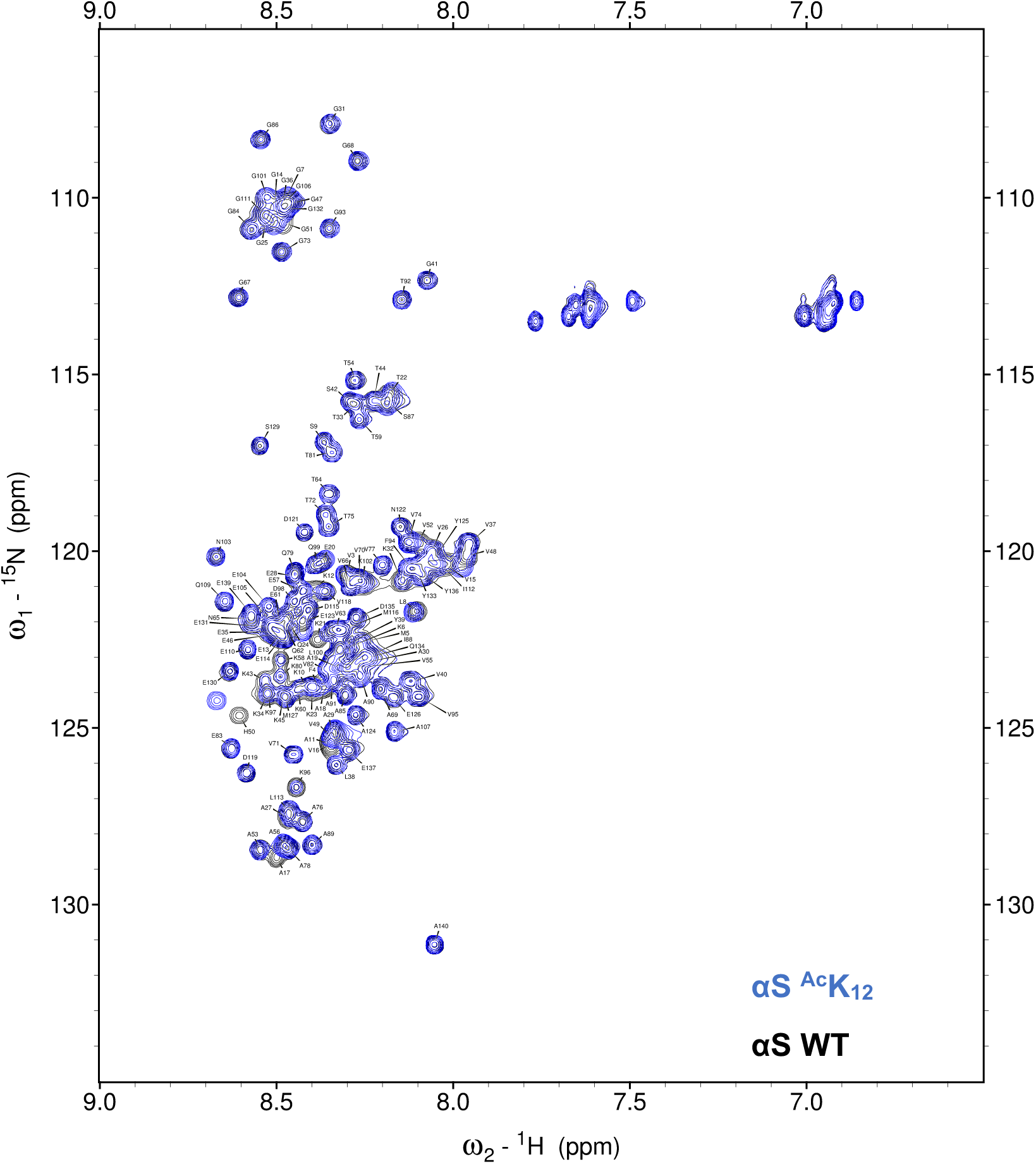
HSQC spectra acquired for αS-^Ac^K_12_ in buffer, overlayed with the spectra of αS WT.

**Figure 25b.**
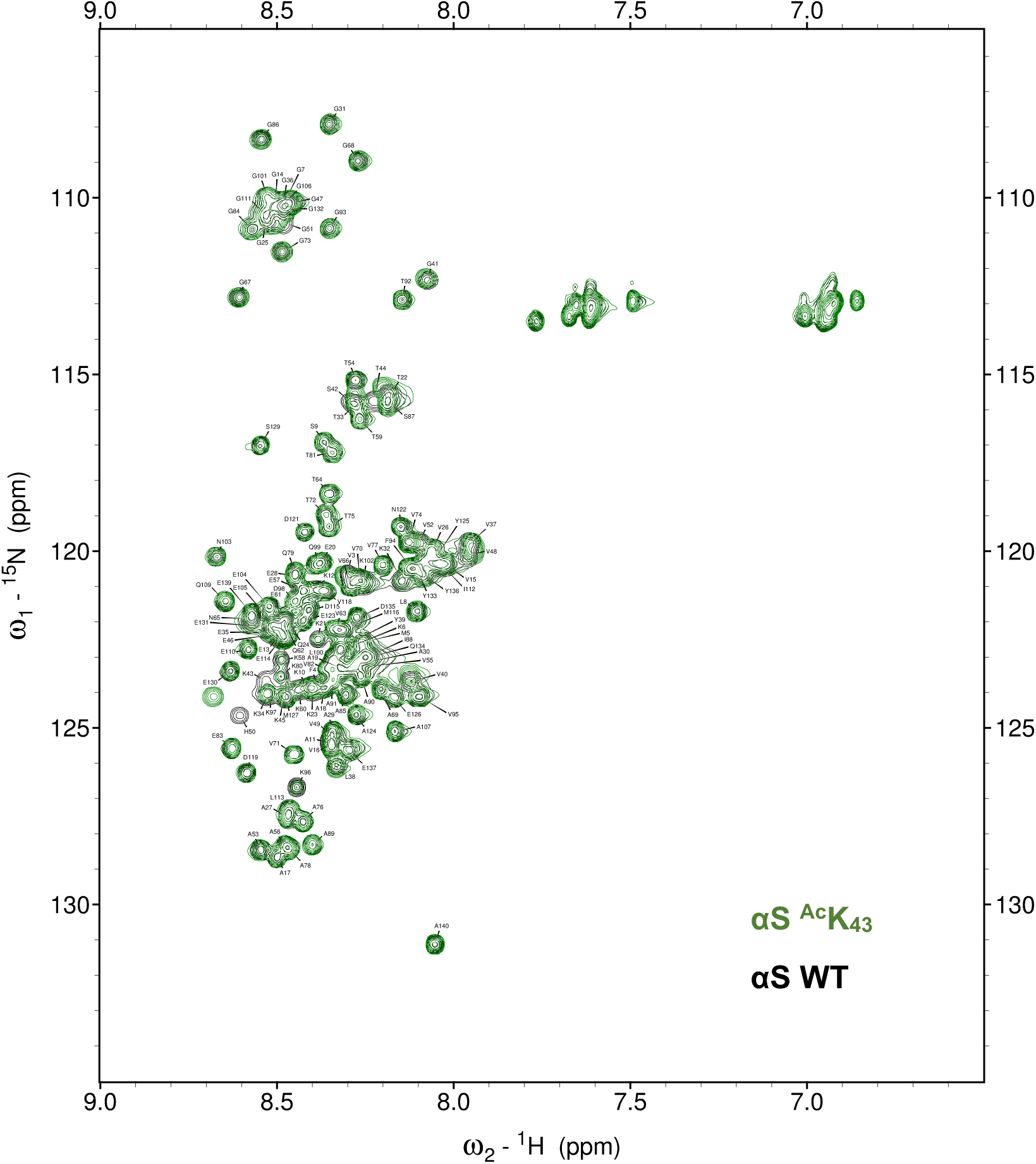
HSQC spectra acquired for αS-^Ac^K_43_ in buffer, overlayed with the spectra of αS WT.

**Figure 25c.**
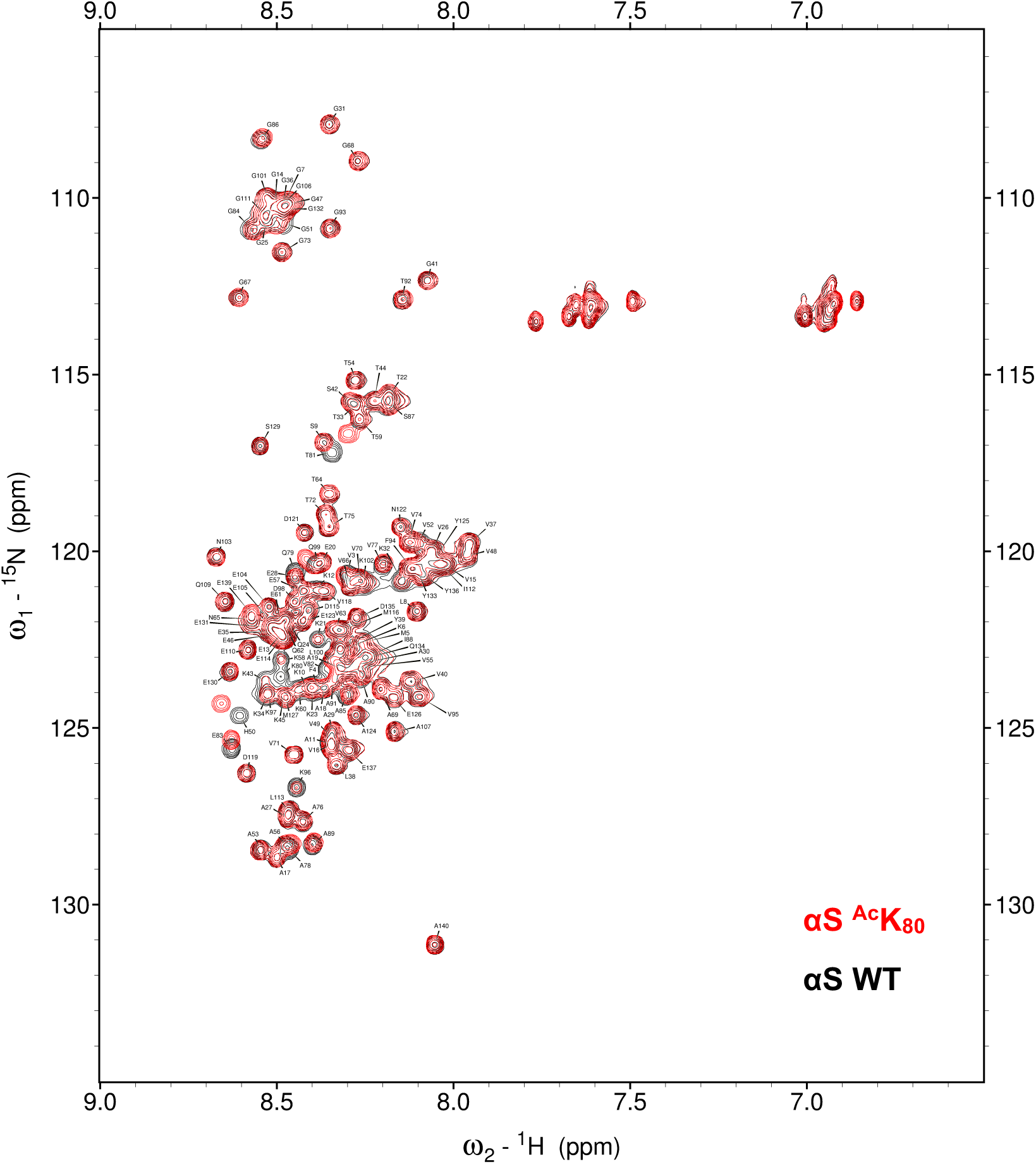
HSQC spectra acquired for αS-^Ac^K_80_ in buffer, overlayed with the spectra of αS WT.

**Figure 26.**
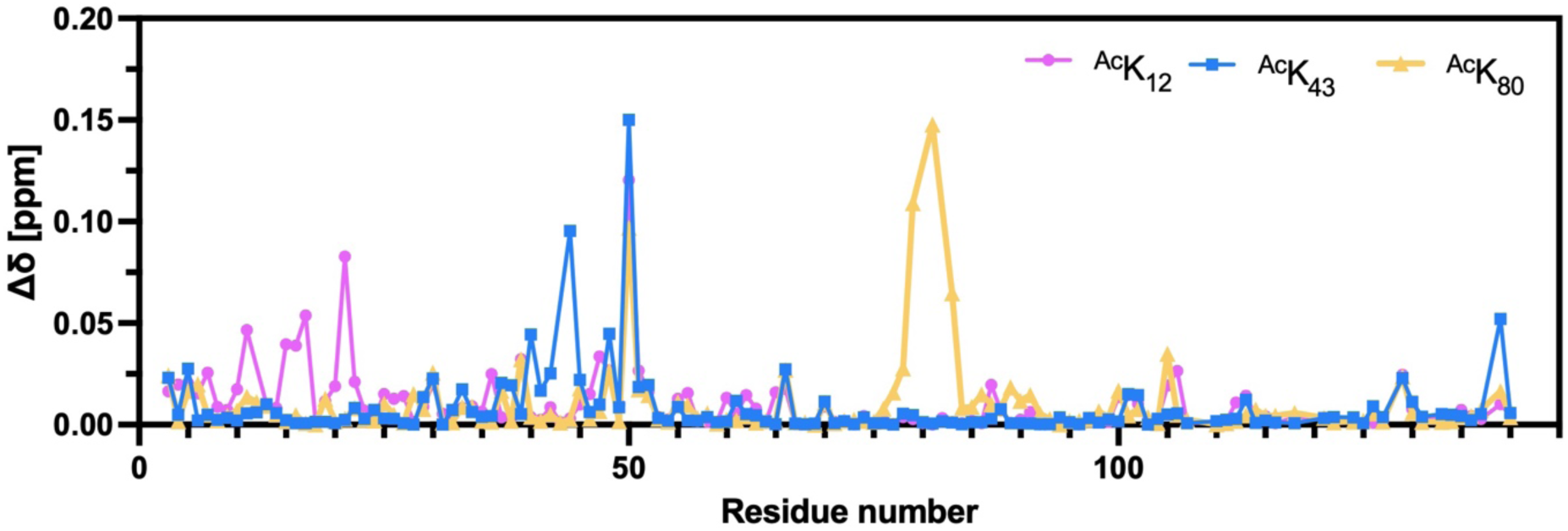
Chemical shift perturbation (Δδ; CSP) from αS-WT calculated at each residue of αS-^Ac^K_12_, αS-^Ac^K_43_ or αS-^Ac^K_80_

**Figure 27.**
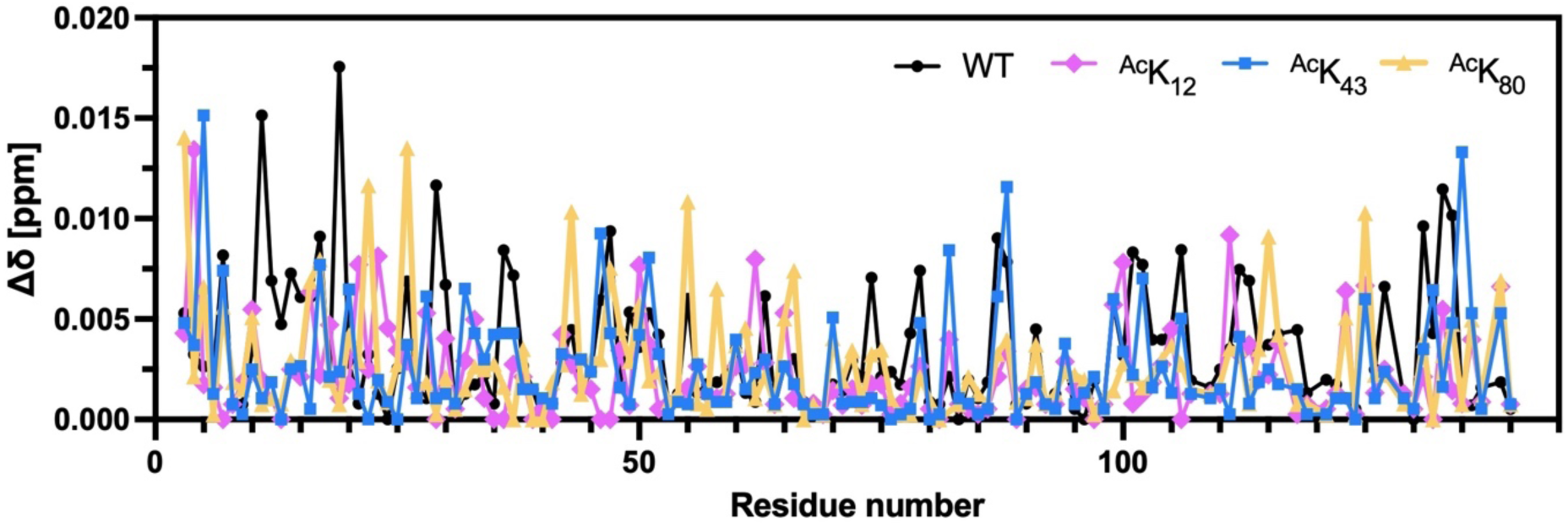
Chemical shift perturbation (Δδ; CSP) calculated at each residue of vesicle-bound αS-WT, αS-^Ac^K_43_ or αS-^Ac^K_80_. CSP was calculated by comparing spectra acquired for vesicle-bound state and spectra of free state.

**Figure 28a.**
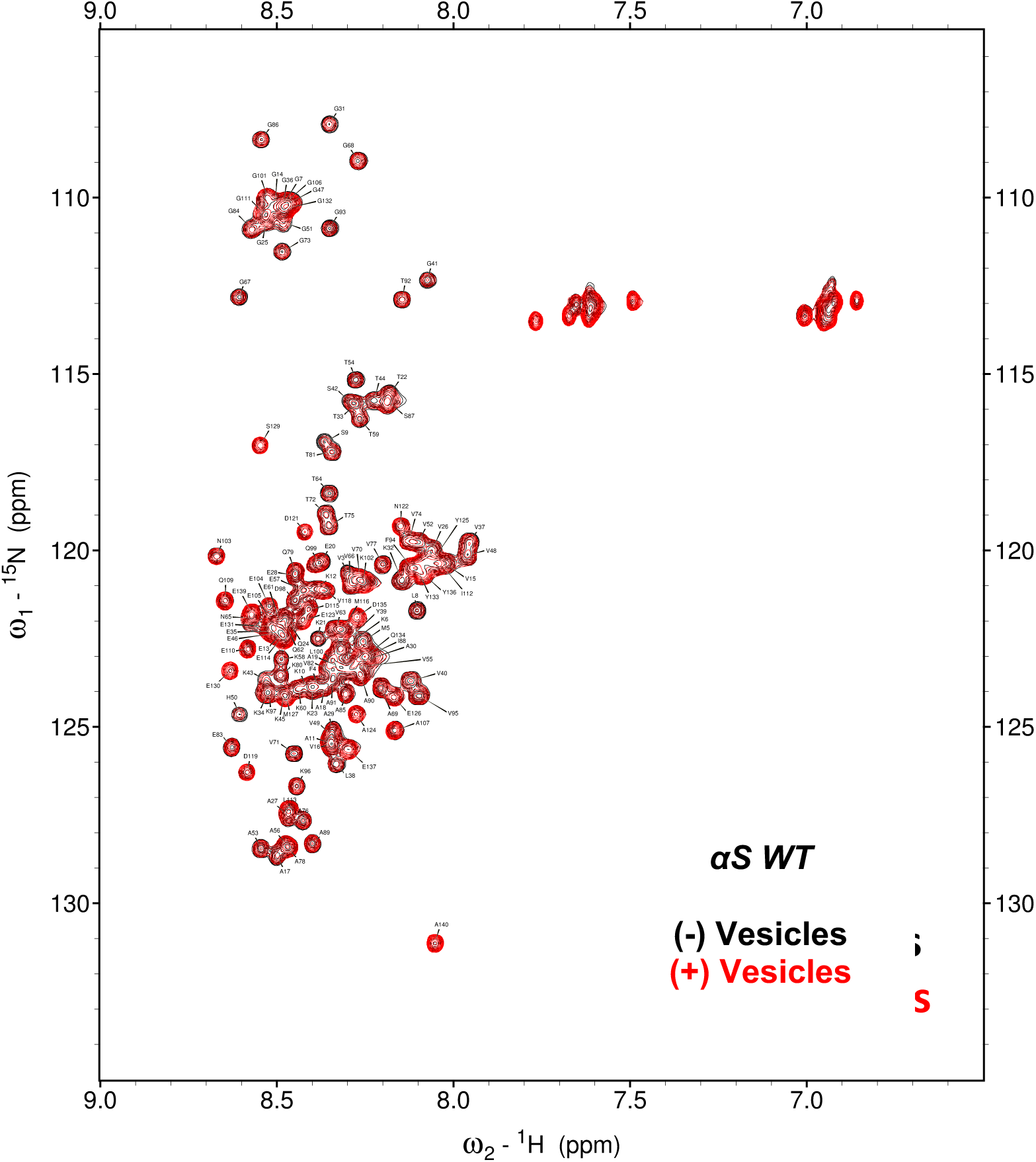
HSQC spectra acquired for free and vesicle-bound αS-WT.

**Figure 28b.**
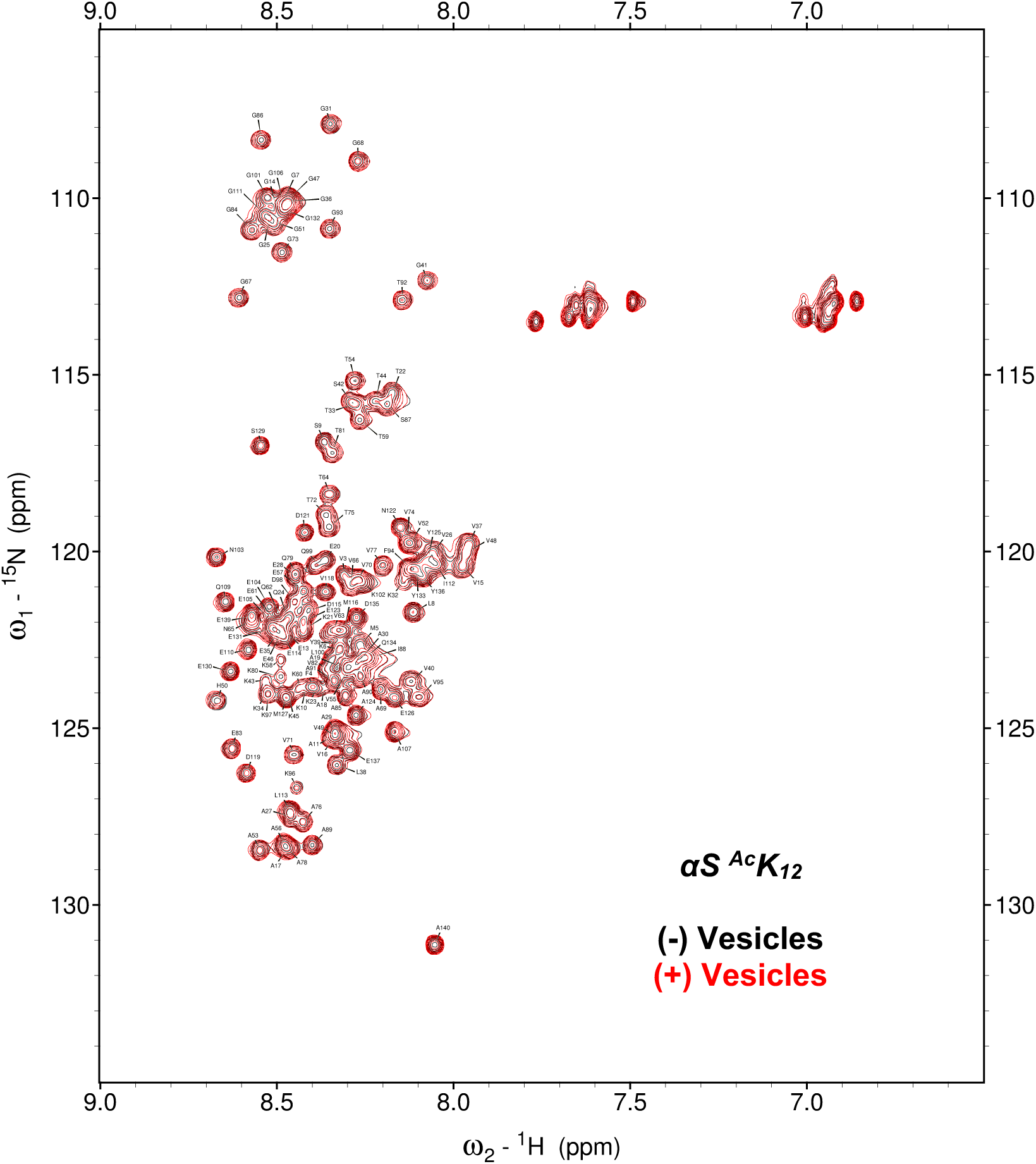
HSQC spectra acquired for free and vesicle-bound αS-^Ac^K_12_.

**Figure 28c.**
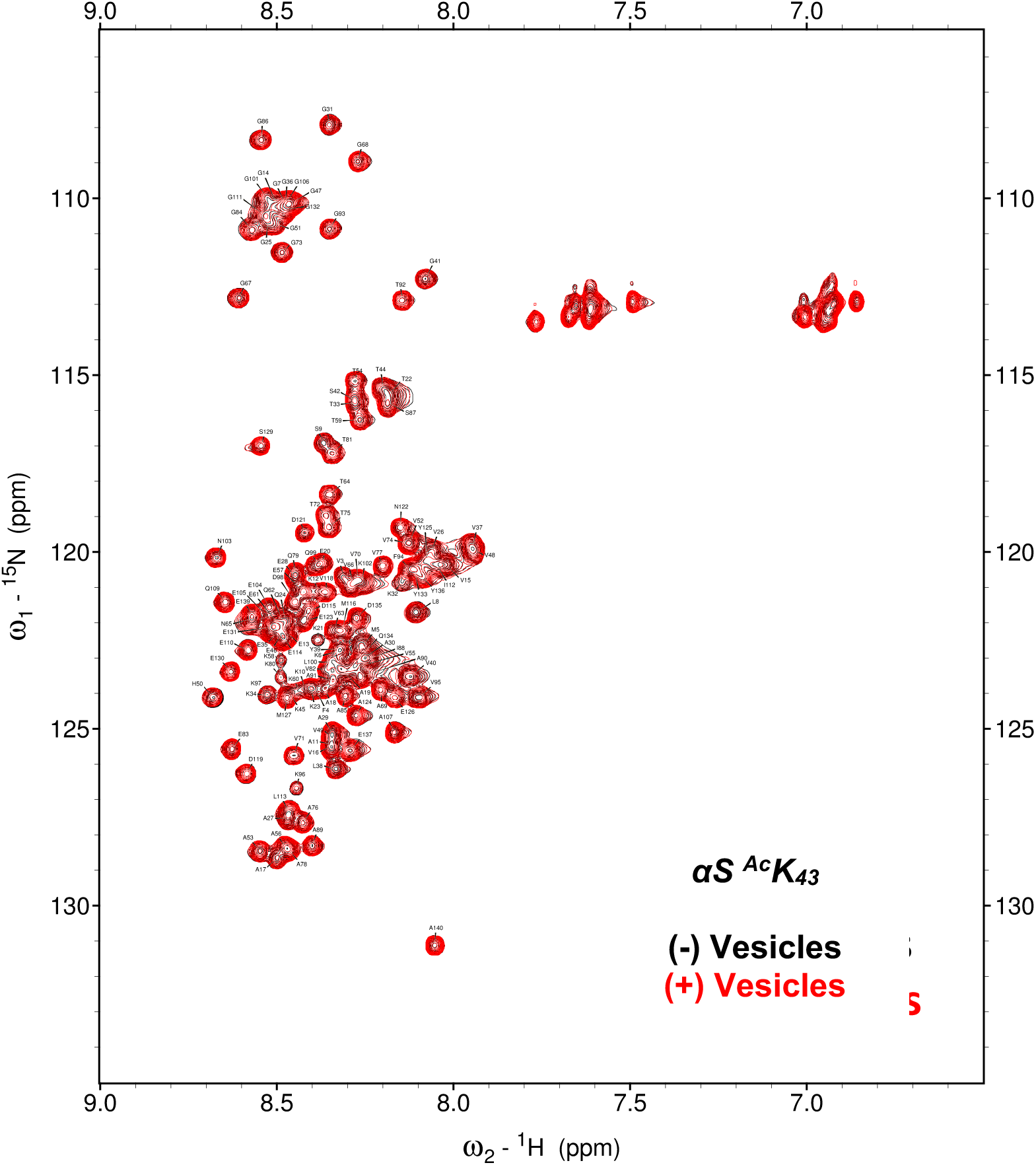
HSQC spectra acquired for free and vesicle-bound αS-^Ac^K_43_.

**Figure 28d.**
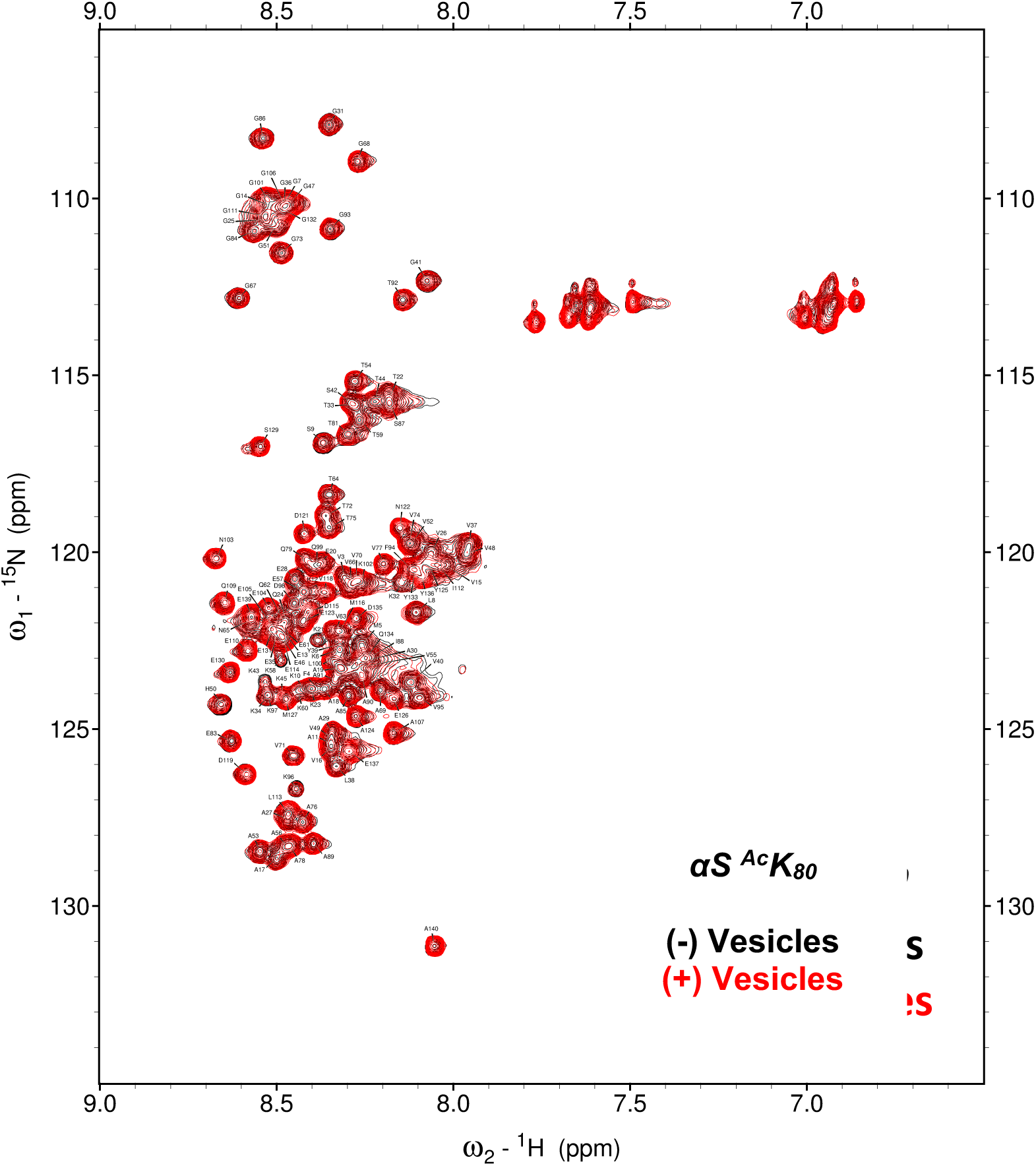
HSQC spectra acquired for free and vesicle-bound αS-^Ac^K_80_.

**Figure 29.**
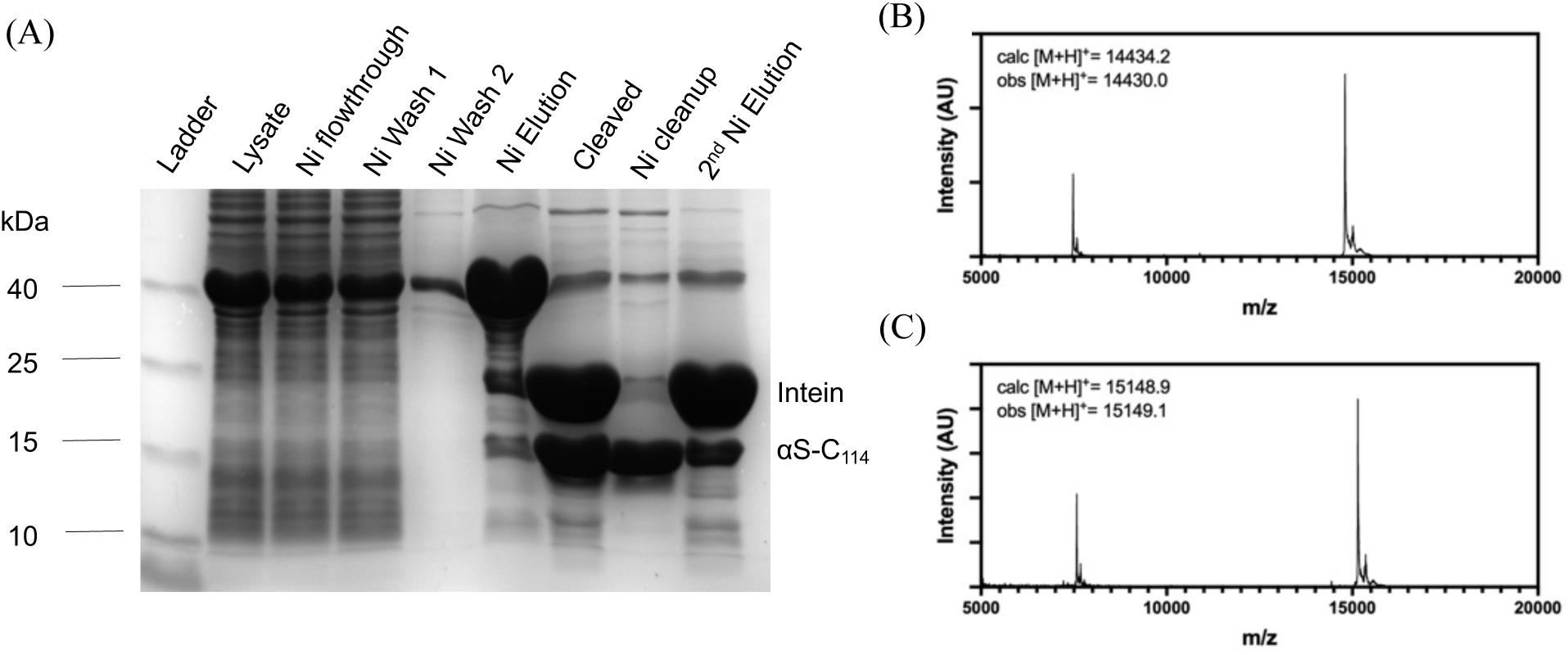
Recombinant αS-C_114_ and fluorescent labeling (a) SDS-PAGE with Coomassie staining to show affinity purification (b) MALDI-MS of purified product αS-C_114_ (c) MALDI-MS of fluorescently labeled, purified product _αS-C_Atto488_114_

**Figure 30.**
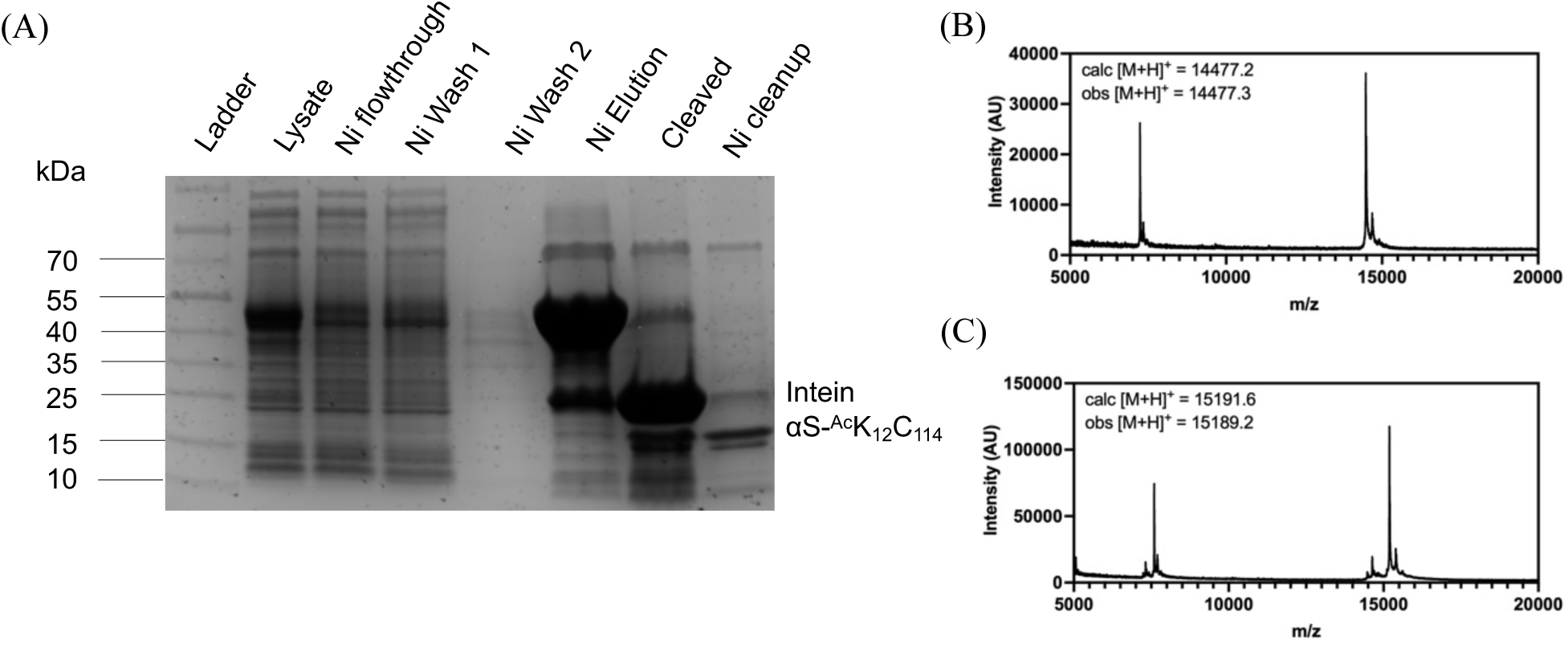
Recombinant αS- ^Ac^K_12_C_114_ and fluorescent labeling (A) SDS-PAGE with Coomassie staining to show affinity purification (B) MALDI-MS of purified product αS- ^Ac^K_12_C_114_ (C) MALDI-MS of fluorescently labeled, purified product αS- ^Ac^K_12_C^Atto488^_114_

**Figure 31.**
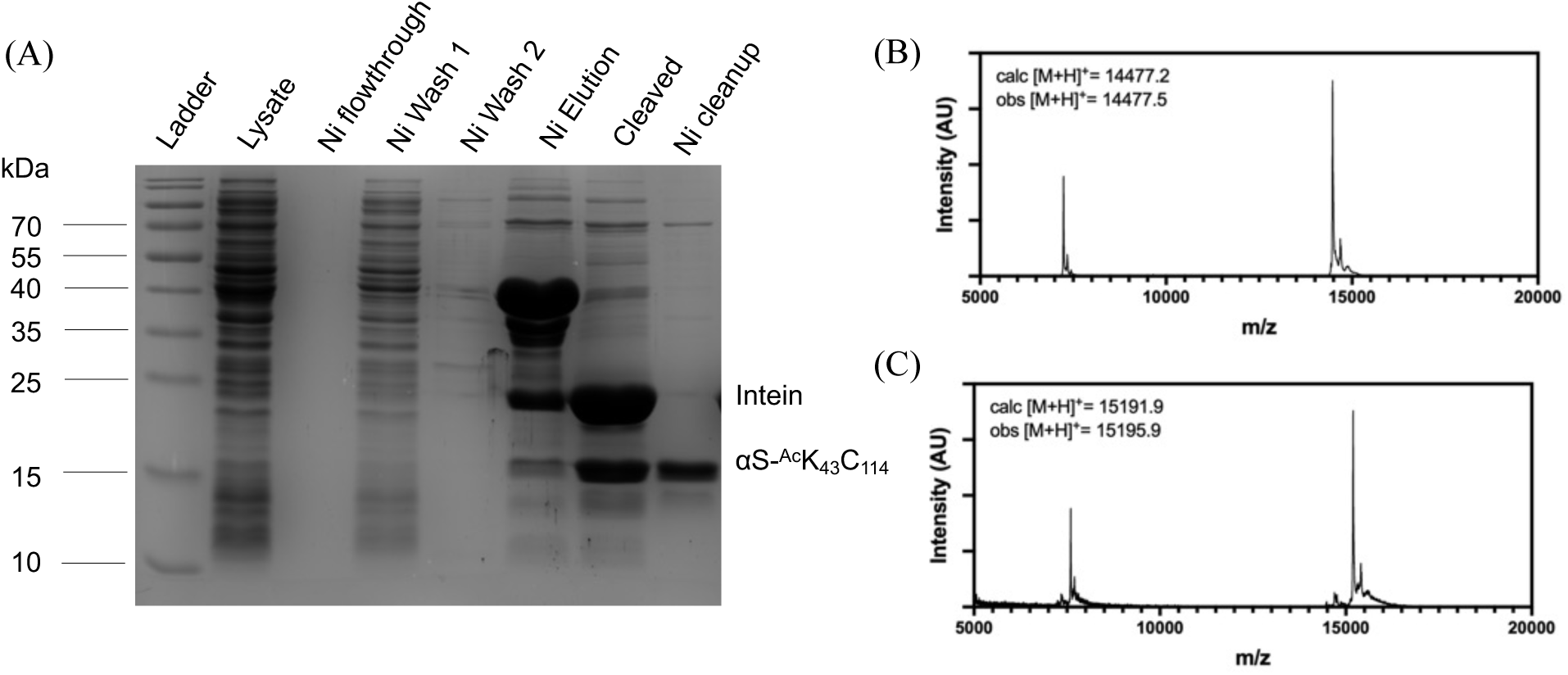
Recombinant αS- ^Ac^K_43_C_114_ and fluorescent labeling (A) SDS-PAGE with Coomassie staining to show affinity purification (B) MALDI-MS of purified product αS- ^Ac^K_43_C_114_ (C) MALDI-MS of fluorescently labeled, purified product αS- ^Ac^K_43_C^Atto488^_114_

**Figure 32.**
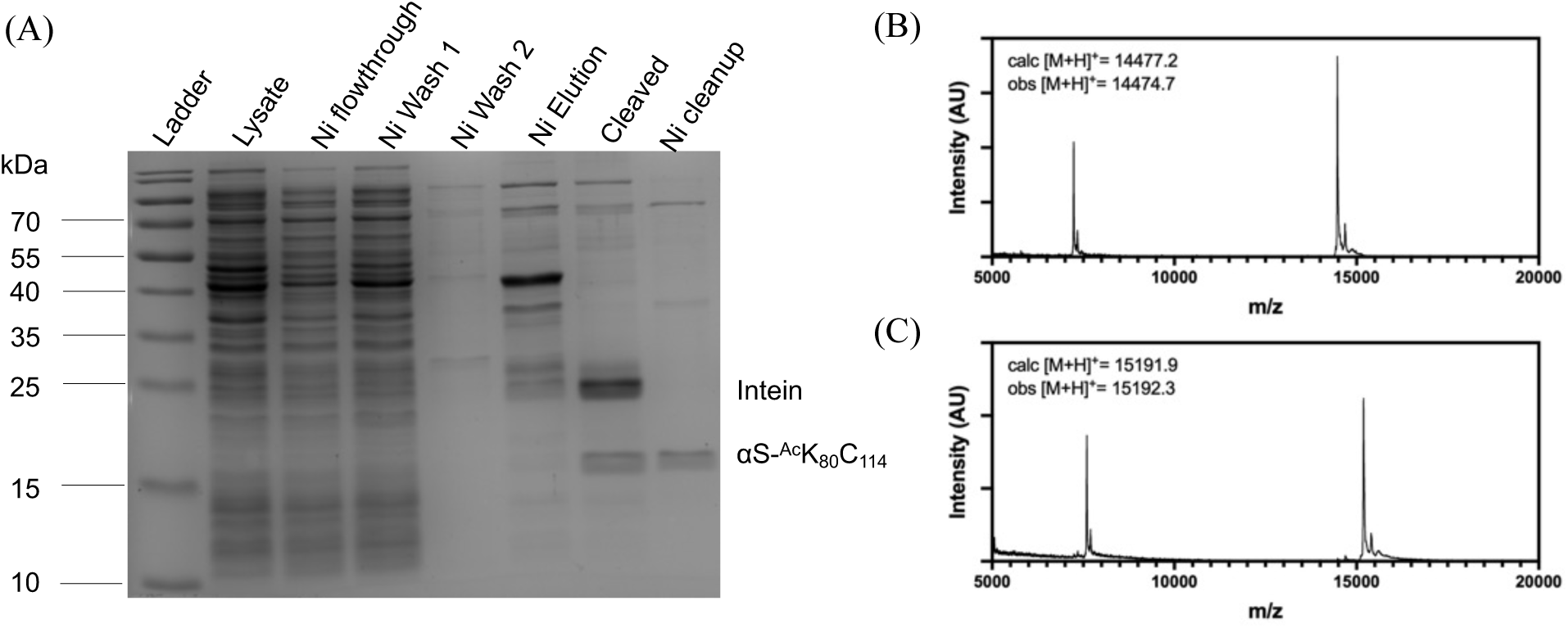
Recombinant αS- ^Ac^K_80_C_114_ and fluorescent labeling (A) SDS-PAGE with Coomassie staining to show affinity purification (B) MALDI-MS of purified product αS- ^Ac^K_80_C_114_ (C) MALDI-MS of fluorescently labeled, purified product αS- ^Ac^K_80_C^Atto488^_114_

**Figure 33.**
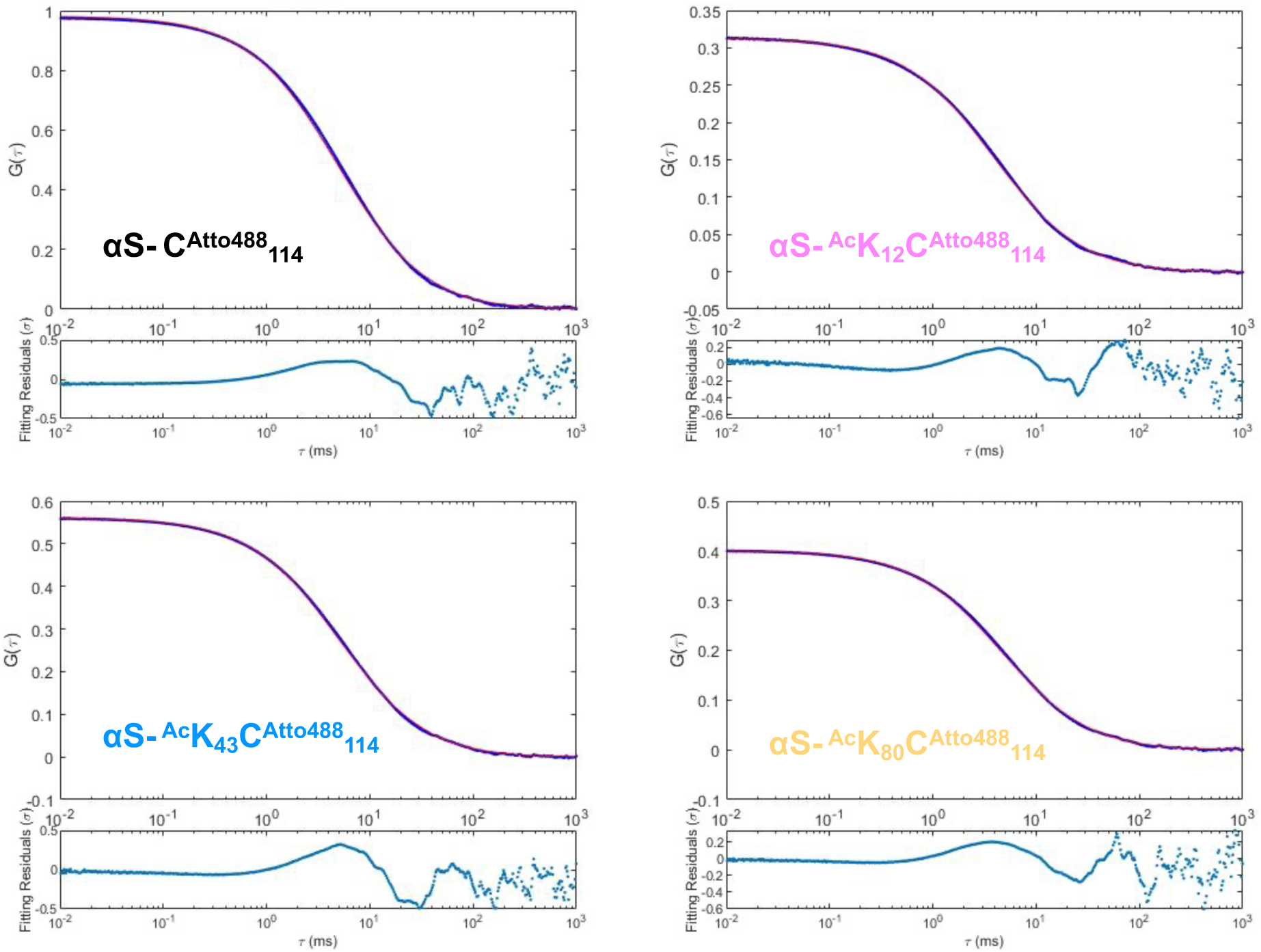
Vesicle FCS data. FCS autocorrelation curves for αS-C^Atto488^_114_ control (WT), αS-^Ac^K_12_-C^Atto488^_114,_ αS-^Ac^K_43_-C^Atto488^_114_, or αS- ^Ac^K_80_C^Atto488^_114_ with 0.1 mM, 50:50 POPS/POPC vesicles. 30 autocorrelation curves were averaged and fit to a single-component autocorrelation function to determine diffusion time.

**Figure 34.**
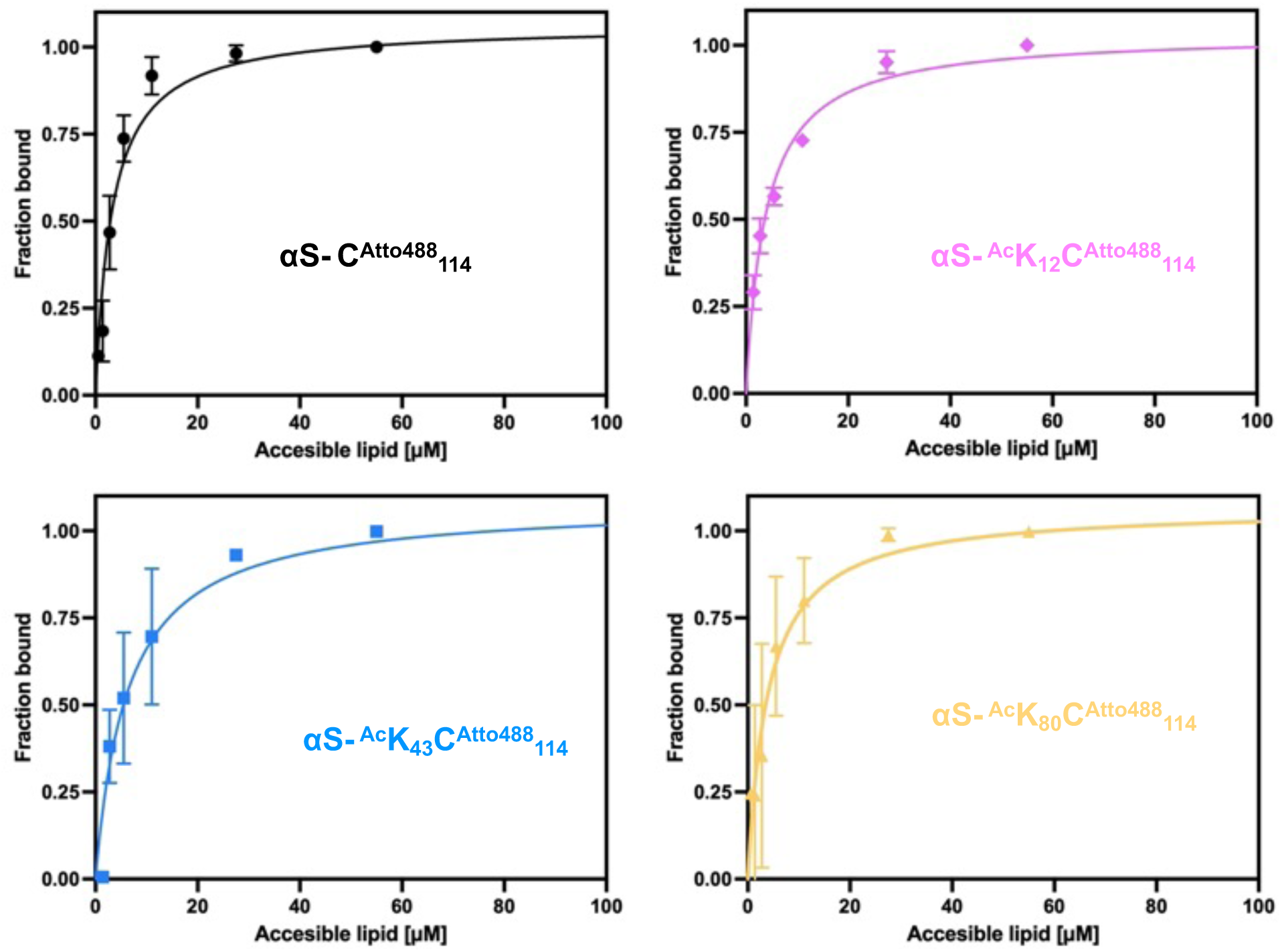
Lipid binding affinity of acetylated αS determined by FCS. Individual binding curves for αS constructs with varying concentrations of 50:50 POPS/POPC vesicles.

**Figure 35.**
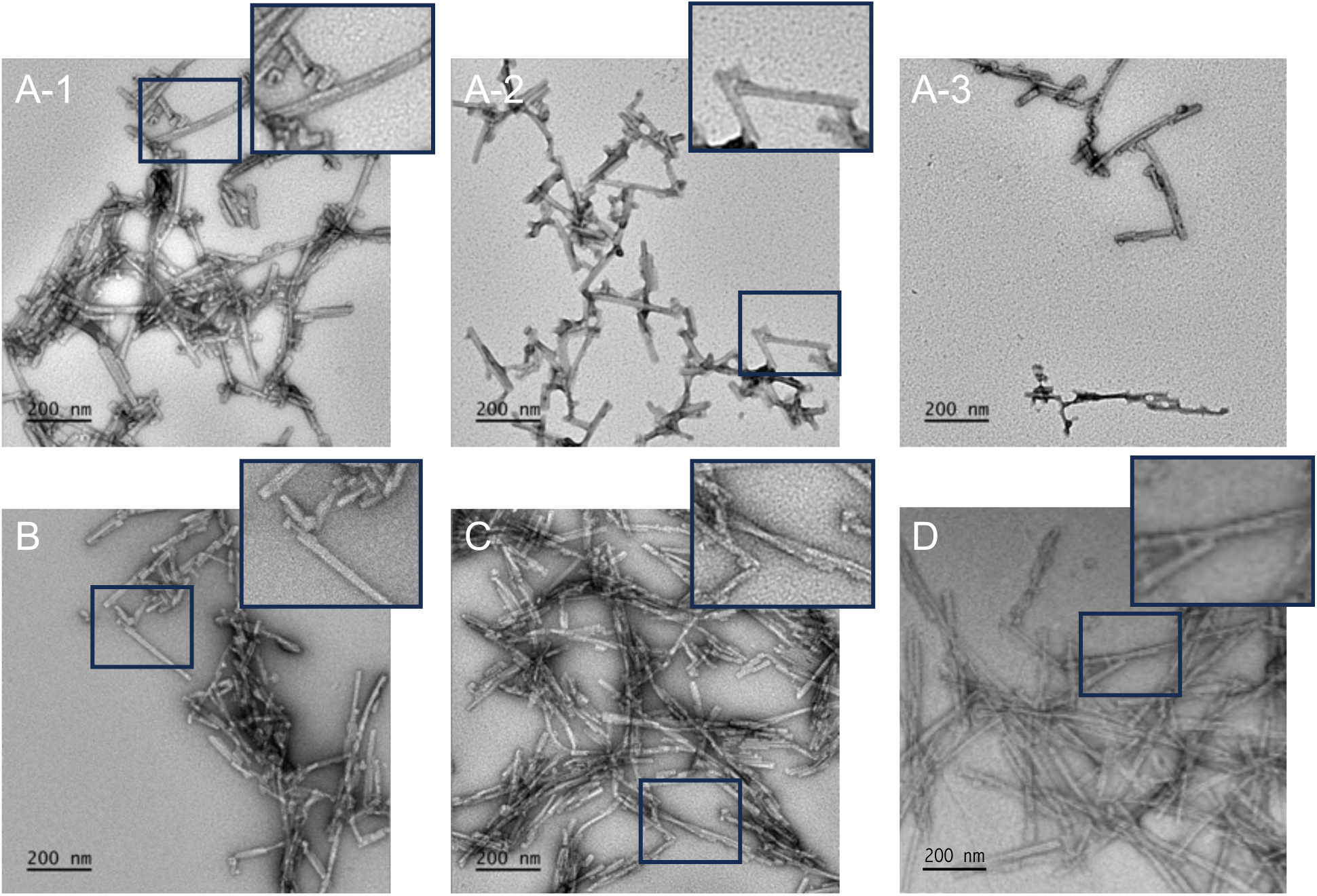
TEM images of ^Ac^K fibrils. TEM images of fibrils formed from αS monomers comprised of (A-1,A-2,A-3) 25% αS-^Ac^K_12_ (B) 25% αS-^Ac^K_43_ (C) 25% αS-^Ac^K_80_ (D) 100% αS-WT. Multiple fields of view are shown for 25% αS-^Ac^K_12_ fibrils to capture the observed heterogeneity in fibril morphology.

**Figure 36.**
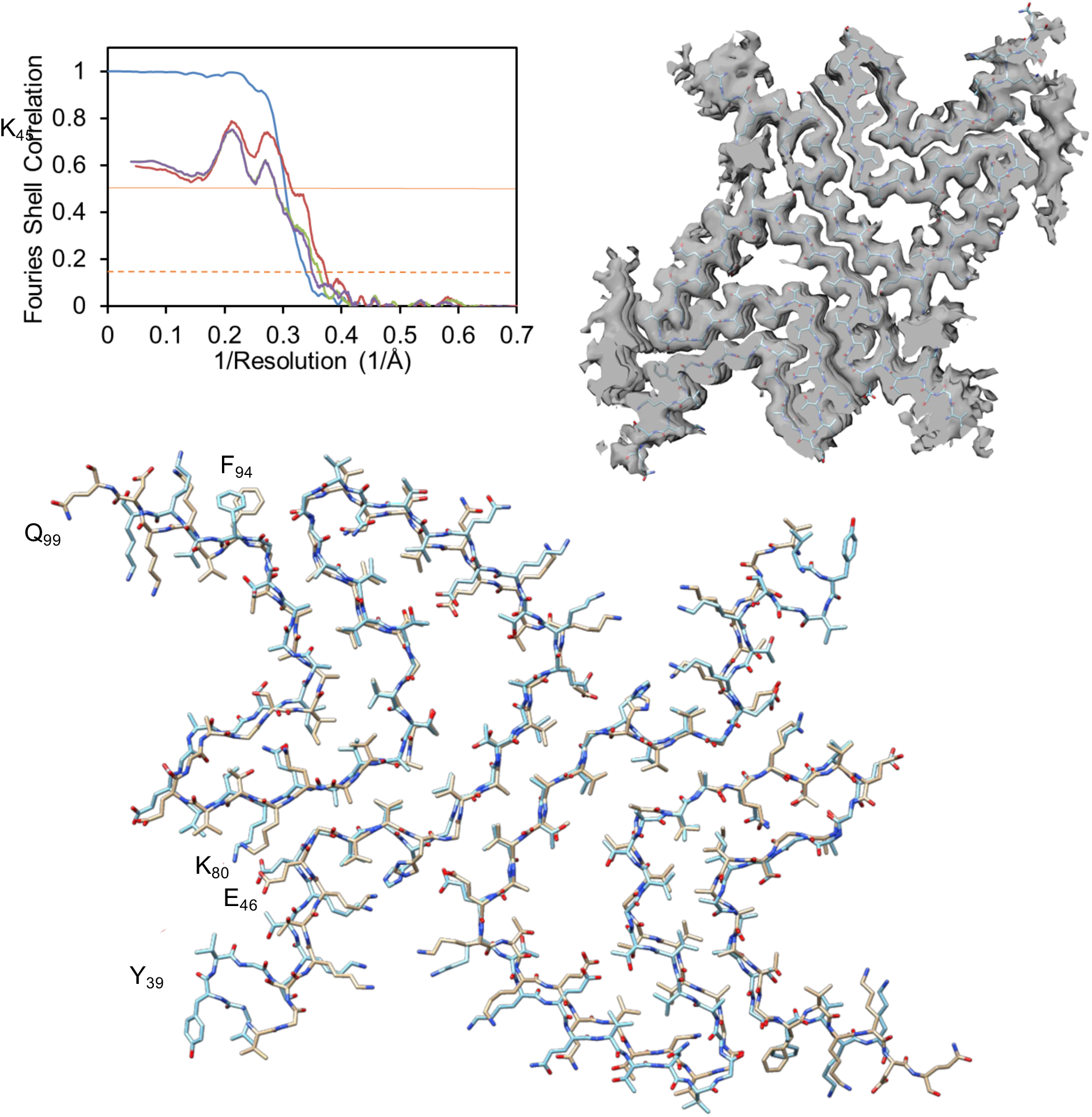
Cryo-EM structure from 25% ^Ac^K_80_ fibril preparation. A. Map and model validation. Fourier shell correlations (FSC) curves for density map and model validation. FSC curve for the density map (FSC-masked, blue), refined models versus full maps (FSC-sum, red), half maps for cross-validation (FSC-work, green and FSC-free, purple). Orange solid line and dash line correspond to FSC values of 0.5 and 0.143, respectively. B. Overlay of the cryo-EM map and the atomic model of 25% αS-^Ac^K_80_. C. Overlay of 6cu7 (cyan) and 25% αS-^Ac^K_80_ (light brown).

**Figure 37.**
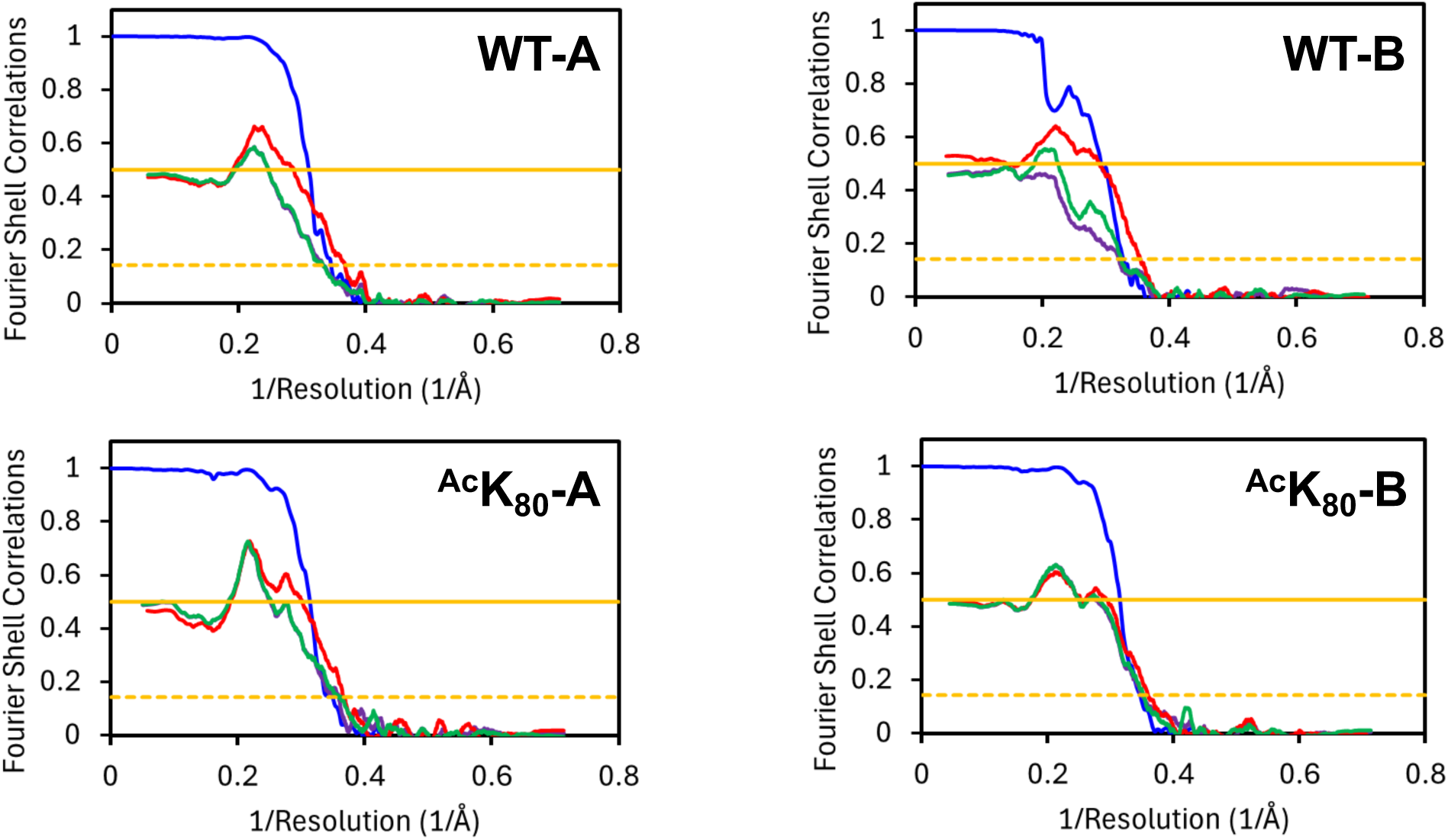
Map and model validation. Fourier shell correlations (FSC) curves for density map and model validation. FSC curve for the density map (FSC-masked, blue), refined models versus full maps (FSC-sum, red), half maps for cross-validation (FSC-work, green and FSC-free, purple). Orange solid line and dash line correspond to FSC values of 0.5 and 0.143, respectively.

**Figure 38.**
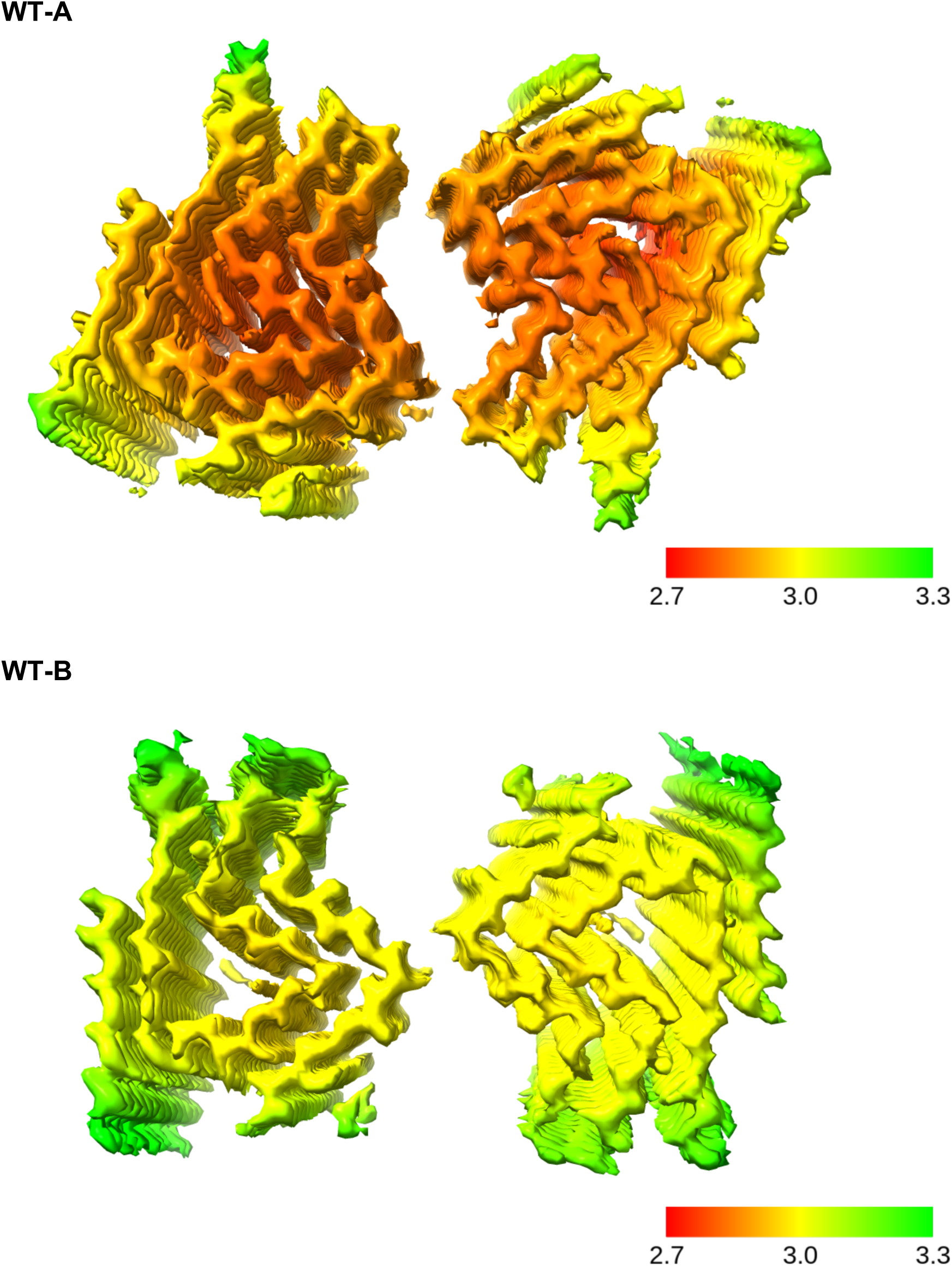
Local resolution cryo-EM maps for WT fibrils. Scale in Å.

**Figure 39.**
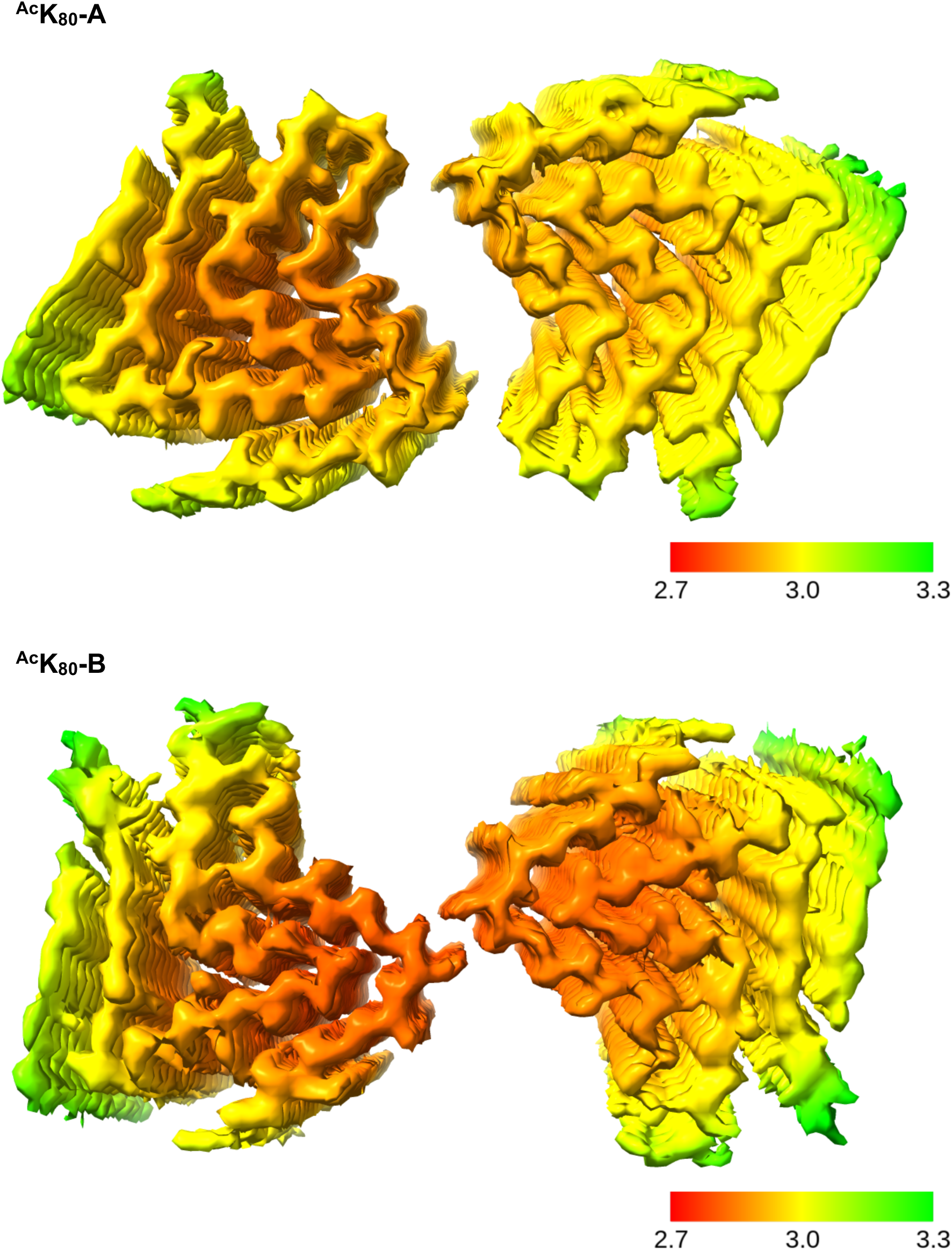
Local resolution cryo-EM maps for ^Ac^K_80_ fibrils. Scale bar in Å.

**Figure 40.**
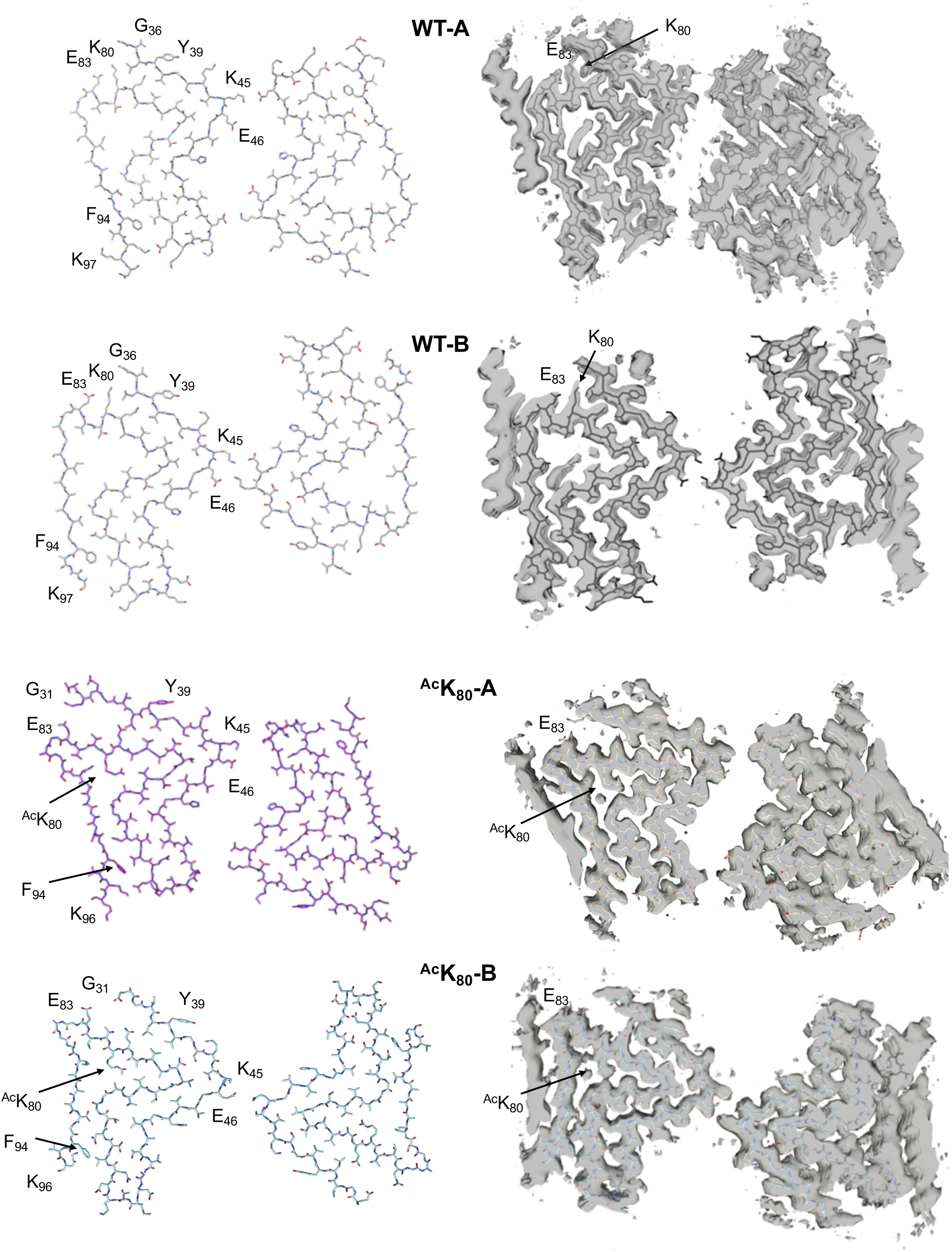
Cryo-EM structures of WT and ^Ac^K_80_ fibrils. Left: Atomic models. Right: Overlays of the cryo-EM maps and the atomic models.

**Figure 41.**
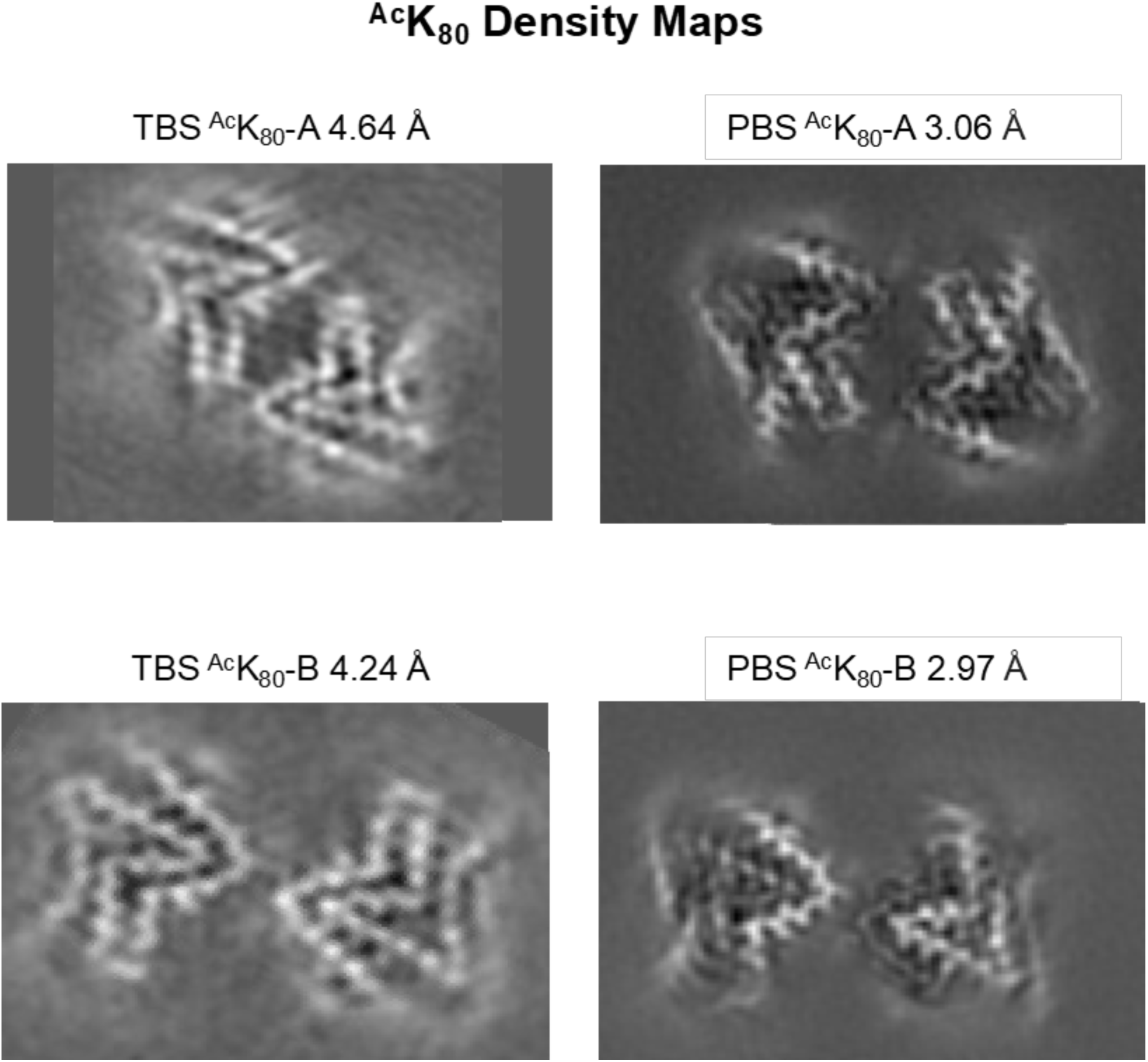
Comparison of cryo-EM density for ^Ac^K_80_ fibrils formed in TBS and PBS.

**Figure 42.**
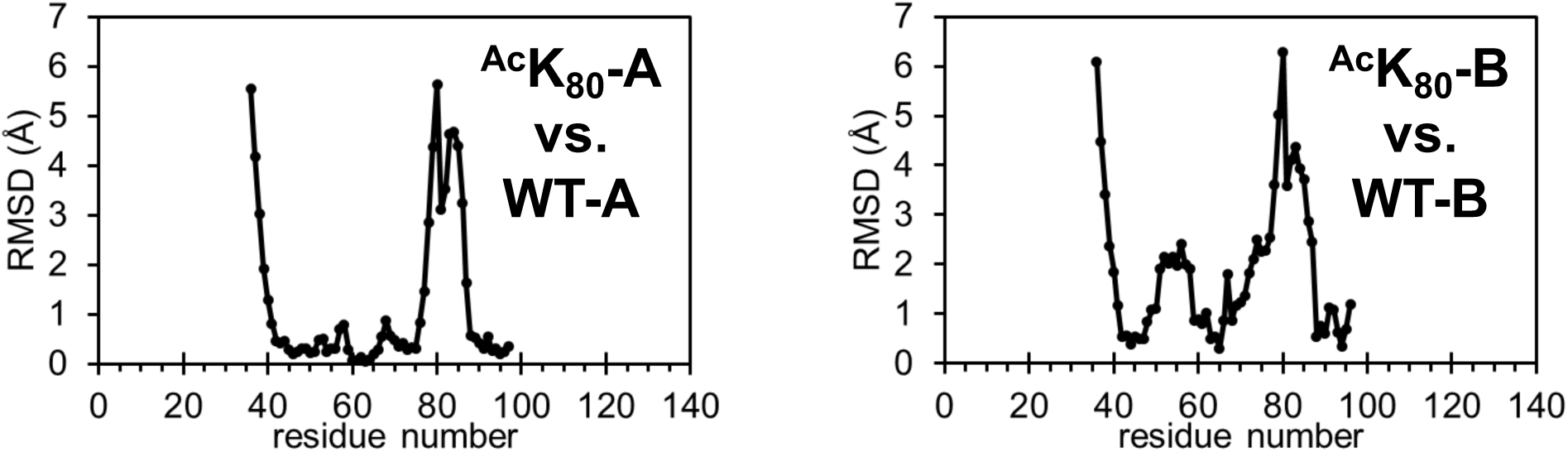
Root mean square deviation (RMSD) of Cα atoms for (left) WT-A and ^Ac^K_80_-A and (right) WT-B and ^Ac^K_80_-B, calculated using ChimeraX.

**Figure 43.**
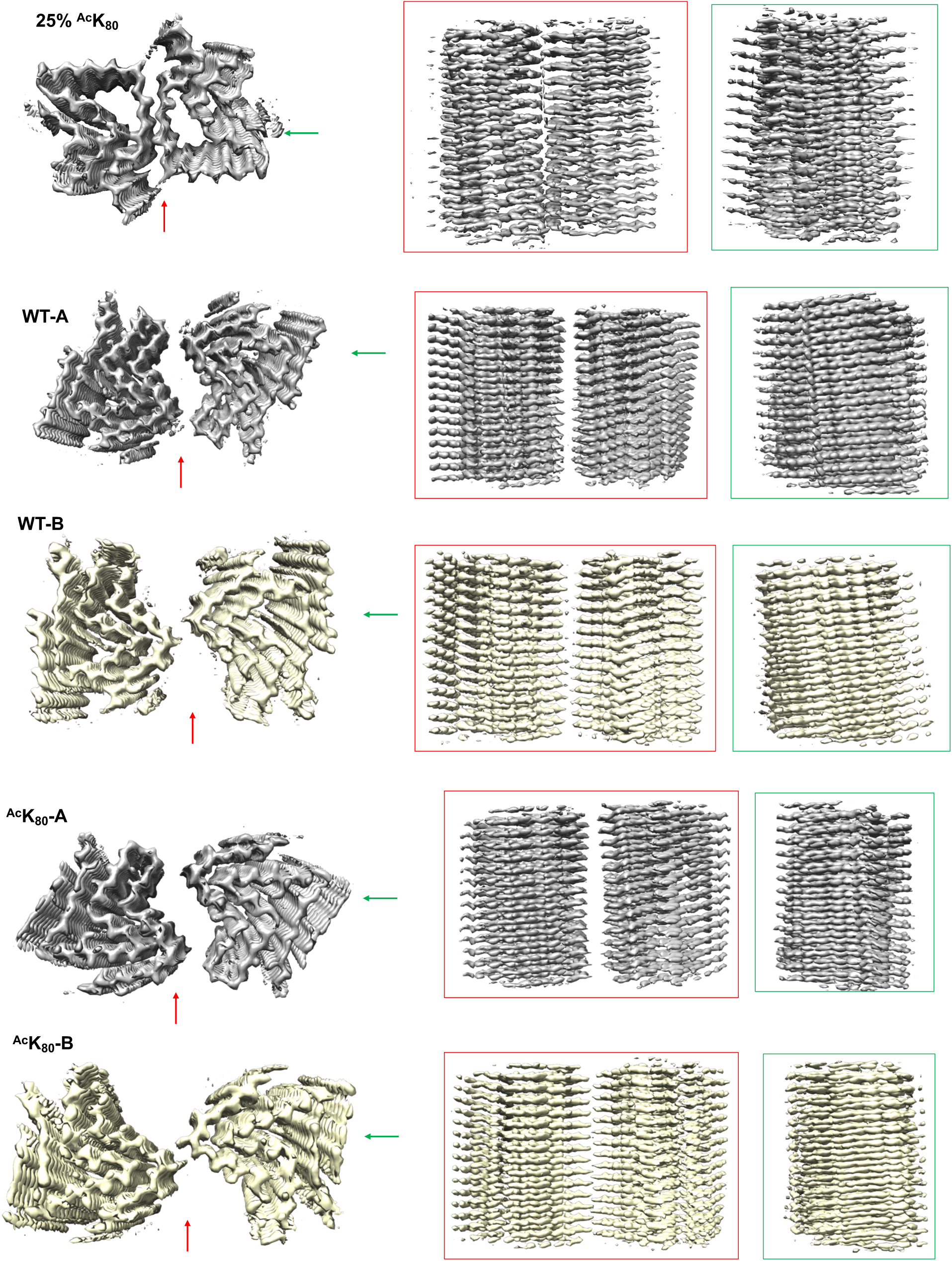
Views of cryo-EM density perpendicular to fibril axis for all structures.

**Figure 44.**
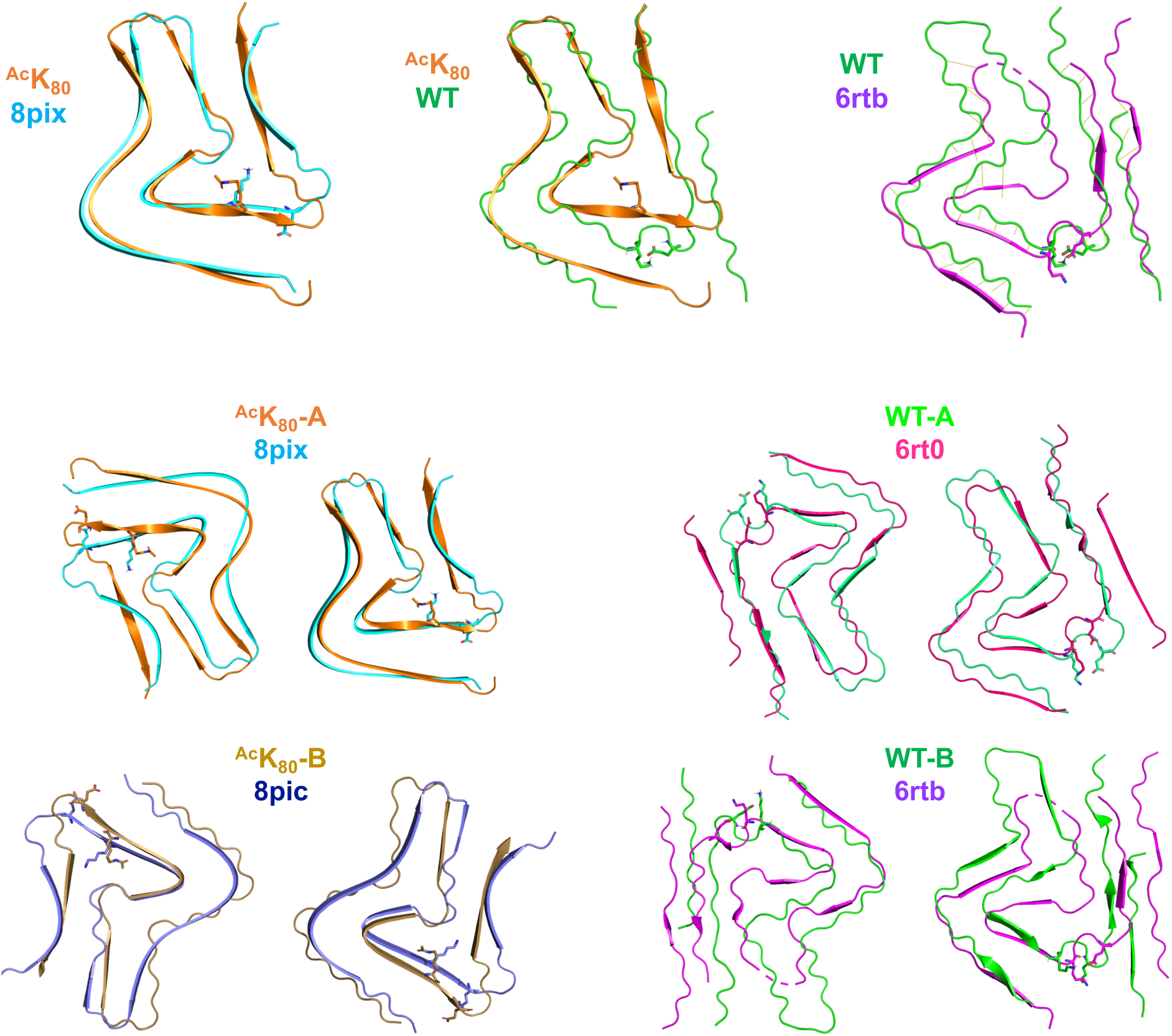
Comparison of ^Ac^K_80_ fibril structure to other cryo-EM structures. Overlays performed using alignment tool in PyMOL show similarity to published fibril polymorphs. 8pix/8pic: Frey et. al. (2024) On the pH-dependence of α-synuclein amyloid polymorphism and the role of secondary nucleation in seed-based amyloid propagation. *eLife* 12, RP93562. 10.7554/eLife.93562. 6rto/6rtb: Guerrero-Ferreira et al. (2019). Two new polymorphic structures of human full-length alpha-synuclein fibrils solved by cryo-electron microscopy. *eLife* 8, e48907. 10.7554/eLife.48907.

**Figure 45.**
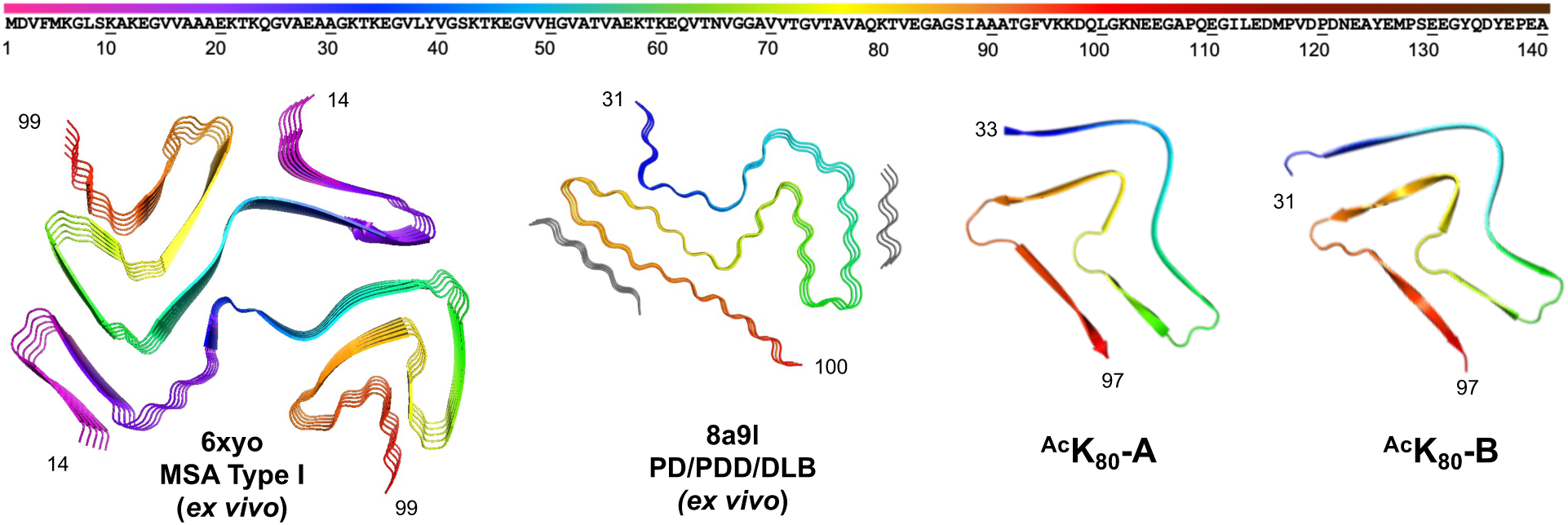
Comparison of ^Ac^K_80_ fibril structures to *ex vivo* cryo-EM structures. 6xyo: Schweighauser et al. (2020). Structures of α-synuclein filaments from multiple system atrophy. *Nature*. 10.1038/s41586-020-2317-6. Yang, et al. (2022). Structures of α-synuclein filaments from human brains with Lewy pathology. Nature 610, 791-795. 10.1038/s41586-022-05319-3.

**Figure 46.**
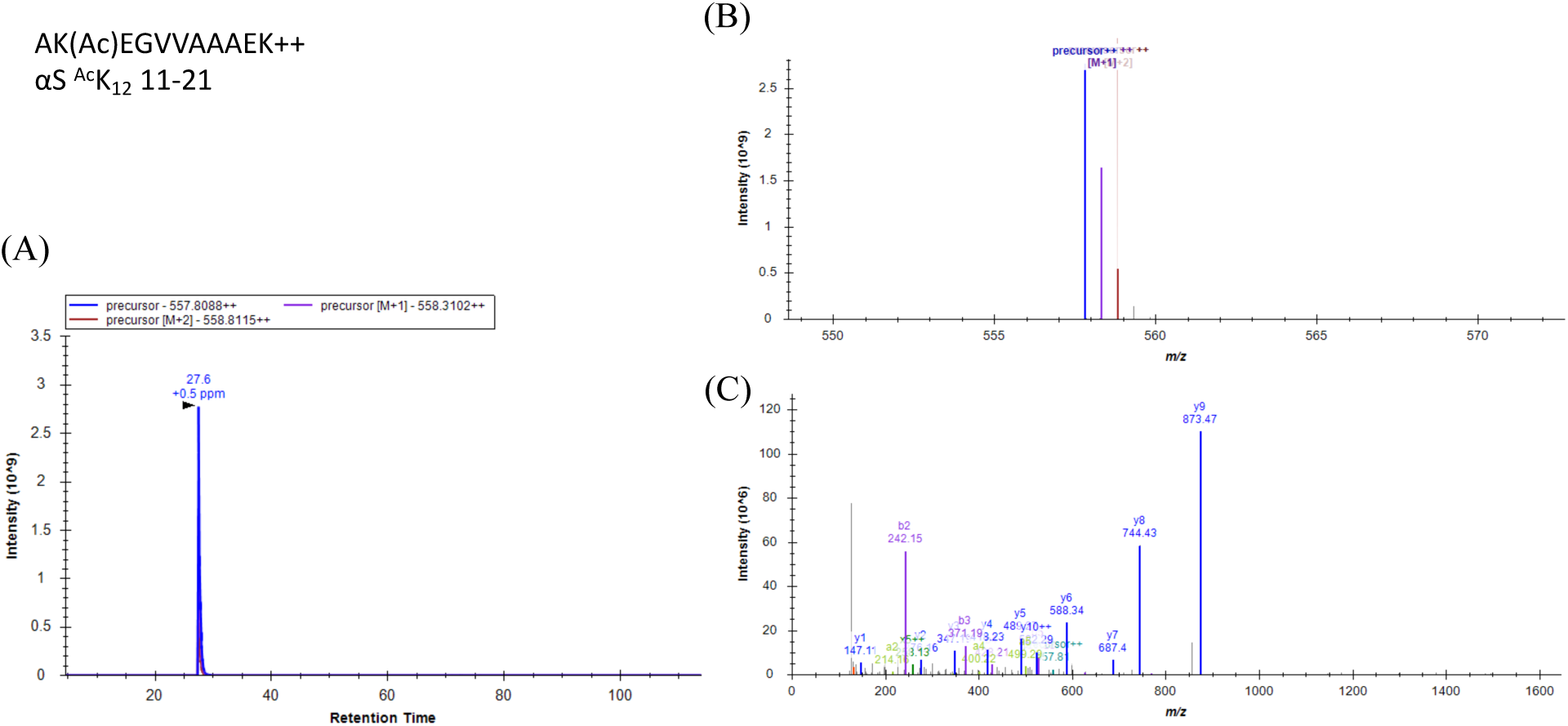
MS spectra of ^Ac^K_12_ peptide from αS ^Ac^K_12_ standard. (A) Extracted ion chromatogram (EIC) (B) MS1 spectrum (C) MS2 spectrum with fragment annotation

**Figure 47.**
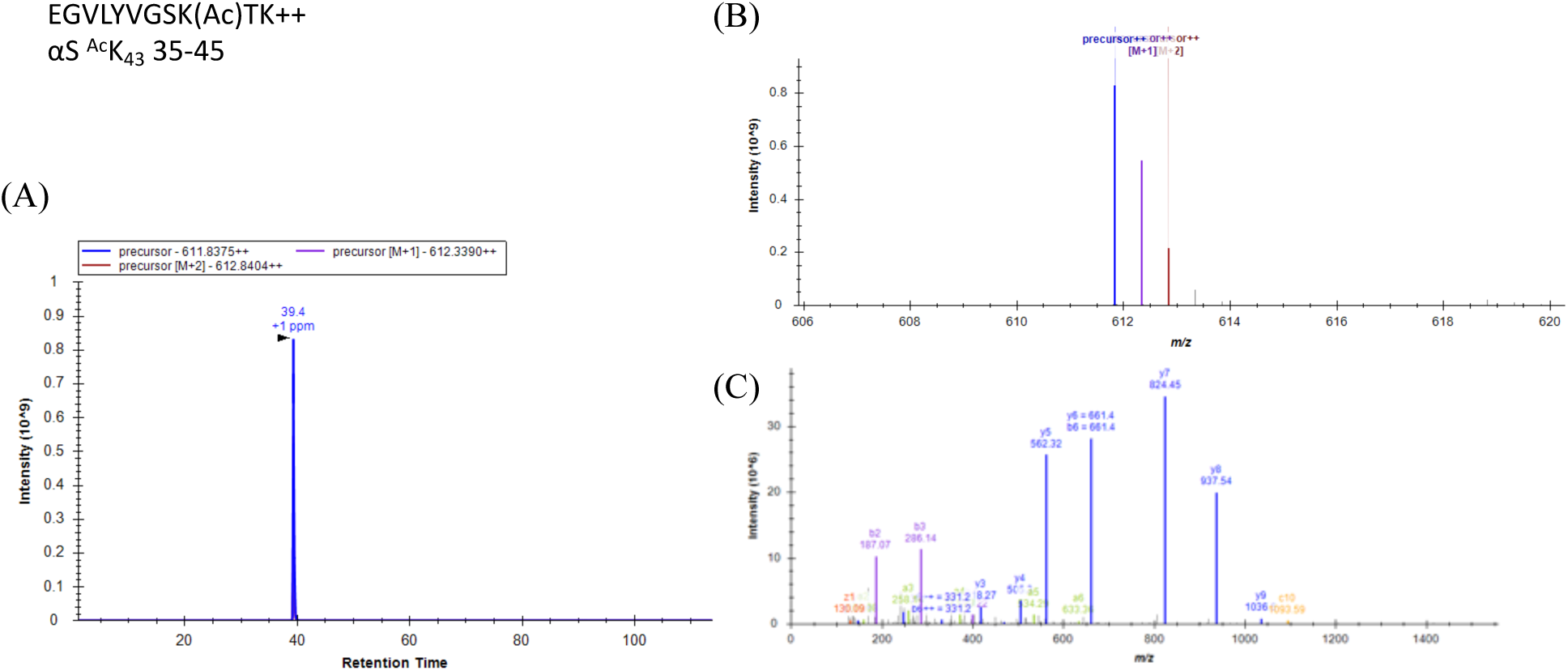
MS spectra of ^Ac^K_43_ peptide from αS ^Ac^K_43_ standard. (A) Extracted ion chromatogram (EIC) (B) MS1 spectrum (C) MS2 spectrum with fragment annotation.

**Figure 48.**
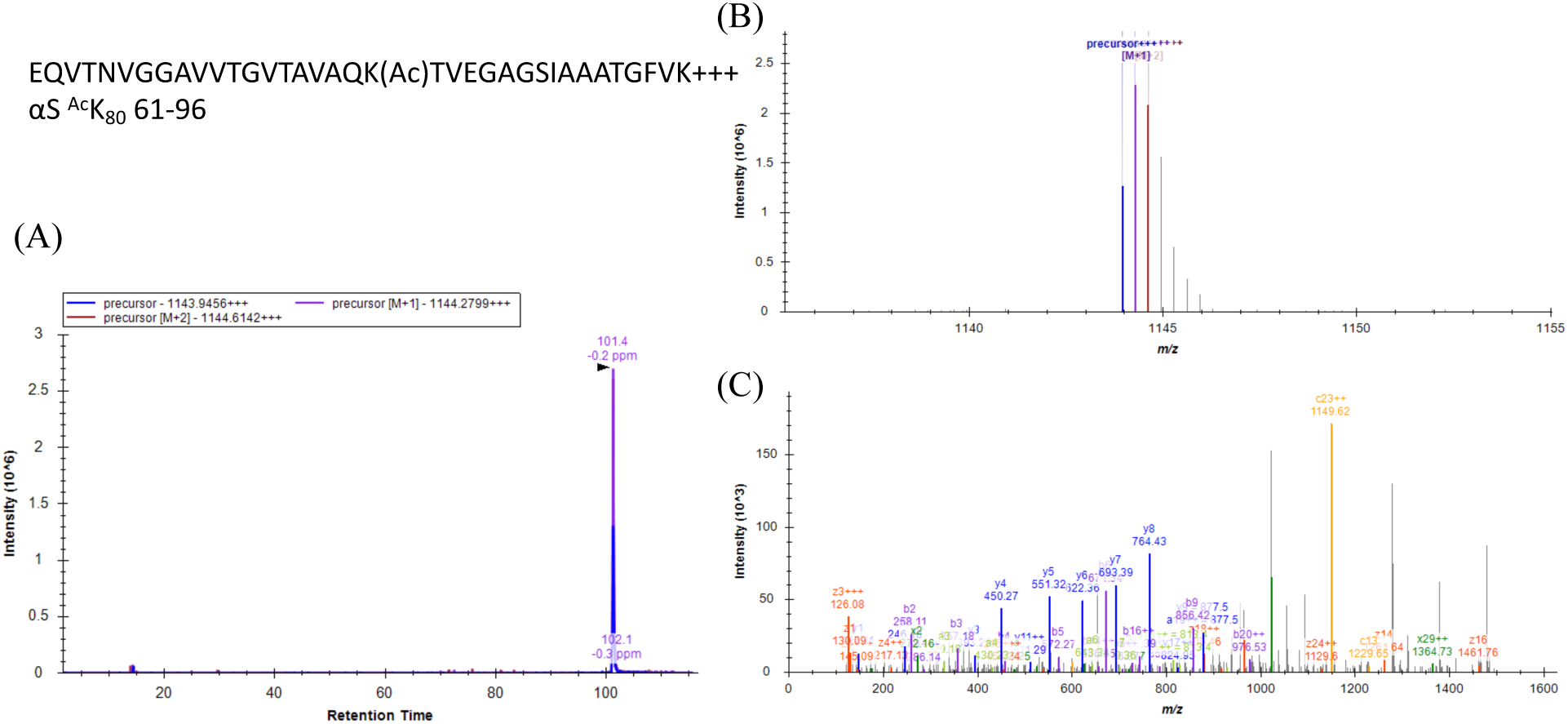
MS spectra of ^Ac^K_80_ peptide from αS ^Ac^K_80_ standard. (A) Extracted ion chromatogram (EIC) (B) MS1 spectrum (C) MS2 spectrum with fragment annotation.

**Figure 49.**
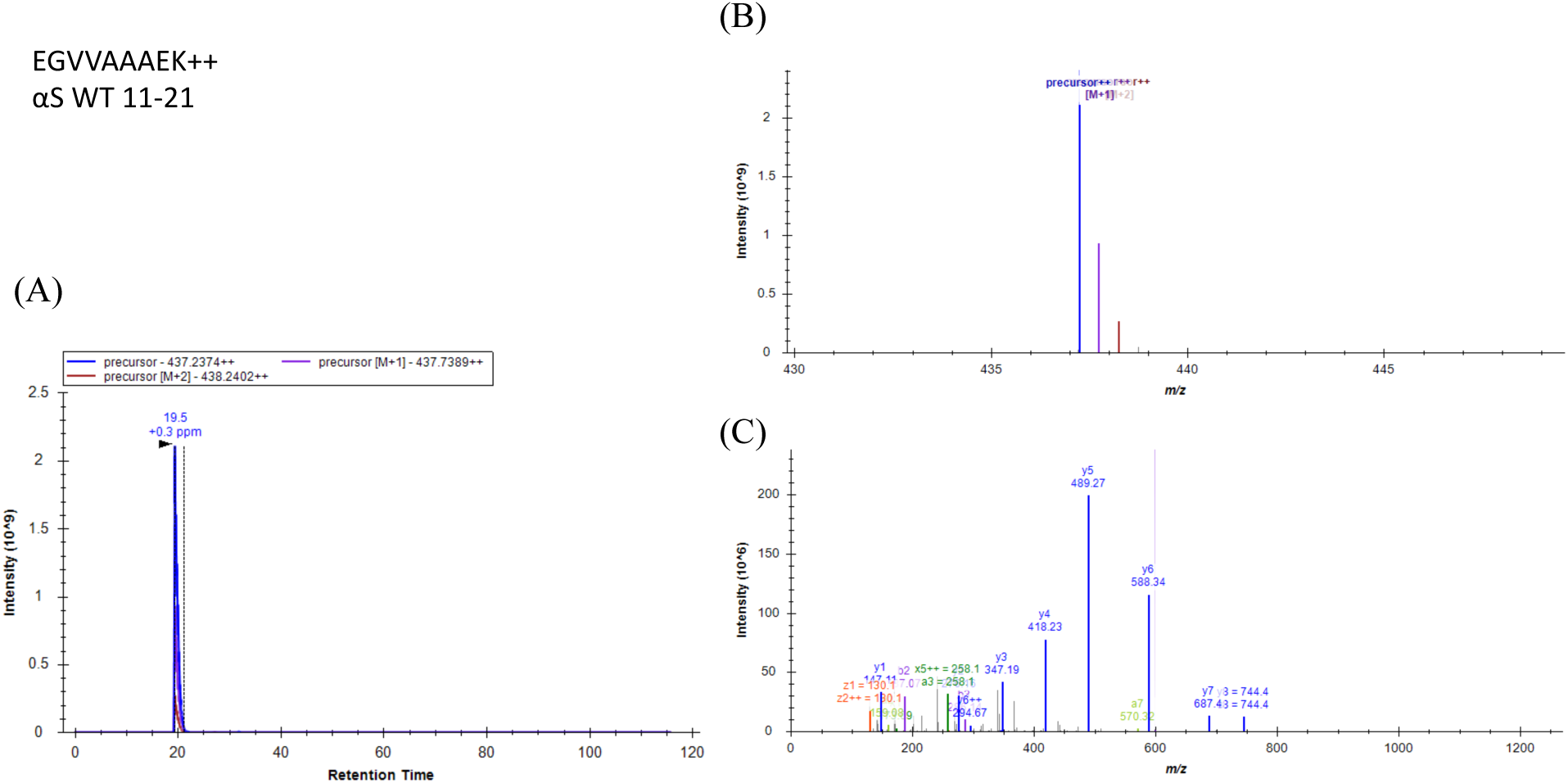
MS spectra of unmodified K_12_ peptide from αS WT standard. (A) Extracted ion chromatogram (EIC) (B) MS1 spectrum (C) MS2 spectrum with fragment annotation.

**Figure 50.**
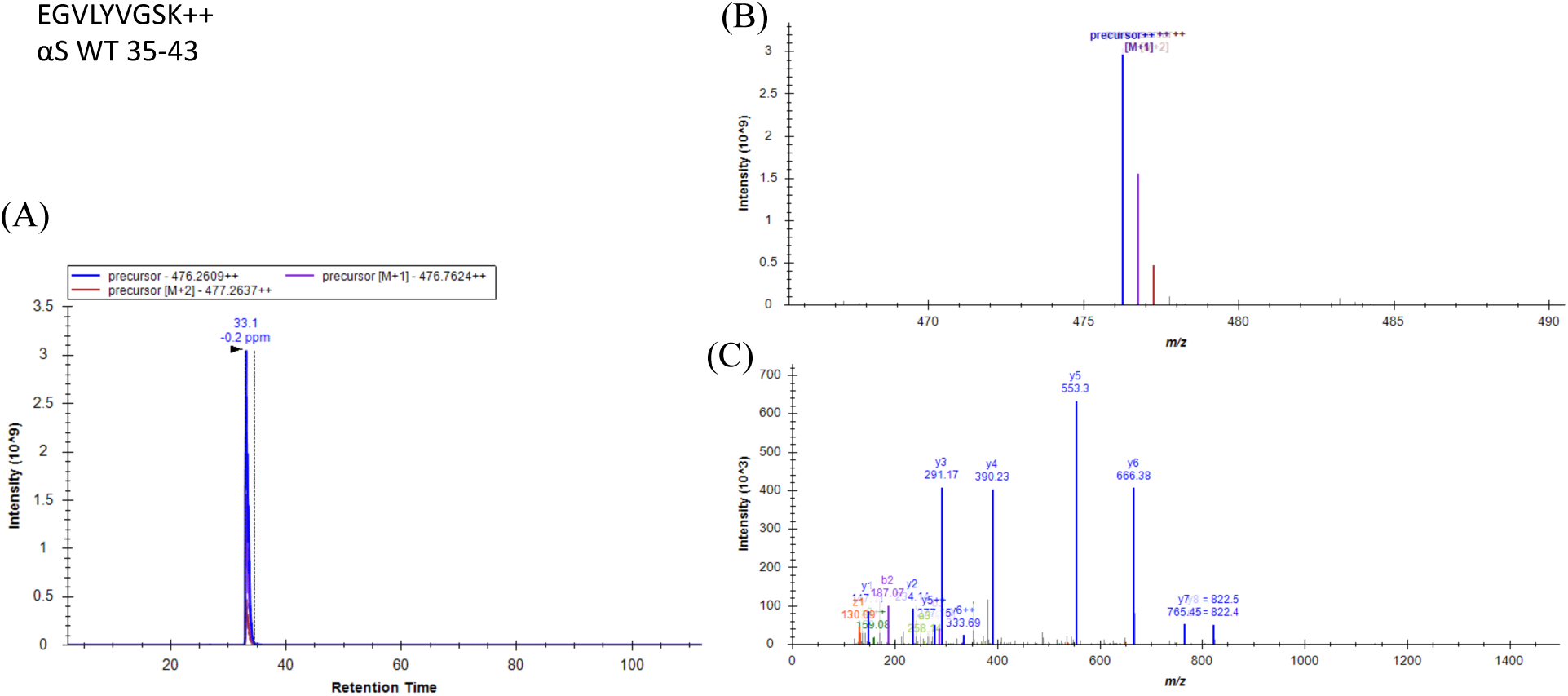
MS spectra of unmodified K_43_ peptide from αS WT standard. (A) Extracted ion chromatogram (EIC) (B) MS1 spectrum (C) MS2 spectrum with fragment annotation.

**Figure 51.**
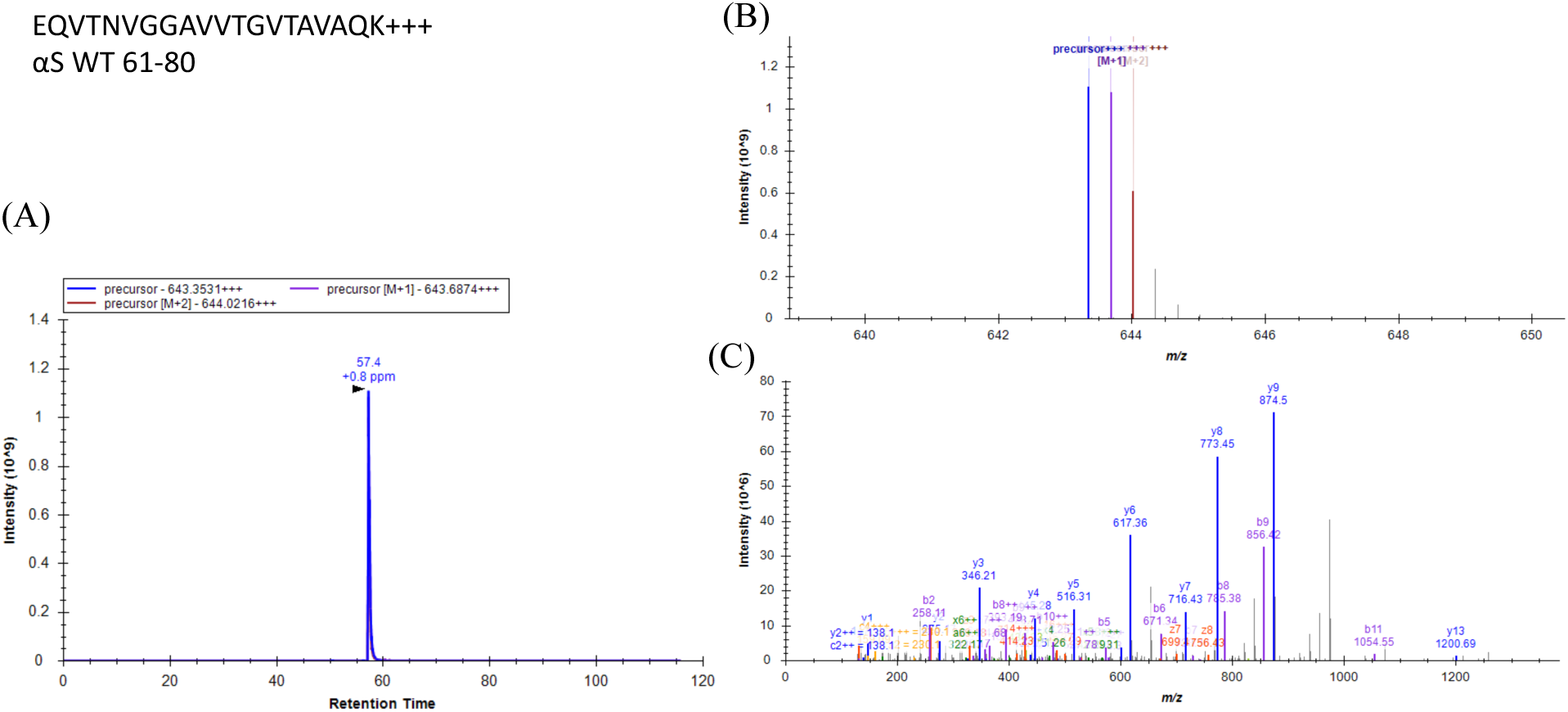
MS spectra of unmodified K_80_ peptide from αS WT standard. (A) Extracted ion chromatogram (EIC) (B) MS1 spectrum (C) MS2 spectrum with fragment annotation.

**Figure 52.**
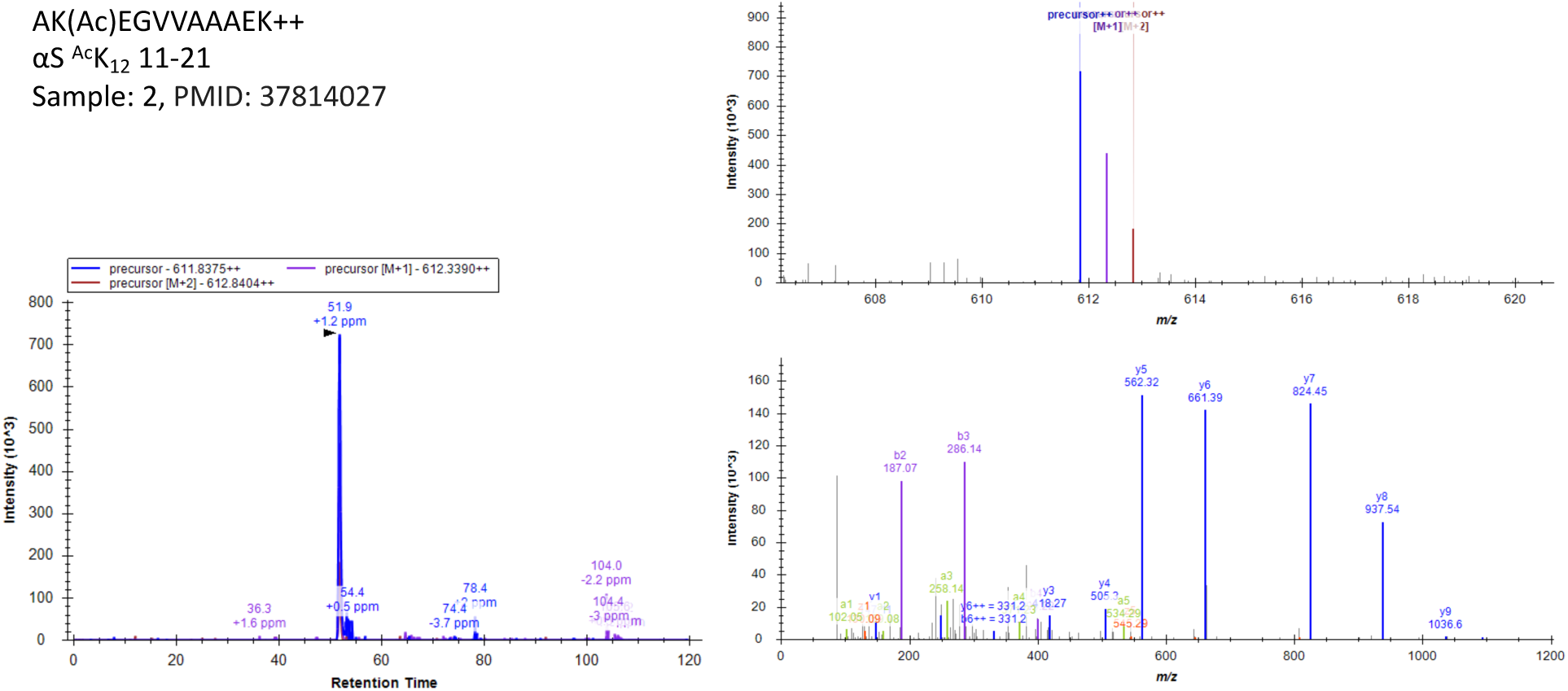
Representative MS spectra of ^Ac^K_12_ peptide from αS in patient sample 2. (A) Extracted ion chromatogram (EIC) (B) MS1 spectrum (C) MS2 spectrum with fragment annotation

**Figure 53.**
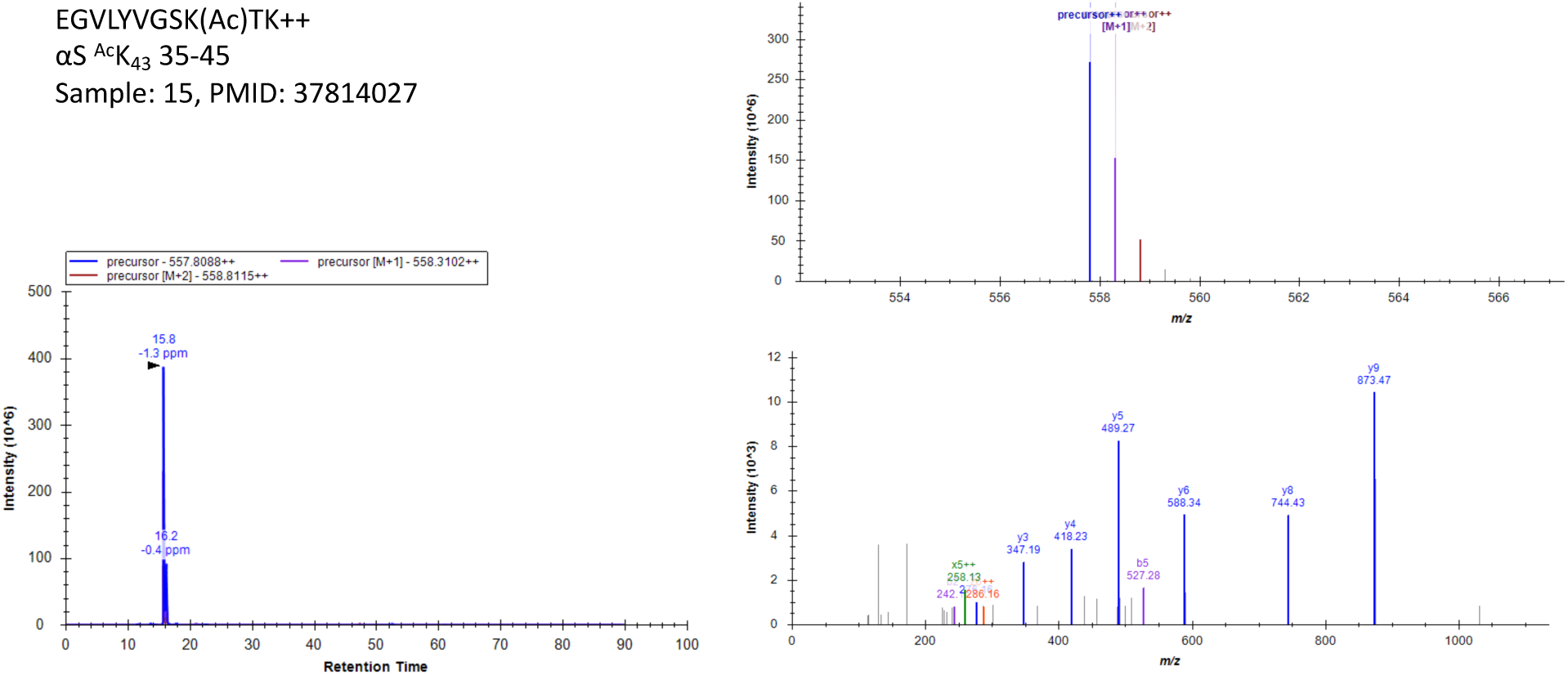
Representative MS spectra of ^Ac^K_43_ peptide from αS in patient sample 15. (A) Extracted ion chromatogram (EIC) (B) MS1 spectrum (C) MS2 spectrum with fragment annotation

**Figure 54.**
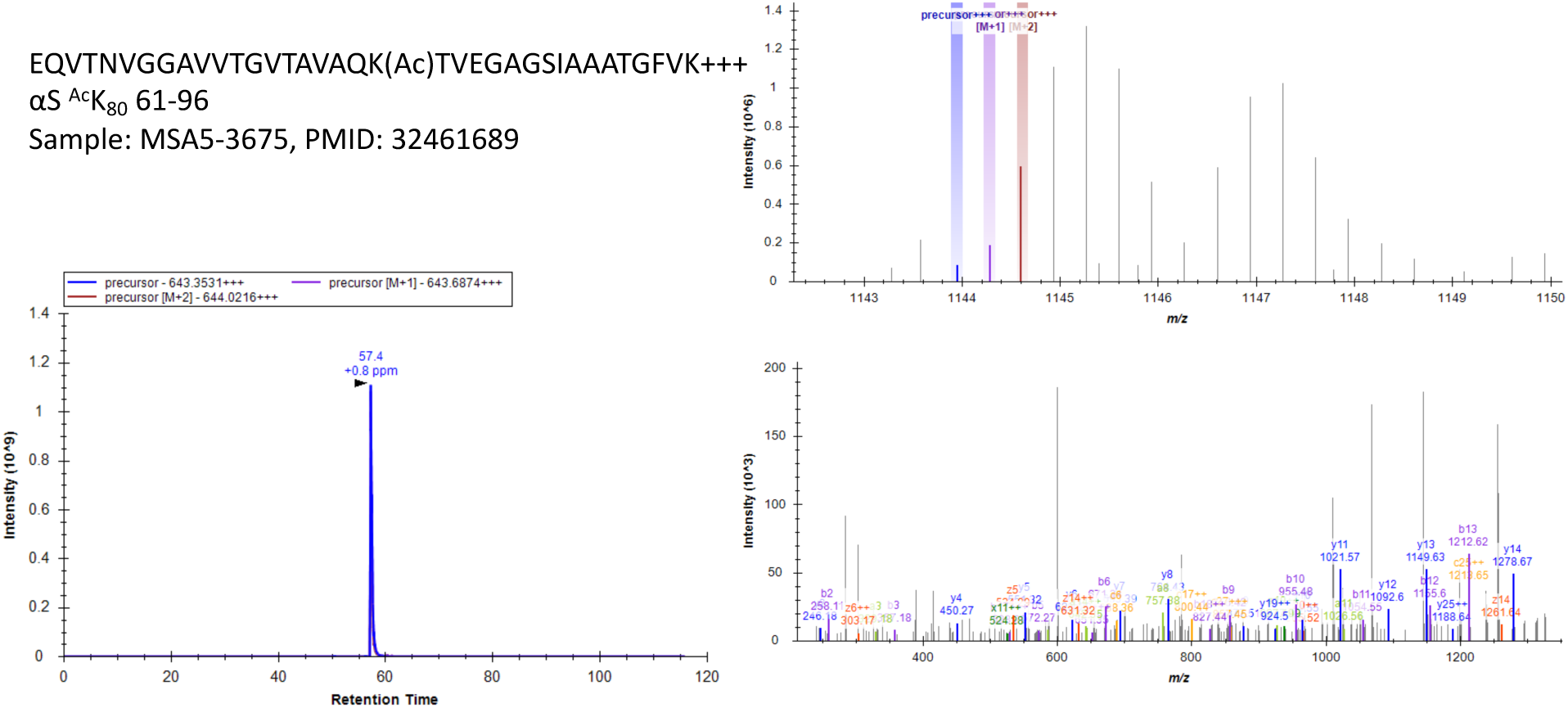
Representative MS spectra of ^Ac^K_80_ peptide from αS in patient sample MSA5-3675. (A) Extracted ion chromatogram (EIC) (B) MS1 spectrum (C) MS2 spectrum with fragment annotation

**Table 1.**
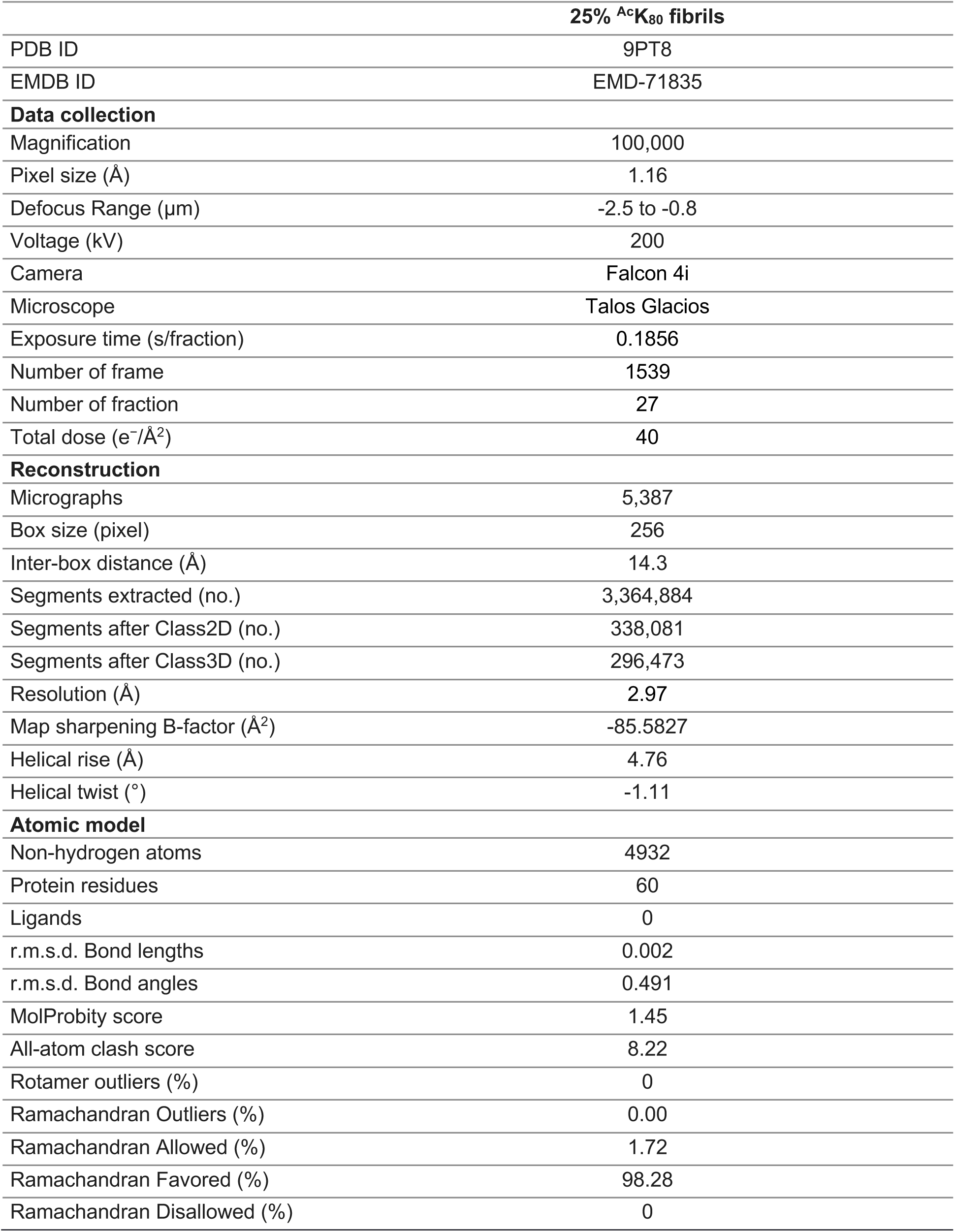
Statistics of cryo-EM data collection and refinement for 25% ^Ac^K_80_ fibrils.

**Table 2.** Statistics of cryo-EM data collection and refinement for WT and 100% ^Ac^K_80_ fibrils.

|  | **WT-A** | **WT-B** | **<sup>Ac</sup>K<sub>80</sub>-A** | **<sup>Ac</sup>K<sub>80</sub>-B** |
| --- | --- | --- | --- | --- |
| PDB ID | 9PTA | 9PT9 | 9PTC | 9PTB |
| EMDB ID | EMD-71837 | EMD-71836 | EMD-71839 | EMD-71838 |
| **Data collection** |  |  |  |  |
| Magnification | 100,000 | 100,000 | 100,000 | 100,000 |
| Pixel size (Å) | 1.16 | 1.16 | 1.16 | 1.16 |
| Defocus Range (µm) | -2.5 to -0.8 | -2.5 to -0.8 | -2.5 to -0.8 | -2.5 to -0.8 |
| Voltage (kV) | 200 | 200 | 200 | 200 |
| Camera | Falcon 4i | Falcon 4i | Falcon 4i | Falcon 4i |
| Microscope | Talos Glacios | Talos Glacios | Talos Glacios | Talos Glacios |
| Exposure time (s/fraction) | 0.1856 | 0.1856 | 0.1856 | 0.1856 |
| Number of frame | 1539 | 1539 | 1539 | 1539 |
| Number of fraction | 27 | 27 | 27 | 27 |
| Total dose (e<sup>-</sup>/Å<sup>2</sup>) | 40 | 40 | 40 | 40 |
| **Reconstruction** |  |  |  |  |
| Micrographs | 5001 | 5001 | 4038 | 4038 |
| Box size (pixel) | 256 | 256 | 256 | 256 |
| Inter-box distance (Å) | 14.3 | 14.3 | 14.3 | 14.3 |
| Segments extracted (no.) | 2,341,469 | 2,341,469 | 614,398 | 614,398 |
| Segments after Class3D (no.) | 696,830 | 1,317,407 | 102,778 | 200,042 |
| Resolution (Å) | 3.00 | 3.06 | 3.00 | 2.94 |
| Map sharpening B-factor (Å<sup>2</sup>) | -100.97 | -121.21 | -73.00 | -78.18 |
| Helical rise (Å) | -0.83 | 179.62 | -0.88 | 179.58 |
| Helical twist (°) | 4.71 | 2.36 | 4.76 | 2.38 |
| **Atomic model** |  |  |  |  |
| Non-hydrogen atoms | 4330 | 4150 | 4530 | 4190 |
| Protein residues | 630 | 620 | 650 | 610 |
| Ligands | 0 | 0 | 0 | 0 |
| r.m.s.d. Bond lengths | 0.005 | 0.003 | 0.002 | 0.002 |
| r.m.s.d. Bond angles | 0.497 | 0.542 | 0.418 | 0.469 |
| MolProbity score | 2.16 | 2.1 | 1.72 | 2.55 |
| All-atom clash score | 10.71 | 10.65 | 6.78 | 11.57 |
| Rotamer outliers (%) | 2.27 | 0 | 0 | 4.88 |
| Ramachandran Outliers (%) | 0 | 0 | 0 | 0 |
| Ramachandran Allowed (%) | 4.92 | 10.00 | 5.00 | 7.14 |
| Ramachandran Favored (%) | 95.08 | 90.00 | 95.00 | 92.86 |

**Table 3.** Quantification of acetylation %ratio by LC-MS/MS data analysis.

$\alpha S^{AcK_{12}}$
| Case no | Sample no | Pathological diagnosis | Clinical diagnosis | Soluble or insoluble | %ratio |
| --- | --- | --- | --- | --- | --- |
| 5 | 2 | PDD | PDD | Soluble | 10.33% |
| 5 | 7 | PDD | PDD | Soluble | 4.34% |
| 8 | 11 | AD | AD probable | Soluble | 1.79% |
| 9 | 16 | AD | CBS | Soluble | 0.09% |
| 10 | 19 | MSA | MSA-C | Soluble | 1.93% |
| 10 | 20 | MSA | MSA-C | Soluble | 1.91% |
| 13 | 29 | MSA | MSA-P | Soluble | 0.01% |
| 13 | 30 | MSA | MSA-P | Soluble | 3.85% |
| 13 | 31 | MSA | MSA-P | Soluble | 7.63% |
| 19 | 36 | Pathological aging | Normal | Soluble | 0.01% |
| 21 | 37 | Primary age-related tauopathy | Normal | Soluble | 29.29% |
| 21 | 38 | Primary age-related tauopathy | Normal | Soluble | 10.02% |
| 23 | 40 | Normal | Normal | Soluble | 8.62% |

$\alpha S^{AcK_{43}}$
| Case no | Sample no | Pathological diagnosis | Clinical diagnosis | Soluble or insoluble | %ratio |
| --- | --- | --- | --- | --- | --- |
| 9 | 15 | AD | CBS | Soluble | 38.81% |
| 9 | 16 | AD | CBS | Soluble | 0.09% |
| 9 | 17 | AD | CBS | Soluble | 10.44% |
| 9 | 18 | AD | CBS | Soluble | 0.00% |
| 13 | 28 | MSA | MSA-P | Soluble | 10.26% |
| 13 | 29 | MSA | MSA-P | Soluble | 0.00% |
| 14 | 31 | MSA | MSA-C | Soluble | 0.04% |
| 19 | 35 | Pathological aging | Normal | Soluble | 13.10% |
| 21 | 37 | Primary age-related tauopathy | Normal | Soluble | 0.02% |

$\alpha S^{AcK_{80}}$
| Case no | Sample no | Pathological diagnosis | Clinical diagnosis | Soluble or insoluble | %ratio |
| --- | --- | --- | --- | --- | --- |
| 3 | MSA3-9665 | MSA | MSA-C | Insoluble | 37.77% |
| 5 | MSA5-3675 | MSA | MSA-C | Insoluble | 41.77% |

**Table 4.**
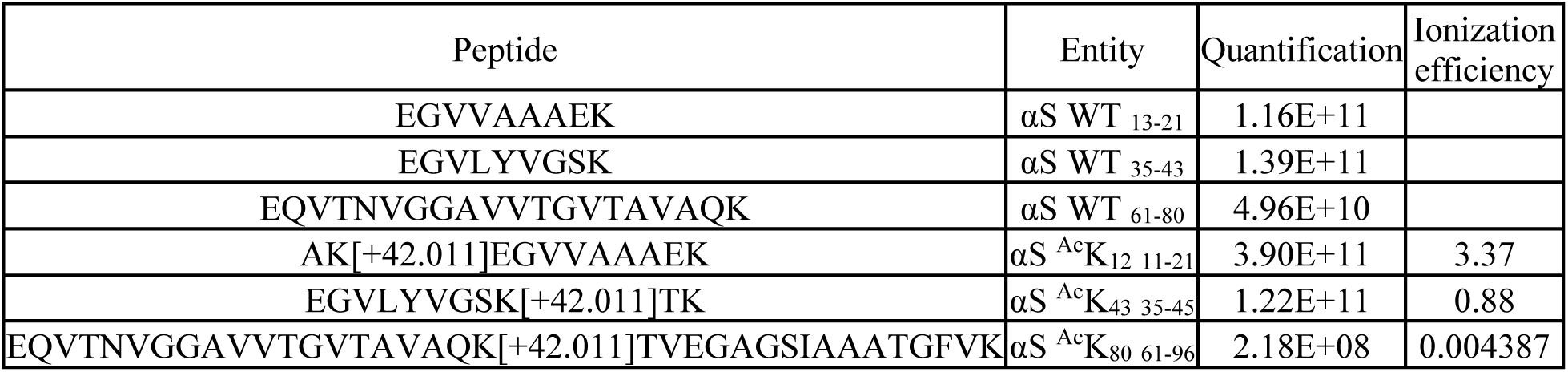
Ionization Efficiency of Acetylated Tryptic Peptides.

| Peptide | Entity | Quantification | Ionization efficiency |
| --- | --- | --- | --- |
| EGVVAAAEK | αS WT<sub>13-21</sub> | 1.16E+11 | |
| EGVLYVGSK | αS WT<sub>35-43</sub> | 1.39E+11 | |
| EQVTNVGGAVVTGVTAVAQK | αS WT<sub>61-80</sub> | 4.96E+10 | |
| AK[+42.011]EGVVAAAEK | αS<sup>Ac</sup>K<sub>12 11-21</sub> | 3.90E+11 | 3.37 |
| EGVLYVGSK[+42.011]TK | αS<sup>Ac</sup>K<sub>43 35-45</sub> | 1.22E+11 | 0.88 |
| EQVTNVGGAVVTGVTAVAQK[+42.011]TVEGAGSIAAATGFVK | αS<sup>Ac</sup>K<sub>80 61-96</sub> | 2.18E+08 | 0.004387 |

